# An Open, Evolving Brain and Cognitive Aging Simulator Enabled by Agentic Artificial Intelligence

**DOI:** 10.64898/2026.11.24.769018

**Authors:** Randy L. Buckner, Kristine B. Walhovd, Anders M. Fjell, Reisa A. Sperling, Deborah Blacker, Emily J. Rogalski, Bradley T. Hyman, Lars Nyberg

**Affiliations:** Department of Psychology and Center for Brain Science, Harvard University, Cambridge, MA, USA; Department of Psychiatry, Massachusetts General Hospital, Boston, MA, USA; Stanley Center of the Broad Institute of MIT and Harvard, Cambridge, MA, USA; Harvard Medical School, Boston, MA, USA; Center for Lifespan Changes in Brain and Cognition, Department of Psychology, University of Oslo, Oslo, Norway; Computational Radiology and Artificial Intelligence, Department of Radiology and Nuclear Medicine, Oslo University Hospital, Oslo, Norway; Department of Neurology, Mass General Brigham, Boston, MA, USA; Department of Epidemiology, Harvard T.H. Chan School of Public Health, Boston, MA, USA; Healthy Aging and Alzheimer’s Research Care (HAARC) Center, Department of Neurology, University of Chicago, Chicago, IL, USA; Department of Diagnostics and Intervention, Umeå University, Umeå, Sweden; Umeå Center for Functional Brain Imaging, Umeå University, Umeå, Sweden; Department of Medical and Translational Biology, Umeå University, Umeå, Sweden

## Abstract

An agentic AI simulator was built to advance our understanding about how the human brain and cognition change with age. The simulator generates worlds based on hypothesized age-linked cascades and then simulates studies across varied experimental designs and selection criteria. Each simulated world begins with 120,000 individual lives initiated by baseline parameters (e.g., genetic risk, birth year) and then changed by age-dependent cascades (e.g., amyloid-dependent, cardiovascular). Examples illustrate how the simulator predicts distinct data patterns in cross-sectional versus longitudinal designs, how alternative participant selection strategies emphasize baseline differences versus age-related decline, and how variable levels of association emerge between brain and cognition dependent upon design choices. Simulations of clinical trials provide information about design, power, and outcome considerations. Experiments on alternative simulated worlds evaluate how well hypothesized cascades accommodate real-world data patterns, capturing the insights and open puzzles inside the application. Most critically, the application is built to evolve. Its code contains a complete description of how to construct and update itself so that any person or AI agent can explore a new finding, recalibrate to emerging data, or build and simulate a novel future study. The simulator is offered as a tool for understanding data patterns, teaching, and generation of hypotheses to be tested in future studies using real-world data collection.

---

Within the field of brain and cognitive aging, a large and ever-growing corpus of observations has defined aging trajectories and begun to reveal the causes of their variability. Longitudinal tracking established differential vulnerability of certain brain regions to aging including some regions following non-linear trajectories (Raz et al., 2005). Hypotheses about how amyloid, tau and neurodegeneration unfold, combined with large-scale data collection efforts, led to models of ordered biomarker change that now organize research on Alzheimer’s disease (AD; Jack et al., 2010, 2013; Hansson, 2021). The observation that people with similar pathology differ in cognition inspired the concepts of reserve, resilience and resistance (Katzman et al., 1988; Stern, 2012; Arenaza-Urquijo and Vemuri, 2018; Stern et al., 2020), and theory and data support the idea that brain maintenance may underlie well-preserved cognition in aging (Nyberg et al., 2012). Dissociations between typical aging and neurodegeneration suggest that cognitive decline has multiple interacting causes (Buckner, 2004; Hedden and Gabrieli, 2004; Small et al., 2011).

As data accumulate across large numbers of studies, the challenge shifts from top-down theory testing to finding models that can accommodate the extant data. The question becomes less about whether a specific study confirms a hypothesis than whether any model can accommodate the growing body of findings, gathered in different samples with different measures and designs. Verbal theories rarely specify enough to say what each of those studies should have found (Guest and Martin, 2021; Yarkoni, 2022), and a literature can accumulate findings from specific datasets without establishing a cumulative framework that integrates all of them (Muthukrishna and Henrich, 2019).

Cognitive aging offers many examples of empirical complexities. Cross-sectional comparisons, influenced by cohort effects, indicate episodic memory decline from early adulthood, while longitudinal studies of the same abilities show little decline before about 60 (Rönnlund et al., 2005; Nilsson et al., 2009; Nyberg et al., 2020; see also Hofer and Sliwinski, 2001; Lindenberger et al., 2011; Raz and Lindenberger, 2011). Other cognitive functions, notably processing speed, display significant longitudinal decline before 60 (Schaie, 1994; Salthouse, 2009; Singh-Manoux et al., 2012). The association between AD pathology and dementia weakens in the oldest old (Savva et al., 2009), even as dementia incidence rises by some estimates (Corrada et al., 2010). Furthermore, the proportion of older adults counted as exhibiting high levels of episodic memory ranges from 2.9% to 43.4% across three Australian samples depending on the exact definition applied (Powell et al., 2024).

Moreover, available data are biased by the selection decisions and measurement processes built into any study. Research volunteers differ from the populations they come from (Falk et al., 2013; van Alten et al., 2024), and who is included in a sample impacts estimated brain trajectories, predictive accuracy, and prevalence (LeWinn et al., 2017; Benkarim et al., 2022; Hayes-Larson et al., 2022). Older participants are survivors, and thus longitudinal samples change as members die or leave a study (Lindenberger et al., 2002; Mayeda et al., 2016, 2018). Estimates of “normal” decline are inherently inflated by inclusion of preclinical dementia (Sliwinski et al., 1996), and repeated cognitive testing produces practice gains that can mask decline (Salthouse, 2010; Goldberg et al., 2015). Each process can be understood on its own and, with proper data and statistical approaches, can sometimes be corrected (e.g., Josefsson et al., 2012; Wang et al., 2026). Nonetheless, how they combine in a particular study is difficult to know (Banack et al., 2019), and the data patterns arising from their combinations are hard to intuit, especially when there are multiple differences between studies.

Simulation helps navigate these challenges by showing how complex and diverse data patterns can arise in real-world studies. Simulated cohorts have been used to quantify survival bias in studies of cognitive decline (Mayeda et al., 2016, 2018), show how retest gains and age-cohort differences confound estimates of within-person change (Hoffman et al., 2011), and ask whether selective survival could explain sex differences in dementia incidence (Shaw et al., 2021). In silico trials test interventions on virtual patients (Viceconti et al., 2016), and data-driven models place individuals along inferred disease timelines (Jedynak et al., 2012; Young et al., 2018) including within digital-twin frameworks (National Academies of Sciences, Engineering, and Medicine, 2024). Models are valuable well beyond prediction as they make assumptions explicit and expose ambiguities (Epstein, 2008; Smaldino, 2017).

The study of brain and cognitive aging can benefit from at least three distinct uses of a simulator. The first is to simulate the chain present in any given study, from how lives unfold in the real world to what the study might observe given study design processes including selection biases and measurement noise. The second is to simulate invented worlds with different combinations of causal cascades to test which best accounts for real-world findings. The third is to simulate data patterns for educational purposes allowing students to flexibly explore how patterns arise in existing studies and might arise in contemplated studies. In the past, a simulator with such capabilities would have required a sustained programming effort to build and modify (Knuth, 1984; Wilson et al., 2014). Large language models now offer an alternative. They write working code from natural-language descriptions (Chen et al., 2021) and can flexibly revise code without engaging expert programmers.

Here we describe a novel approach to simulation that takes advantage of the capabilities of agentic AI: an evolving, self-documenting simulator designed and built to connect the data patterns observed in real observational studies of brain aging to hypotheses about their underlying causes. The AgingBrain Simulator is a single HTML file that runs in a web browser. It was written by an AI agent through a conversation in which the first author, in dialogue with co-authors, specified the scientific components, the design and the checks. Critically, built into the HTML code are the history and guidelines about how the simulator is constructed so that it can evolve. A user instructs the AI agent to update the simulator to accommodate a new set of findings, explore an alternative causal mechanism, or contrast competing hypotheses about how an underlying mechanism might affect aging. The agent then constructs a revised world checked against reference values, rebuilds the simulator, captures insights and puzzles that emerge from the update, and saves it as a new version ready for another request or infusion of novel data. The simulator is thus a first version, not a finished model of brain aging. Its value lies in providing the field a shared, inspectable object that can be tested, criticized and extended as data accumulate.

## The Simulated World

The simulator builds a world of people who are each born with individual abilities and propensities for how their brain and cognitive faculties will change with aging. Within that world, each person’s life progresses as baseline determinants (e.g., genetic risk) interact with age-associated causal cascades (e.g., the AD cascade). The simulated individuals can then be studied using observational methods or experimentation through stages that mimic the full research cycle (Figure 1A). Specifically, the *World* stage defines and simulates the population that studies will draw from. The *Enrollment* stage sets who seeks to join the study, allowing socioeconomic status (SES), health, and source (e.g., memory clinic versus community) to affect volunteerism. The *Selection* stage applies the study’s criteria (e.g., only individuals without detected clinical dementia). The *Measurement* stage sets the error, validity, and practice effects associated with each measure, as well as the expected missed visits and dropout. The *Analysis* and *Follow-up* stages examine the study sample, and the *Intervention* stage can simulate the effects of a clinical trial. Each stage has access to the true simulated values, alongside the simulated results which reflect the selection decisions and application of the noisy measurement process (Figures 1C–1H). Finally, the *Imagine* stage is where a stage can be built that changes the application itself to flexibly explore and incorporate data in an entirely new manner.

**Figure 1.**
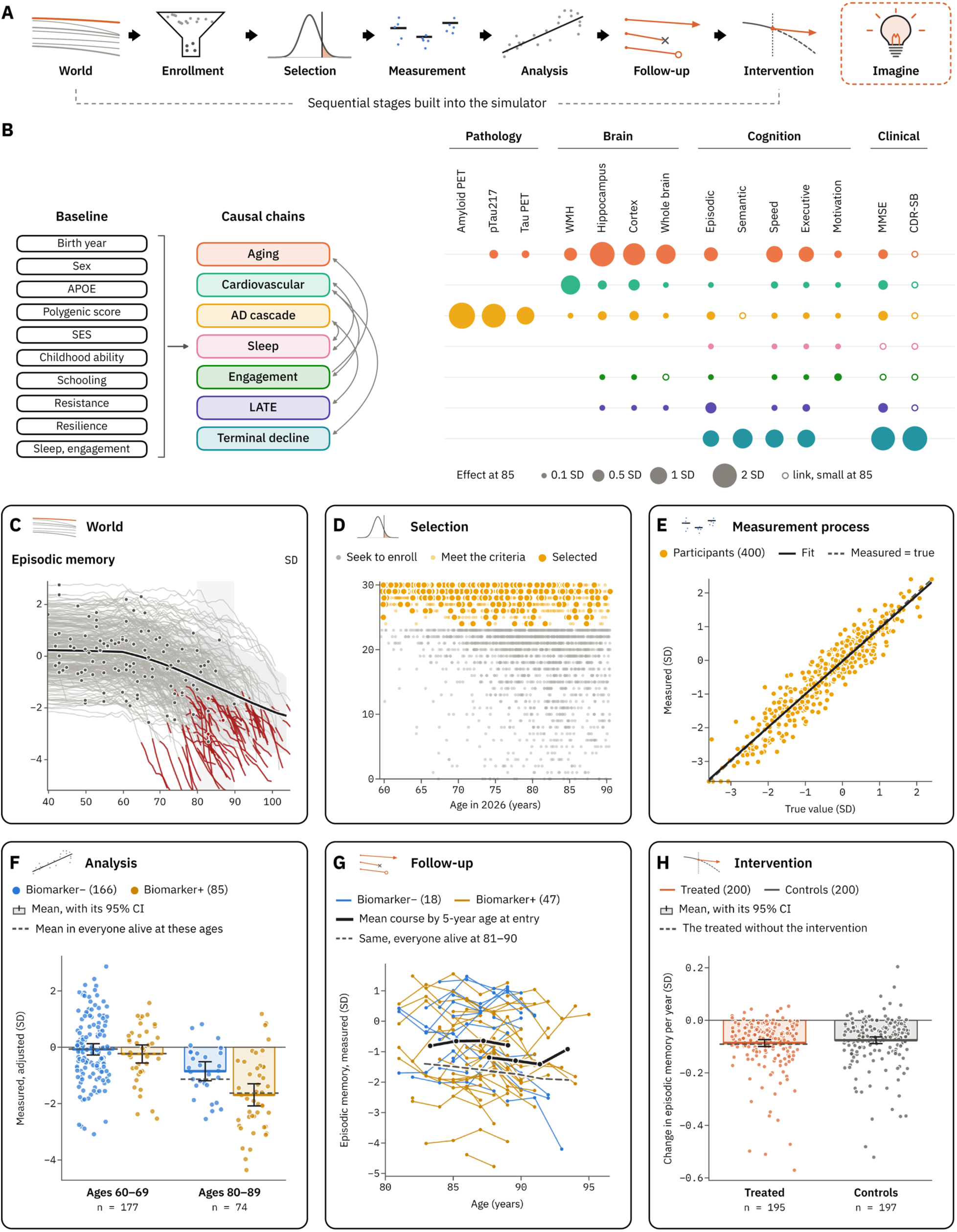
The simulator follows a study from the population in the world to findings observed in an individual experiment. (**A**) The seven stages from World to Intervention are shown. The Imagine stage (dashed) is reserved for a novel stage that can be created by the user (see text). (**B**) The model behind each life in the simulator is diagrammed. The baseline determinants set for each person (left) feed into causal chains (center). Arrows show links between chains. The effect of each chain on each phenotype at age 85 is visualized as each dot’s area (right; key shows 0.1–2 SD of 85-year-olds). Open circles mark links that exist but are small. (**C–H**) The application’s output, stage by stage, for an example study. (**C**) World stage: episodic memory from ages 40 to 100 is illustrated for 240 simulated people, each plotted as their own gray line (which turns red after a dementia diagnosis). The black line is the mean of the living individuals with the 10th–90th percentile band and each life’s value at its age in 2026 (dots). The shaded column highlights ages 80–89. (**D**) Selection stage: Mini-Mental State Examination (MMSE) score is plotted against age for those who seek to enroll (gray), those who meet the criteria (light amber), and those who are selected into the study (MMSE ≥ 24; dark amber). (**E**) Measurement process stage: measured values are plotted against true episodic memory. The fitted line (solid) lies near identity (dashed). (**F**) Analysis stage: in a second sample, measured episodic memory is plotted adjusted for sex and head size, in two age groups and split by biomarker status. Bars, means with 95% confidence intervals. Dashed, the means in everyone alive at those ages. (**G**) Follow-up stage: measured episodic memory is plotted over six years for the participants aged 81–90 at entry. Black, the mean course by 5-year age at entry. Dashed, the same for everyone alive at 81–90. (**H**) Intervention stage: change in true episodic memory per year is plotted after half the sample is randomized to slow the AD cascade by 50%, followed for 10 years. Bars, means with 95% confidence intervals. Dashed, the treated participants’ own lives without the intervention. The trial’s estimate brackets the true effect in these same participants.

The default world contains 120,000 lives, born between 1926 and 2008 and simulated year by year from age 18 to death (Figure 1B; Supplementary Figure 2B). Each life begins from baseline determinants including birth year, sex, genetic risks, childhood ability, and individual traits for resistance and resilience (Figure 1B; Supplementary Figure 2). A cohort module expresses these determinants through scaling that is affected by birth year (e.g., childhood mortality by age 18 decreases from the 1920s to the 1990s and can be set to interact with SES). For the simulation, each person’s levels are set at age 40, LATE and white matter lesions begin from that age, and amyloid from 50, where its risk is anchored to a population-based study (Roberts et al., 2018). Age 40 is chosen because of the availability of data to tune the simulator’s parameters in relation to advanced aging (e.g., the UK Biobank spans 40–69 years at baseline, Sudlow et al., 2015; the Alzheimer’s Disease Neuroimaging Initiative, ADNI, spans 55 to 90 years, Petersen et al., 2010).

Causal chains then change the brain and cognition including through aging^1^; a cardiovascular chain that gives rise to white matter lesions; an AD cascade in which amyloid precedes neocortical tau, and AD’s toll on the brain and cognition follows an ordered biomarker cascade (Jack et al., 2010, 2013, 2024; Supplementary Figure 3); sleep; engagement; limbic-predominant age-related TDP-43 encephalopathy, or LATE (Nelson et al., 2019); and terminal decline, a phase of steep cognitive loss just before death (Boyle et al., 2013). The chains interact and each variably affects phenotypes spanning pathology and biomarkers, brain structure, cognition and clinical ratings (Figure 1B, right).

In aging without the effects of a specific disease cascade, episodic memory and executive function decline linearly from age 60 (0.03 SD per year) and processing speed from age 25 (0.018 SD per year), with each decade of birth adding to the level reached (using a conservative cohort effect from the Betula study, cross-sectional differences of almost 1 SD between ages 35 and 60, against stable longitudinal scores; see Rönnlund et al., 2005; Rönnlund and Nilsson, 2008; Salthouse, 2009; Singh-Manoux et al., 2012; Nyberg et al., 2020). Part of the cohort gain comes through schooling, which raises the level reached rather than the subsequent rate of decline (Ritchie and Tucker-Drob, 2018; Lövdén et al., 2020; Nyberg et al., 2021). All three cognitive functions also decline faster late in life, by an amount that approximately doubles every 6 years, and each person’s cognition drifts on its own from 40 by amounts set to match how much of cognitive change is specific to episodic memory, speed and executive function (Tucker-Drob et al., 2019). About 30% of deaths after 40 come at the end of a terminal phase, which is set to last 3 years (2.4 on average, because some people die suddenly before it ends) and during which cognitive domains fall by a mean of 0.45 SD a year, at a pace that varies widely between people, mirroring how cognitive decline speeds up in the last years of life in clinical-pathologic cohorts (Boyle et al., 2013; Wilson et al., 2020). In those cohorts, terminal decline accounts for about 70% of late-life cognitive loss (Wilson et al., 2020). Because the risk of entering a terminal phase follows each person’s own risk of death, most terminal phases begin late in life. Dementia requires both a low level and a fall: global cognition at or below −2.3 SD after a decline of at least 1.2 SD from 40. Once it begins, it is staged by the Clinical Dementia Rating (CDR), whose mild, moderate and severe stages parallel stages 4–6 of the clinical staging in the revised Alzheimer’s Association criteria (Jack et al., 2024), and the risk of death rises with the stage (Andersen et al., 2010).

## Comparing Selection Rules in the Same Simulated Lives Reveals Cohort Structure

Because every life is simulated, any individual can be followed and explained. This aspect of the simulator is especially useful for understanding how different selection strategies based on age, absolute and changing cognitive levels, and clinical criteria relate to one another and how their results can be interpreted. Figure 2A shows five simulated women at 85 selected by different criteria. The first is the median woman at 85. The second has mild cognitive impairment (MCI) at 85, which she reached at 82. A third is a SuperAger at 85 by the selection rule of Harrison et al. (2012) and Rogalski et al. (2013). A fourth is a Josefsson maintainer, by the rule of Josefsson et al. (2012), which is one implementation of the concept of brain and memory maintenance developed in Nyberg et al. (2012). And the fifth, a slope maintainer, defined as someone who loses no more than 0.3 SD of memory from 75 to 85, reflects an alternative selection strategy also directed toward the concept of maintenance. Slope maintenance is intended to identify successful aging by lack of change (e.g., Willis and Schaie, 2005; Persson et al., 2006). These selection strategies thus include rules developed across distinct lineages to identify successful aging. Of the women who reach 85 in the simulation, 19.9% are SuperAgers, 14.0% are Josefsson maintainers, and 18.1% are slope maintainers. To visualize the causes of episodic memory performance, Figure 2B shows each simulated individual referenced to their episodic memory level.

**Figure 2.**
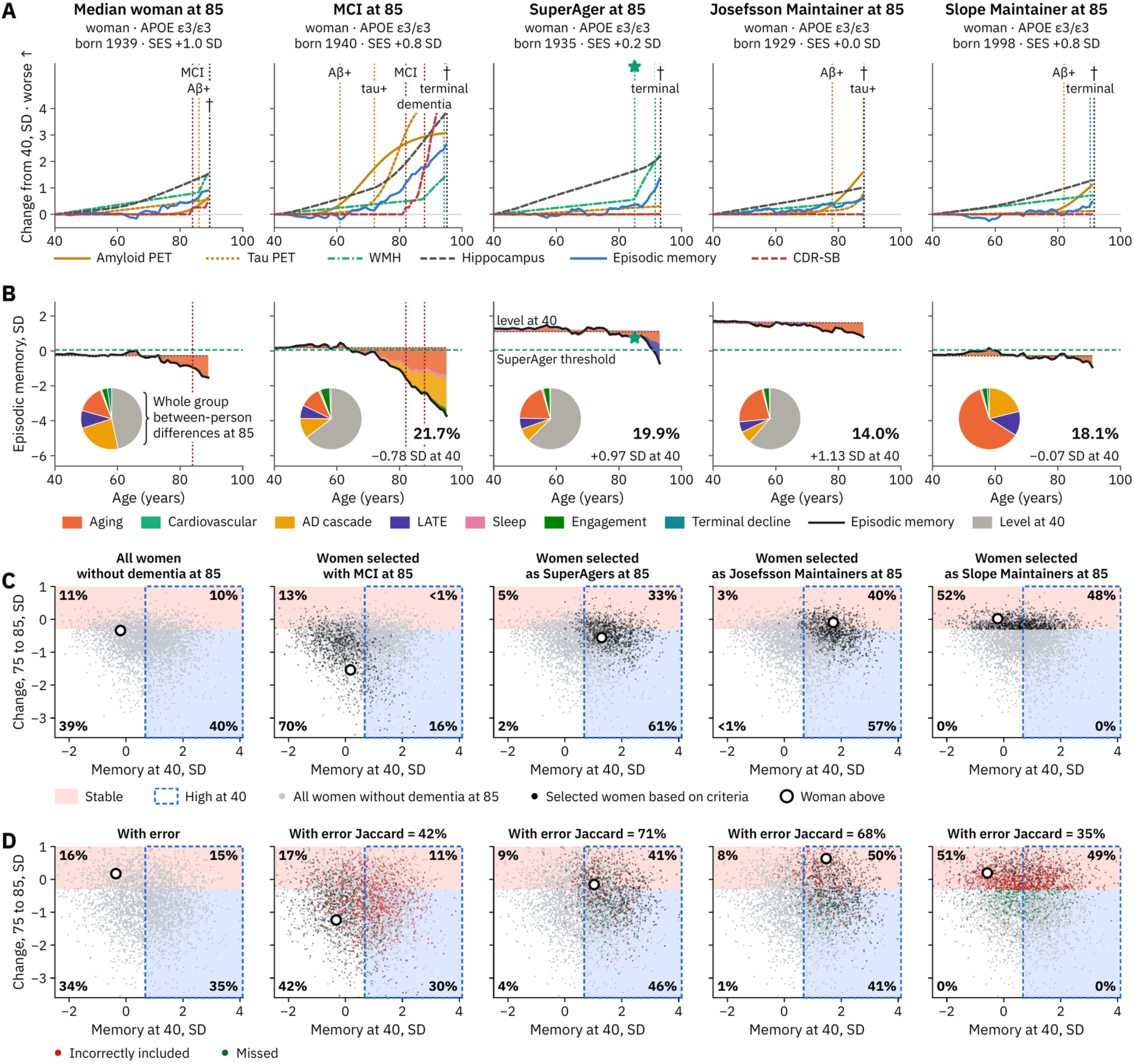
The simulated world illustrates how selection criteria impact data patterns and interpretation. (**A**) Five examples of simulated women at age 85 are plotted as change from the person’s own level at 40. Each line within the plot shows the value of a measured phenotype for that person in SD of 85-year-olds (e.g., amyloid PET is shown in orange). Worse is plotted upwards. Dotted vertical lines mark amyloid positivity (Aβ+), tau positivity, mild cognitive impairment (MCI; red), dementia (red), the start of a terminal phase (teal) and death (†). First column, the median woman at 85. Of the women who reach 85, this person is at the median of episodic memory. She has MCI from 84, becomes amyloid positive at 86 but never tau positive, and dies at 89. Second column, a woman with MCI at 85 (she converted at 82). Third column, a SuperAger at 85 by the rule of Harrison et al. (2012) and Rogalski et al. (2013) (star). Fourth column, a maintainer at 85, by the rule of Josefsson et al. (2012). Fifth column, a slope maintainer at 85. (**B**) Each life’s episodic memory (black) split into its level at 40 (dotted horizontal) plus the contribution of each causal chain (colored areas that fill inside the line with losses plotted below and gains above). The dashed green line shows the SuperAger threshold for episodic memory. Each group’s pie chart shows what sets the selected women apart from the average dementia-free woman, splitting the pie into their (baseline) level at 40 (gray wedge) and each causal chain (colored wedges). For the selected groups, the contributions are calculated relative to the mean performance of dementia-free women; for the Median woman, the contributions reflect variability across all simulated women, including those with dementia. The bottom right percentage is the share of women reaching 85 who meet each criterion, and the value below it is the selected group’s mean level at 40 (in SD relative to all dementia-free women). (**C**) Each woman’s true episodic memory at 40 (level) is plotted against her true change from 75 to 85 (slope). Gray dots are all women without dementia at 85. Black dots are the subset of selected women. For reference, the open circle shows the woman from the A and B panels above. The stable memory zone is highlighted in pink (reflecting a loss of no more than 0.3 SD from 75 to 85). The dashed blue box shows individuals whose memory at 40 is above the median; the shaded blue zones indicate individuals whose memory declined. The numbers in the corners give the share of each group in each quadrant. Note how the multiple successful aging criteria select overlapping but distinct groups of individuals (three right columns). The SuperAger and Josefsson Maintainer rules select women who mostly start high (and often decline from 75 to 85) while the Slope Maintainer rule selects stable women, from below and above the median at 40. (**D**) The same women and criteria as in C, but with each measured value including error. Black dots are women selected on measured values who meet the criterion on true values (i.e., correctly included). Red dots are incorrectly included, and dark green dots are misses. Each title gives the Jaccard index, which would be 100% if measurement error changed no one’s selection (the first panel has none). Measurement error blurs every boundary.

Aggregating all 85-year-olds by these criteria provides information about what the definitions capture (Figure 2B, pies). Compared with women free of dementia at 85, SuperAgers stand out mainly because they start higher, and so do many Josefsson maintainers. Slope maintainers are selected without regard to baseline level primarily because they lose less to aging, the AD cascade and LATE. Figure 2C makes these contrasts explicit by plotting every woman in the selected group by her memory at 40 (level) against its change from 75 to 85 (slope). The SuperAger and Josefsson rules select women who mostly start high (94% and 97% of each group) and often decline from 75 to 85 (63% and 57%). A large portion lost less than 0.3 SD of memory, many more than would be expected by chance, indicating these two selection rules enrich for maintained memory. The slope rule, by contrast, directly selects only women whose memory holds steady, roughly equally from below and above the median level (52% below and 48% above). Inclusion of measurement error reveals additional features of the selection rules (Figure 2D). The slope rule is strongly and disproportionately affected by measurement error, with a Jaccard index (35%) roughly half that of the SuperAger (71%) and Josefsson (68%) rules. This is because a change between two tests carries the error of both tests. The reliability of measured change in memory in the present simulation is about 0.5, in contrast to the considerably higher reliability of 0.9 for between-person level differences. Thus, complex tradeoffs exist between selection strategies that can profoundly affect who is analyzed within a specific study.

The aggregate behavior of the world can be set against what has been measured in real populations (Figure 3). Individual trajectories fan out with age (Figure 3A), and the average course of each phenotype can be followed separately in people who never develop dementia and in people whose dementia begins at 80 (Figure 3B). Survival, amyloid positivity and dementia can be estimated across age and compared to published estimates (Figure 3C). The world also shows what drives differences between people at different ages. Through age 55, the level at 40 accounts for more than 95% of the between-person variance in episodic memory, and for 92% at 60. After that, what has happened during adult life counts for progressively more. By 80 the level at 40 accounts for 56%, the AD cascade for 17% and aging, which includes each person’s own drift, for 14% (Figure 3D; see Grødem et al., 2026 for recent data consistent with these estimates).

**Figure 3.**
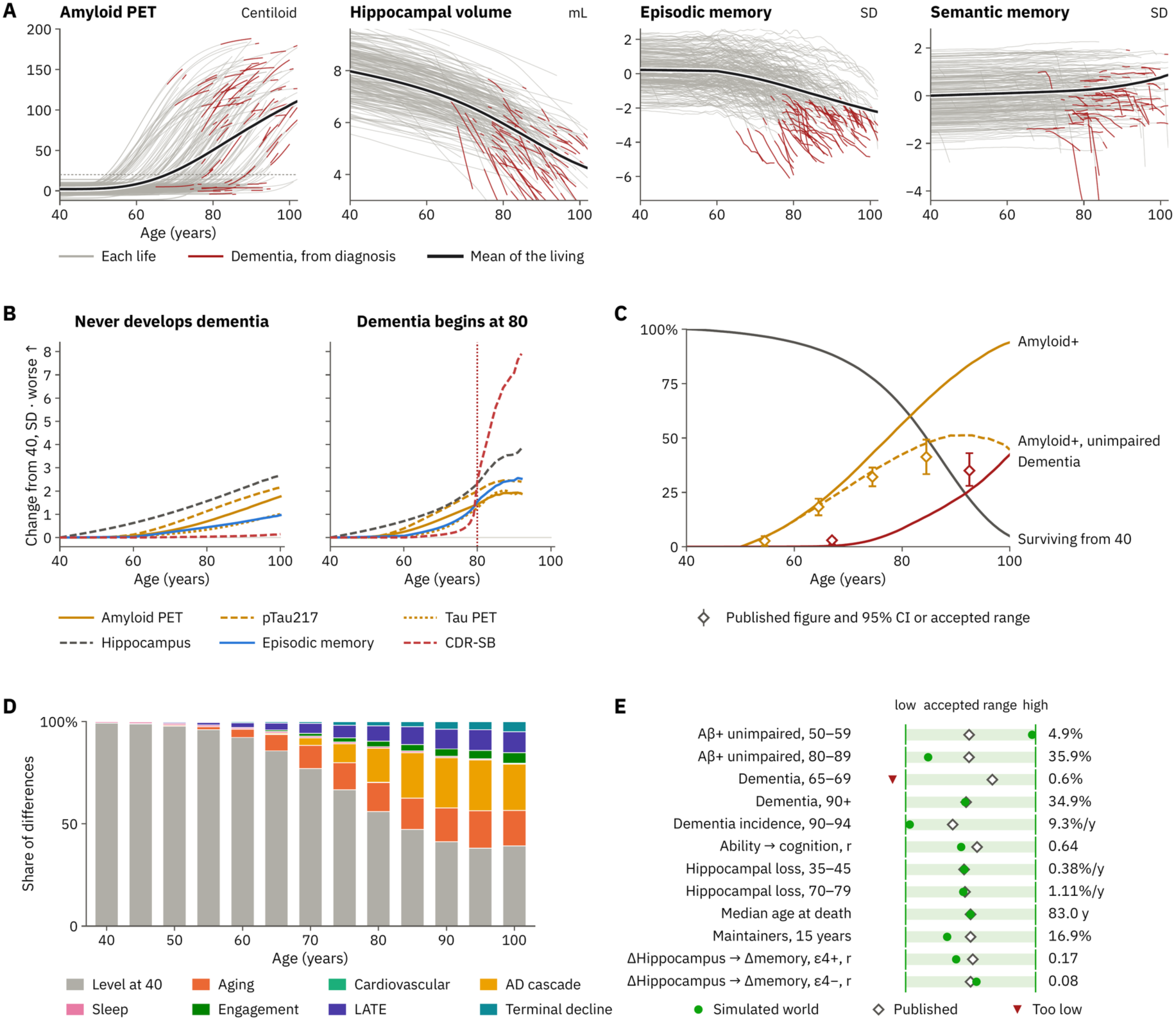
The simulated world reproduces population-level values measured in different real-world studies. (**A**) A subset of the phenotype values for 240 of the 120,000 lives is illustrated from age 40 until death for amyloid PET (Centiloid; fine dashed line, positivity at 20), raw bilateral hippocampal volume (mL), episodic memory (SD) and semantic memory (SD). Each gray line is an individual person; the line turns red after a dementia diagnosis. Black, the mean of everyone alive at each age. (**B**) Two average lives: the mean of each phenotype among the 78,451 lives that never develop dementia (left) and the 1,467 whose dementia begins at 80 (right, dotted red vertical line marks dementia). Worse is upward. (**C**) By age: survival from 40, amyloid positivity among the living, amyloid-positive and cognitively unimpaired people as a share of those without dementia, and dementia among the living, with published estimates (diamonds) and their 95% confidence intervals (bars; from Roberts et al., 2018, and Manly et al., 2022). (**D**) The share of between-person differences in episodic memory due to the level at 40 and to each chain, among the living every 5 years (these breakdowns are the same as the pie charts in Figure 2B but include everyone alive at that age). (**E**) The twelve reality checks, each scaled to its accepted range (0, low end; 1, high end). Green dots plot the values of the simulated world that are in range. The red triangle indicates a failed check. One simulated world, built from assumptions about individual lives and calibrated against these checks, produces values within the accepted range for 11 of these 12 population values.

Reality checks compare the simulated world with published reference values (Table 1): cognitively unimpaired people who are amyloid positive (Roberts et al., 2018); dementia prevalence at 65–69 and at 90 and older (Manly et al., 2022); dementia incidence at 90–94 (Corrada et al., 2010); the correlation of childhood ability with late-life cognition (Gow et al., 2011); hippocampal atrophy in studies whose participants averaged under 55 and 70 or older (Fraser et al., 2015); median age at death (Social Security Administration, 2026); the share of memory maintainers over 15 years (Josefsson et al., 2012; see also Josefsson et al., 2023); and how closely hippocampal and memory change go together in APOE ε4 carriers and in non-carriers (Gorbach et al., 2020).

**Table 1.** Reality checks compare the simulated world with published values. Each row gives a check (as plotted in Figure 3E), its value in the default world (version 104; world seed 7, 120,000 lives, study year 2026), the published value, the target range within which the application counts the check as passed, and the source. Methods (Reality checks) describe how each check is computed. Because the defaults were calibrated against these values, passing 11 of the 12 checks shows that one set of assumptions can accommodate nearly all of them together, not that the assumptions are correct. Note that some checks evolved across simulator versions. For example, a check for the expected share of SuperAgers was included in version 88 (Figure 6) that was later removed because expected values are still emerging.

| Check | Simulated | Published | Target range | Source |
|---|---|---|---|---|
| Amyloid positive and cognitively unimpaired, ages 50–59 | 4.9% | 2.7% (95% CI 0.5–4.9%) | 0.5–5% | Roberts et al. (2018): Mayo Clinic Study of Aging, amyloid PET in a population-based sample without dementia |
| Amyloid positive and cognitively unimpaired, ages 80–89 | 35.9% | 41.3% (95% CI 33.4–49.2%) | 33–50% | Roberts et al. (2018) |
| Dementia prevalence, ages 65–69 | 0.6% (below range) | 3% | 1–4% | Manly et al. (2022): Health and Retirement Study, Harmonized Cognitive Assessment Protocol; a nationally representative US sample, 2016 |
| Dementia prevalence, ages 90 and older | 34.9% | 35% | 28–43% | Manly et al. (2022) |
| New dementia per year, ages 90–94 | 9.3% | 12.7 per 100 person-years (11.9% a year once converted) | 9–17% | Corrada et al. (2010): The 90+ Study |
| Early-life ability and global cognition at 77–81, r | 0.64 | 0.66 (same test at 11 and 79) | 0.55–0.75 | Gow et al. (2011): Lothian Birth Cohort 1921 |
| Hippocampal volume loss per year, ages 35–45 | 0.38% | 0.38% (studies averaging age 40) | 0.2–0.6% | Fraser et al. (2015): meta-analysis of longitudinal hippocampal atrophy in healthy aging |
| Hippocampal volume loss per year, ages 70–79 | 1.11% | 1.12% (studies averaging age 74) | 0.8–1.5% | Fraser et al. (2015) |
| Median age at death, lives reaching 40 | 83.0 years | ≈83 years (women 85, men 81) | 80–86 years | Social Security Administration (2026): US period life table, 2023 |
| Memory maintainers over 15 years, relative to age peers | 16.9% | 18% (attrition-corrected) | 15–21% | Josefsson et al. (2012): the Betula study |
| Hippocampal and episodic memory change, APOE ε4 carriers, r | 0.17 | 0.21 | 0.06–0.35 | Gorbach et al. (2020): Lifebrain consortium, non-demented adults measured twice |
| Hippocampal and episodic memory change, non-carriers, r | 0.08 | 0.07 | −0.02 to 0.16 | Gorbach et al. (2020) |

In the default world, eleven of these twelve checks fall within their accepted ranges (Figure 3E). The exception is dementia prevalence at 65–69 (0.6% against a published 3%, 95% CI 1–4%; Manly et al., 2022). In the model, few people become amyloid positive early enough for the AD cascade to cause dementia by these ages, and the model has no frontotemporal or other early-onset dementias. Because the defaults were calibrated against these checks, passing eleven of them shows that one set of assumptions can accommodate nearly all of these values together. The calibration is tuned to the default world; the values therefore do not validate the assumptions. Rather, they indicate one set of assumptions can accommodate a wide range of real-world data patterns. The ranges and sources, with notes on several checks, are written into the application, and the checks are recomputed whenever a setting changes. Users may wish to calibrate against alternative check values or construct entirely novel checks.

## Familiar Patterns and Novel Predictions from Study Decisions

The simulated world can be interrogated by changing experimental decisions and examining how resultant data patterns compare to real-world experimental data (Figure 4). The results are not written into the simulator; each result arises when programmed study processes, such as selection rules, measurement error and practice effects, act on the simulated lives.

**Figure 4.**
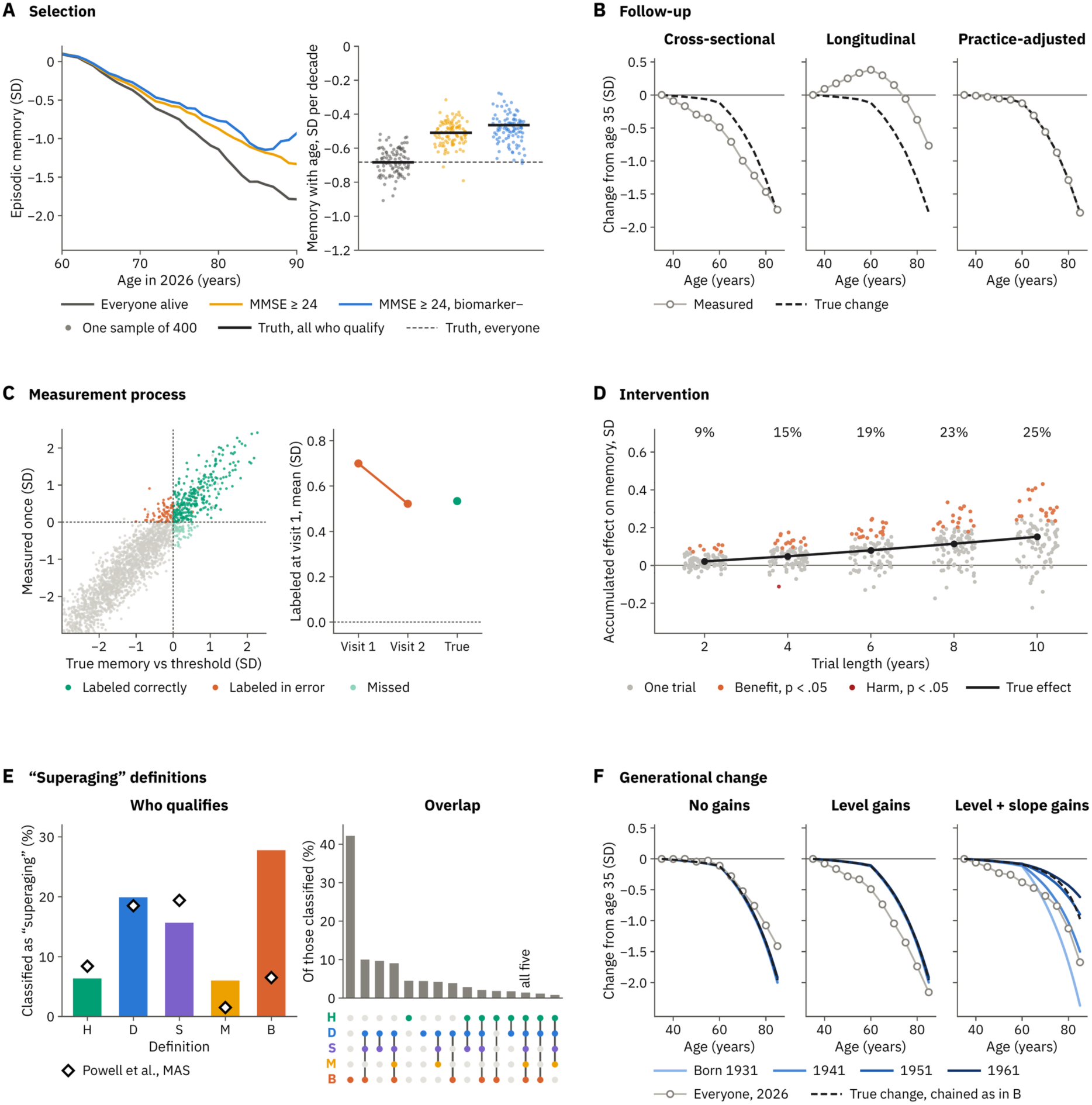
Examples of data pattern predictions that emerge from study and analysis decisions. (**A**) Selection. Left, mean episodic memory is plotted by age among everyone alive aged 60–90 (gray), the subset with MMSE ≥ 24 (amber), and the subset who are also free of amyloid and tau at a strict cutoff (blue). Right, the cross-sectional slope of measured memory on age in each of 100 samples (each N=400) plotted for each criterion (dots). The black bar is the slope among all who qualify; the dashed line is the reference from everyone alive. (**B**) Follow-up illustrates differences between cross-sectional and longitudinal estimates of episodic memory decline, reflecting the cohort and practice effects built into the model (Rönnlund et al., 2005; Nyberg et al., 2012, 2020). Everyone aged 33–87 with MMSE ≥ 24 is measured and, if alive, measured again five years later. The measured estimates are shown in circles. Left, cross-sectional, each 5-year age group’s mean memory at entry relative to the youngest group; middle and right, longitudinal, the five-year changes of successive age groups chained from 35, with practice effects (middle) and with the practice gain removed (right). The true within-person change is shown by the dashed line which overlaps the practice-adjusted line. (**C**) Measurement process. Left, measurements are plotted against true episodic memory, each relative to the SuperAger memory threshold, for half of the dementia-free people aged 80–89 in 2026. Correctly labeled measures are shown in green, erroneously labeled in orange, and missed in light green. Gray dots are the remaining people. Right, among the 712 labeled at one test (Visit 1), mean memory relative to the threshold at that first test and at a second test with new error (Visit 2), and the true value for reference. Note the regression to the mean. (**D**) Intervention. One hundred simulated A4-like trials in amyloid-positive (at A4’s approximated cutoff, see Methods), cognitively unimpaired adults aged 65–85, half randomized to a therapy that clears 30% of plaque (see Methods), each analyzed after 2, 4, 6, 8 and 10 years. Dots, each trial’s estimated effect on memory, accumulated over the trial (mean slope, treated minus control, times trial length); orange, significant benefit (p < 0.05); red, significant harm; black, the true effect, the mean difference between each participant’s life with and without the intervention; percentages, power to detect the true effect. (**E**) Varied “superaging” definitions. Five published definitions that compare older with younger adults are plotted from Powell et al. (2024). They refer to: H, Harrison et al. (2012) and Rogalski et al. (2013); D, Dang et al. (2019); S, Sun et al. (2016); M, Maccora et al. (2021); B, Bott et al. (2017). Each was applied to the dementia-free people aged 70–90 with MMSE ≥ 24. The bars plot who qualifies from the simulated values, and the diamonds plot the values from the Sydney Memory and Ageing Study (MAS) published in Powell et al. (2024). Overlap plots the share of people classified by at least one definition with where the overlap comes from shown below (dots). Combinations with fewer than 45 people are not included. (**F**) Generational change. The same simulated people from three worlds are plotted that differ only in their generational settings: no gains; level gains (the default world); and with level and slope gains (later-born people decline more slowly). The measured estimates are shown in circles. The true changes in episodic memory from age 35 of people born in 1931, 1941, 1951 and 1961 are shown as separate-colored lines. The dashed line, as in B, plots the true change chained across the age groups of 2026.

For example, selection dramatically changes the answer to the simple question: how much does memory decline with advancing age? In 100 random samples of 400 people aged 60–90, the cross-sectional decline in measured episodic memory averaged 0.68 SD per decade, close to the 0.68 SD per decade among everyone alive at those ages (Figure 4A). Excluding people with an MMSE score (Folstein et al., 1975) below 24, a common criterion in studies of normal aging, cut the estimate by about a quarter, to 0.51 SD per decade. Excluding individuals with amyloid or tau at strict cutoffs further lowered it to 0.49. This set of simulations illustrates that what a study reports as normal aging depends on its definition, and contamination by preclinical dementia can impact estimates (Sliwinski et al., 1996).

Selection also changes how strongly two variables appear to relate, even when every sample is drawn from the same world (Figure 5; see Vidal-Piñeiro et al., 2025 for context). Among the 3,496 people aged 80–84 in 2026, hippocampal volume and episodic memory correlate at *r* = 0.46 once sex is regressed from both and head size from hippocampal volume. A random sample of 400 recovers that association (*r* = 0.43), but samples of 400 drawn with four of the simulator’s other strategies weaken it: *r* = 0.31 among nondemented people, 0.28 among those also free of amyloid and tau measured via biomarkers, 0.14 among people with MCI and 0.08 among SuperAgers. Across 100 samples per strategy and at every age from 70 to 89, each strategy’s typical association falls below that of a random sample (Figure 5F), and selecting people for their memory, as SuperAger studies do, largely removes its association with brain structure. A weak association between brain and memory in a selected sample can therefore reflect who was enrolled rather than a weak link between them. Note also that the absolute hippocampal volume levels still reflect the selection processes (e.g., compare the bottom triangles in Figure 5D to 5E).

**Figure 5.**
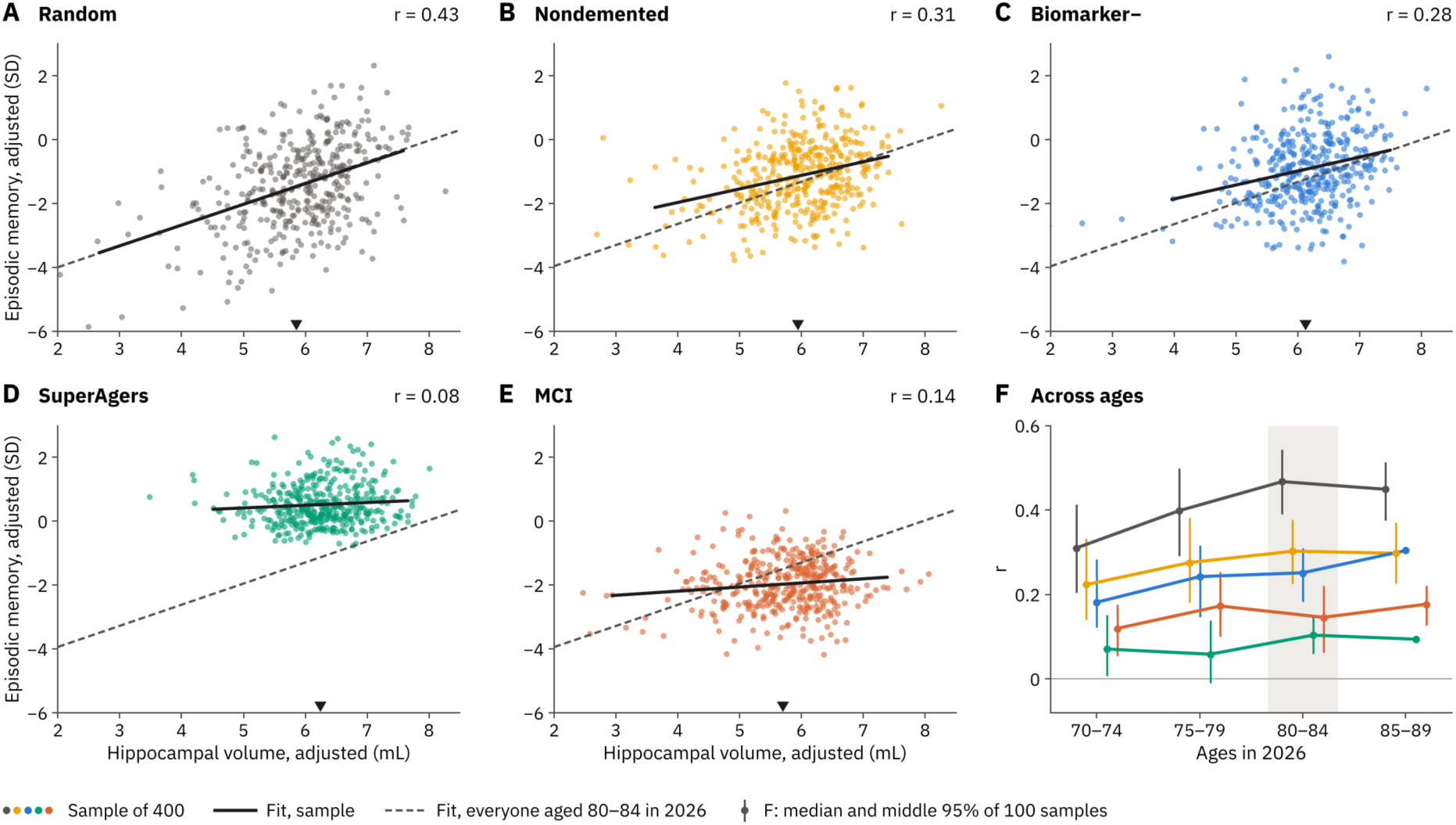
Study enrollment decisions change how strongly hippocampal volume tracks episodic memory. Each scatter plot shows one sample of 400 people aged 80–84 in 2026 using one Selection strategy and measured with error. Both variables are adjusted for sex, and hippocampal volume is additionally adjusted for total intracranial volume (head size). *r* is the correlation between the adjusted values. The sample’s best fit is plotted in black; dashed, as a reference, plots the best fit for everyone alive at 80–84 (*r* = 0.46). Triangles at the bottom show the sample’s mean adjusted hippocampal volume. (**A**) Random: *r* = 0.43. (**B**) Nondemented, MMSE ≥ 24: *r* = 0.31. (**C**) Biomarker negative, nondemented and free of amyloid and tau at strict cutoffs: *r* = 0.28. (**D**) SuperAger: *r* = 0.08. (**E**) MCI: *r* = 0.14. (**F**) The median *r* of 100 samples of 400 for each strategy at four ages in 2026, colored as in A–E. Bars span the middle 95% of samples, and shading marks 80–84. At 85–89, each SuperAger and biomarker-negative sample uses the same simulated people. Each rule that selects a subset of subjects weakens the association. Choosing people for memory attenuates the association. Note that a remnant of the selection process is expressed in the mean hippocampal volume, which is greater in SuperAgers than in those with MCI or in a random sample.

As another example, how change is expressed shapes what is captured and visualized. Analyzed as in the Betula study (Rönnlund et al., 2005; Nyberg et al., 2012, 2020), everyone in the simulated world aged 33–87 with MMSE scores of 24 or more was measured once and again five years later, and the five-year changes of successive age groups were chained from age 35 (Figure 4B). Cross-sectionally, memory fell steadily from 35, by 0.49 SD at 60 and 1.47 SD at 80, although within-person memory changed little before 60 (0.12 SD by 60). The earlier cross-sectional fall reflects cohort gains, with later-born people starting higher (Rönnlund and Nilsson, 2008). Longitudinally, memory appeared to rise until 60 because practice at the second test more than offset decline in many groups, and by 85 it appeared to have fallen 0.77 SD, less than half the true 1.78 SD. With the practice gain removed, the longitudinal estimate matches the true change as expected given the assumptions of the model – flat until about 60 and declining thereafter, and by 85 it slightly exceeded the cross-sectional estimate (1.74 SD), because those who decline leave the sample as they develop dementia or die. The simulated curves resemble Betula’s (Nilsson et al., 2009; see also Schaie, 2009; Salthouse, 2009, 2019). Because the values in the simulation are known, the simulation shows how cross-sectional plots can distort intuitions. Longitudinal change with the practice gain removed reflects the true world values in this model that assumes consistent practice effects across the sample (Figure 4B, right). Other model assumptions, such as variable practice effects between people, will lead to other distortions and multiple variations can be simulated to gain intuitions.

As a third example, the simulation illustrates how single measurements can misclassify individuals. Of 4,587 dementia-free people aged 80–89 in 2026, 713 (16%) met the rule for SuperAgers (Harrison et al., 2012; Rogalski et al., 2013). One assessment with typical error labeled 712 people as SuperAgers, of whom 81% met the rule, and a second, equally good assessment agreed for only 72% of them (Figure 4C). The labeled group’s mean memory fell from 0.70 to 0.52 SD above the threshold on retest, close to its true value of 0.53 SD – a form of regression to the mean, produced entirely by measurement error (Barnett et al., 2005; see also Cronbach and Furby, 1970).

As a fourth example, the simulation illustrates how design decisions can influence what an intervention trial might find. A therapy that clears 30% of plaque was tested in 100 simulated trials modeled on the A4 study (Sperling et al., 2014, 2023), each enrolling 1,200 amyloid-positive (at A4’s cutoff; Methods), cognitively unimpaired adults aged 65–85, measuring episodic memory every six months and analyzed after 2, 4, 6, 8 and 10 years. Because each participant’s life is also simulated without the intervention, the true effect in the very same people is known. That effect was small and grew with trial length, with the power to detect it growing from 9% to 25% (Figure 4D). One of the 500 analyses, at 4 years, found significant harm by chance. The A4 trial itself found no slowing of cognitive decline with solanezumab over 240 weeks (Sperling et al., 2023). The simulation says little about that treatment, but it shows how, in this world where AD acts slowly on cognition, the length of the trial substantively impacts what can be detected. The level of AD pathology at baseline also affects the rate of decline that can be observed within a given period, such that cognitively unimpaired people with high levels of amyloid and tau show accelerated decline (Sperling et al., 2024; Buckley et al., 2026).

As a fifth example, the simulation can apply competing definitions to the same people. Under five published definitions of “superaging” that compare older with younger adults (Harrison et al., 2012 and Rogalski et al., 2013; Dang et al., 2019; Sun et al., 2016; Maccora et al., 2021; Bott et al., 2017), between 1.5% and 19.4% of participants in the same real-world sample qualified (Powell et al., 2024). The same definitions were applied to the 14,405 dementia-free simulated people aged 70–90 (Figure 4E). The three definitions that compare memory at one visit with that of younger adults classified 6.4%, 19.9% and 15.7%, against 8.4%, 18.5% and 19.4% in the real sample, and the first definition, which is limited to those 80 or older, classified 20% of them, against 21%. The definition that requires memory to be maintained over three waves classified 6.0%, against 1.5%, and the one based on processing speed 27.8%, against 6.5%: simulated people aged 60–80 without cognitive impairment are 1.4 young-adult SDs slower than people in their 20s, against 2.3 in the cohort in which that definition was devised (Bott et al., 2017). Overlap in who each definition selected was modest.

Finally, demonstrating the flexibility of the simulator, worlds can be compared that differ in their core assumptions (Figure 4F). In an analysis based on Fjell et al. (2026), three worlds were modeled that share the same simulated people but differ in terms of how memory declines. One world is without generational gains in episodic memory. In the second, default world, birth in later decades leads to higher levels but all decline at the same rate in older age. In a third world, later cohorts also decline more slowly. Plotted as change from age 35, each birth cohort’s true change can be set against everyone alive in 2026. Without generational gains, the cross-sectional pattern follows the cohorts until about 70 and then falls less than people do, 1.41 SD by 85 against about 2.0 SD, because those who decline are most often lost to the sample. With gains in level alone, the cross-sectional pattern falls 0.49 SD by 60 while each cohort loses 0.10–0.12 SD, the gap seen in Figure 4B, and changes chained across age groups recover the true change. With slower generational decline added, the cohorts fan out. By 85, those born in 1931 have lost 2.37 SD and those born in 1961 0.62 SD, and changes chained across the age groups of 2026 show 0.97 SD, well short of the 1.50 SD lost by the 85-year-olds born in 1941. When generations decline at different rates, neither the existing cross-sectional patterns nor chained change recovers the true values. Such observations could be used to revise the underlying model of the simulator in future versions, including expansions for selective mortality and the numerous other factors that differentially shape populations before they enter midlife (see Supplementary Figure 7).

## Example of an Insight that Emerged from a Simulation Failure

The simulation is generative in a further sense as it forces critical evaluation of how well the hypothesized causal cascades accommodate real-world data. Figure 6 follows one example of a failure that led to a recorded insight, as analyzed in version 88, and then another in version 90.

**Figure 6.**
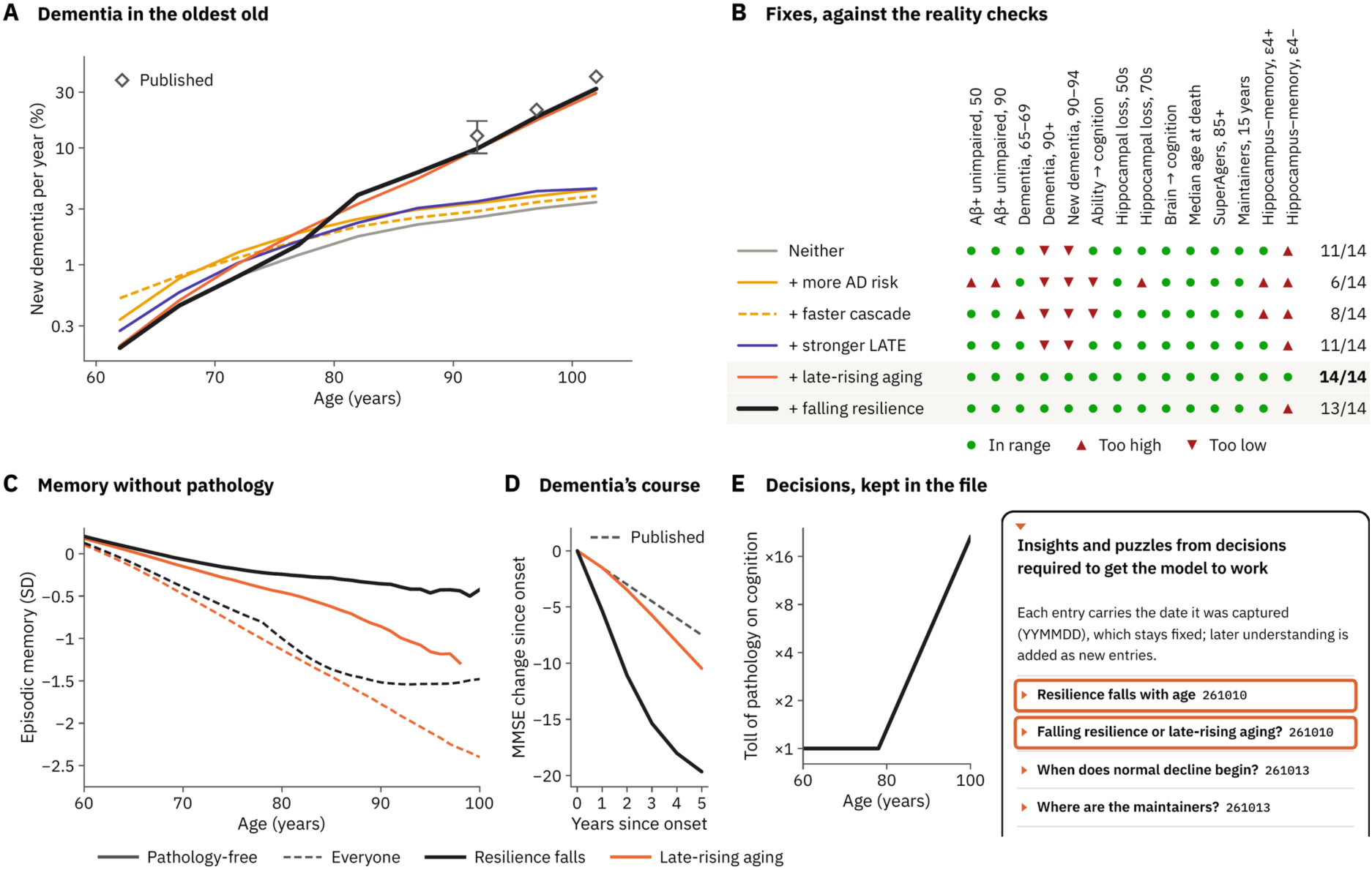
A failure of the model yields two competing hypotheses: aging may accelerate, or resilience may fall, late in life. (**A**) New dementia per year (%) by 5-year age group in six variants of the version 88 world, with published rates plotted at 90–94, 95–99 and 100 and older (diamonds; bar, the accepted range at 90–94; Corrada et al., 2010). Gray, the original model; amber, more AD risk; amber dashed, a faster AD cascade; purple, stronger LATE; orange, aging that accelerates late in life (the default in version 88); black, resilience that falls with age (the default from version 75 to version 87). (**B**) Each variant is shown against version 88’s fourteen reality checks: green, within range; red triangles, above or below it. (**C**) Mean episodic memory by age among people free of pathology (solid) and among everyone (dashed), with falling resilience (black) or late-rising aging (orange). (**D**) Dementia’s course plots the mean change in MMSE since onset for dementia beginning at 80–89, with falling resilience (black) or late-rising aging (orange), against 1.5 points a year for Alzheimer dementia in the Cache County study (dashed line estimated from Tschanz et al., 2011). (**E**) The decisions, kept in the file. Left, the toll of AD, LATE and white matter lesions on cognition is plotted under falling resilience. Right, the application’s list of insights and puzzles, which opens with the entry for the decision to let resilience fall with age (outlined), followed by the puzzle that it left open (outlined). Some attempted repairs fell short. The two highlighted here both brought dementia in the oldest old into range but differed in which people were affected by the newly hypothesized cascades, pointing to prospective data that could decide between the two alternative hypotheses.

With general effects of aging being set as small relative to the AD cascade, and an AD cascade calibrated to the amyloid checks, the model could not produce the amount of dementia expected for the oldest old. Dementia prevalence at 90 and older was 13% against a published 35%, and incidence at 90–94 was 2.6% per year against 12.7 per 100 person-years (11.9% a year; Figures 6A and 6B). As the application’s record of this decision explains, it took a median of about 22 years in that world to progress from amyloid positivity to dementia. Dementia therefore rose only about sevenfold from the late 60s to the late 90s, where populations show a twentyfold to thirtyfold rise (Jorm and Jolley, 1998; Corrada et al., 2010). Certain attempted repairs to the simulation model fell short (Figures 6A and 6B). With more AD risk, a faster cascade or stronger LATE, dementia at 90 and older stayed between 15% and 17%, and incidence at 90–94 between 2.9% and 3.5% per year. Two of them also failed checks that were not targeted. Raising AD risk raised dementia at every age rather than steepening its rise and pushed five other checks out of range, among them amyloid positivity among unimpaired 50- and 90-year-olds (Jansen et al., 2015), and a faster cascade raised dementia at 65– 69 too far. Those amyloid checks initially drew on pooled research cohorts; later versions anchor both amyloid onset and the amyloid checks to the population-based Mayo Clinic Study of Aging instead (Roberts et al., 2018).

However, two repairs brought dementia in the oldest old into range: normal aging that accelerates late in life, and resilience that falls with age. Resilience that falls with aging was operationalized as a multiplier on the cognitive toll of AD, LATE and white matter lesions that applies to all lives and doubles every five years from age 78, so that the same pathology does twice the harm to cognition at 83 and four times the harm at 88. Late-rising aging passed all fourteen of version 88’s checks, and falling resilience thirteen, missing one by a small margin (a correlation of 0.17 between hippocampal and memory change in non-carriers of APOE ε4, against an upper limit of 0.16).

The two surviving hypotheses fit the checks similarly, each closer to the reference values on some of them. Interestingly, and reinforcing that they are distinct mechanistic hypotheses, the two alternatives put the decline of the oldest old into different people (Figure 6C). Pathology-free 90-year-olds average −0.86 SD with late-rising aging but −0.35 SD with falling resilience, about the same as with neither repair (−0.34 SD), because the fall multiplies only the toll of pathology.

Returning to the simulation, falling resilience was initially chosen as a mechanism to incorporate in version 75, and the choice was captured inside the application as the first entry under “Insights and puzzles from decisions required to get the model to work” (Figure 6E). The entry recorded what failed and why, what was decided, and its limits, and the next entry records the still-open puzzle. A later check reversed the choice. Once every life was staged by the CDR (version 87), dementia that began in the 80s was seen to progress three to four times as fast as Alzheimer dementia in the Cache County study (Tschanz et al., 2011), and version 88 made late-rising aging the default, recording why in a new entry, “Early dementia passes too quickly.”

The reversal does not resolve the issue, because each hypothesis fits some aspects of real data at a cost to other aspects. Falling resilience spares people without pathology, whose memory declines little in their 90s, but it makes dementia progress faster than populations show. Late-rising aging gives dementia a course closer to the one observed, but it lowers everyone’s episodic memory in the 80s and 90s, and it makes dementia that begins at 90 or later progress faster than dementia that begins earlier, where the Cache County study found, if anything, the reverse. Published findings offer conflicting constraints. Eleven indices of common age-related neuropathologies together explained only 43% of the differences between people in the rate of late-life cognitive decline (Boyle et al., 2021), which leaves room for additional cascades, whereas the faster decline that comes with combined AD and LATE pathology was about 40% weaker in people who died at 90 or older than in those who died younger, the opposite of a toll that grows with age, although the authors note that differential survival may contribute (Kapasi et al., 2020). Either alternative mechanism should be viewed as a hypothesis to be tested by novel analyses of real-world data. Specifically, here, episodic memory trajectories in very old people with pathology measured, at autopsy or by biomarkers, might help decide between the two alternatives.

The simulation continued to evolve. Explorations from version 90 onwards took up existing clinical-pathologic data: only 22% of people in the 90+ Study who died without any of the common pathologies had dementia (Kawas et al., 2015), and in the Rush cohorts cognitive decline sped up more than sevenfold about 3 years before death (Boyle et al., 2013). Based on these constraints, version 90 added terminal decline as a seventh chain, cutting aging’s late rise, and recording why in a new entry, “Where is terminal decline?” And version 100 expanded the AD cascade to explore direct and indirect effects of amyloid and tau levels on decline. These revisions, inspired by grappling with the data through simulation, reflect the kinds of insights that can be gleaned.

## Self-Documenting Code

The simulator is self-encapsulated (Supplementary Figure 1). Everything is in one HTML file that opens in a web browser without installation or a server, and can simulate the 120,000 lives in under ten seconds on a laptop. The file is written to be read by its next developer, who may be a person or an AI agent. Its first lines tell the reader to begin with the developer guide inside the file. The developer guide covers what the simulator is, how to add a stage, and how to capture insights and puzzles (Supplementary Figure 1B). The equations under “How the model works” and the reality checks are recomputed with every new world. For example, switching off aging’s late rise rewrites the description and moves checks out of range (Supplementary Figure 1C). Settings and a copy of the application itself can be saved (Supplementary Figure 1D). This strategy thus incorporates literate programming (Knuth, 1984) and documentation embedding (Wilson et al., 2014) at the scale of a whole scientific model, so that it can be iteratively modified by future researchers co-working with AI agents.

## The Simulator Is Designed to Evolve

The Imagine stage builds into the application a route to flexibly change the simulation and its outputs (Figure 7A). This is a key feature. A researcher downloads the page, attaches it to a new conversation with an AI agent, and pastes a prompt describing a new desired stage. The current version of the application provides examples (e.g., Biomarker timelines, Compare definitions, Alternative worlds and Childhood mortality worlds) as well as a template for one’s own novel request. The AI agent then reads the guide, builds the stage in Imagine’s slot, and returns the complete file, which the researcher can explore and further modify in the same conversation.

**Figure 7.**
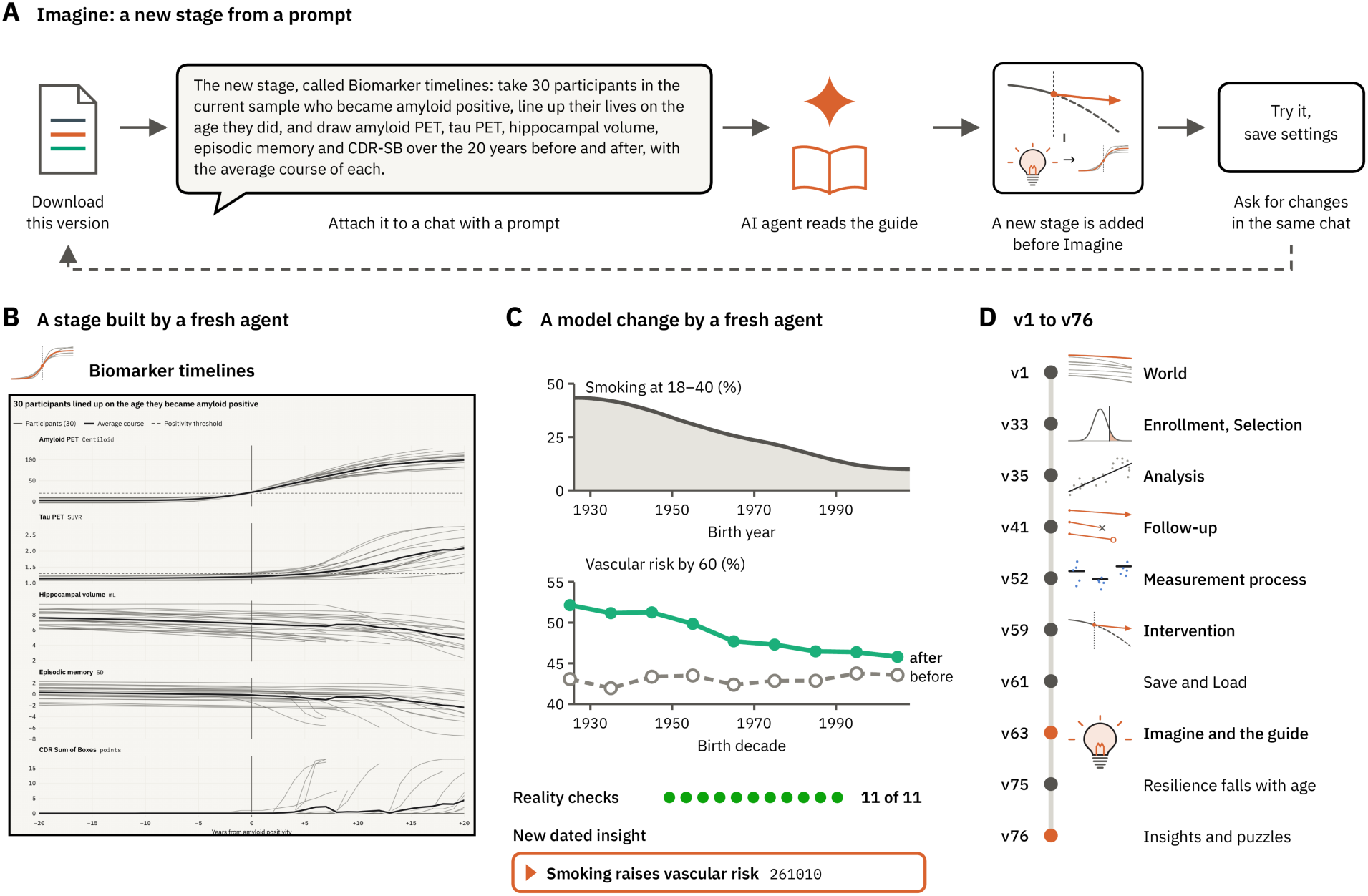
The simulator evolves by adding new stages and model changes. (**A**) The Imagine stage provides a route to flexibly create an entirely new stage. The user downloads the page and attaches it to a conversation with an AI agent with a prompt (the Biomarker timelines example is shown). The AI agent reads the guide inside the file, the new stage is added in Imagine’s slot, with Imagine kept after it, and the researcher tries it, saves settings and asks for any desired changes. The next idea then starts from the new file. (**B**) An example stage built by a fresh agent from the file and the Biomarker timelines prompt alone, in 3.6 hours. Thirty participants are plotted lined up on the age at which they became amyloid positive (year 0), with amyloid PET, tau PET, hippocampal volume, episodic memory and CDR Sum of Boxes over the 20 years before and after, and their average course (bold). (**C**) An example model change built by a second fresh agent, in 1.8 hours, that focused on a cohort effect. Top, smoking exposure is plotted by birth year, which raises the hazard of vascular risk. Middle, vascular risk by age 60 is plotted by birth decade before (gray) and after (green) the model change. All reality checks remained in range, and the change was recorded as a new dated insight (bottom). (**D**) Milestones from version 1 to version 76, with the stages’ icons. The same file that runs the model carries the instructions to change it, so each change, by a person or an AI agent, starts from the available version and ends with a new, documented version of the simulator.

To test whether the file alone suffices, two fresh AI agents were provided the application, with no access to the development history, only the version 78 file and a request. The first request was the Biomarker timelines example. Working autonomously for about 3.6 hours, the agent built a stage that lines up participants on the age at which they became amyloid positive and plots amyloid PET, tau PET, hippocampal volume, episodic memory and CDR Sum of Boxes over the 20 years before and after, with the average course (Figure 7B).

The second request changed the core model: link the curve of smoking prevalence among the simulator’s cohort effects to vascular risk, so that people whose early adulthood fell in years of heavier smoking carry more vascular risk. In about 1.8 hours the agent implemented the link, updated the settings and description of the cardiovascular chain, and reported the reality checks. Six moved slightly and all eleven available within version 78 stayed within range (dementia at 90 and older went from 40.5% to 40.7%; Figure 7C). Vascular risk by 60 rose from 43% to 48% overall, more in early cohorts, which smoked more (52% for people born in the 1920s against 46% for those born in the 2000s). The agent recorded the change as a new dated entry, “Smoking raises vascular risk,” including the settings at which it would push a check out of range. Supplementary Figure 5 shows outputs from all examples offered on the Imagine stage, and Supplementary Figures 6 and 7 follow two of these builds in detail. When building these novel stages, both agents read much of the code beyond the guide and reported where the guide fell short. It explained how to add a stage but not how to change the model; parts of the file had drifted; and a note on one reality check said the model lacked a link that the code in fact contained. Such reports are themselves inputs to the next version. The guide evolves along with the model. The whole simulator grew this way, from the World stage of version 1 to the capture of Insights and puzzles of version 76 (Figure 7D).

## A Simulator Enables a Generative Loop

The simulator described in this paper enables a generative loop. Figure 8 contrasts two routes from data to understanding. Within the familiar framework of normal science (Kuhn, 1962; Platt, 1964; Box, 1976), studies sample an unobserved real world, each through its own enrollment, selection strategy and measurement process, and the results are integrated into a theory that generates the next hypotheses (Figure 8A). In a simulator, the world is generated from explicit assumptions based on past findings, so every life and every cause in it is known (Figure 8B). Simulated studies sample that world as real studies sample ours. The results can then be compared with the true values inside the simulation and novel data from the real world. When a comparison fails, or succeeds for an unexpected reason, the insight or unresolved puzzle can be captured, and a person or an AI agent can revise or extend the model to make the next version.

**Figure 8.**
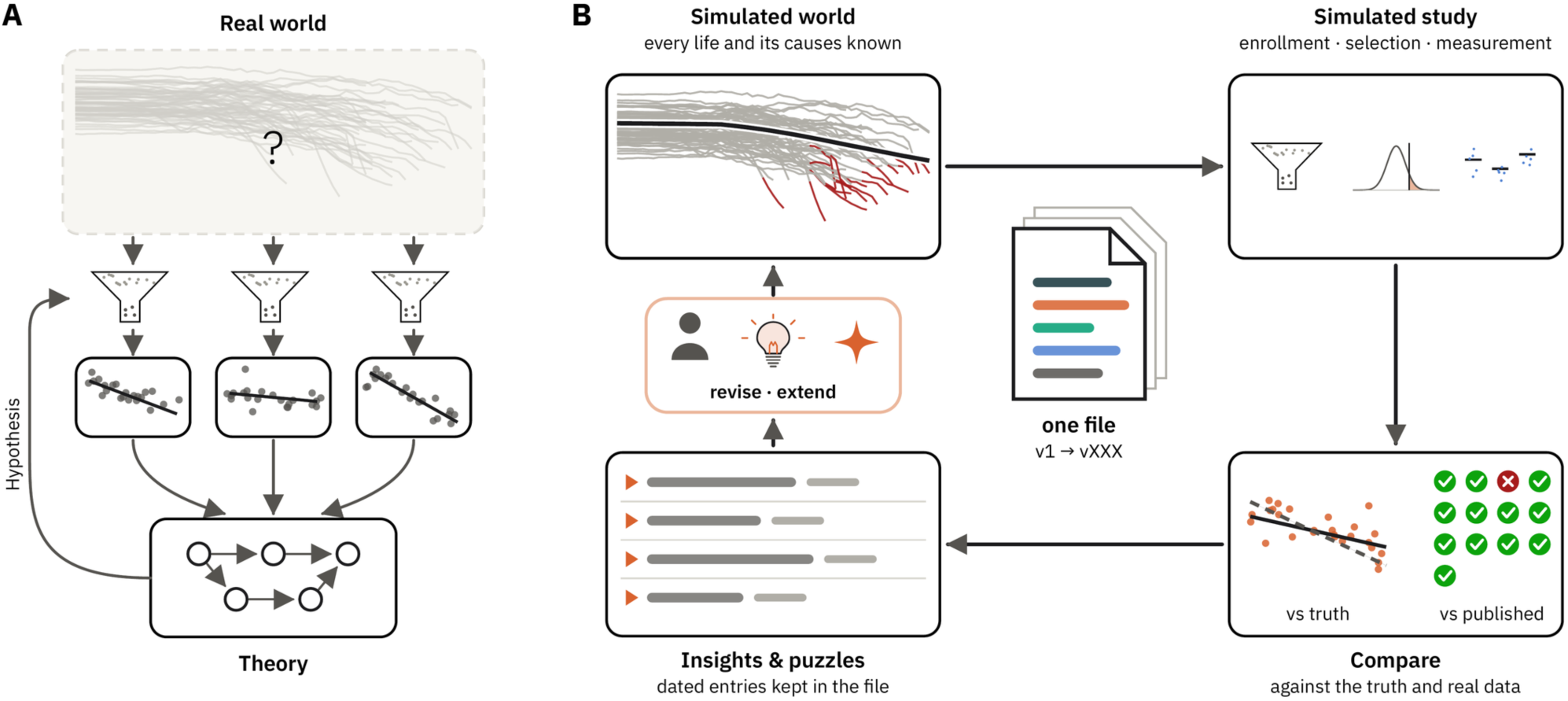
A simulator as generative loop. (**A**) Normal science. Studies sample a real world that is never observed directly (gray lives and question mark). Each study yields data (scatterplots with fitted lines) through its own enrollment and selection (funnels). The data are integrated into a theory (circles and arrows), and the theory generates hypotheses for the next studies to test (left arrow). (**B**) The simulator’s loop. A simulated world is generated from explicit assumptions based on curated existing data, so every life and its causes are known (gray lines plot episodic memory of individual lives by age with red showing dementia; the black line is the mean). Simulated studies enroll, select and measure participants (top right). The results are compared with the true values (a fitted line against the true relationship, dashed) and with published data (reality checks; green, within range; red, outside it). What the comparison teaches is captured as dated entries (Insights & puzzles, bottom left). Then, a person or an AI agent revises the model or extends it (orange), producing the next version (center). The simulator shows what a set of assumptions and a study design can recover. Prospective experiments then can test the assumptions that generate those values. Thus, the simulator does not replace data collection but rather organizes existing data and identifies gaps and possibilities for future collections.

## Limitations and Future Directions

The simulator should be viewed as a series of hypotheses, not results obtained in the real world. Its causal chains are assumptions, and its parameters are choices that cannot be unambiguously resolved without prospective data. In this sense, the simulator does not replace real-world data collection but rather organizes it, formally models its implications under a series of assumptions, and identifies gaps that can only be filled with new data collection.

For example, the default world takes the practice-adjusted longitudinal estimate for episodic memory as its working assumption and keeps early decline for speed; this is a setting. Practice gains, too, are the same at every age and level of impairment, whereas in real data they shrink with age and are smaller in MCI and AD (Salthouse, 2010; Goldberg et al., 2015). Its benchmarks come mainly from populations in the United States and Europe. Moreover, the current model leaves out a great deal that is known about brain and cognitive aging, including co-pathologies such as arteriolosclerosis, cerebral amyloid angiopathy and Lewy bodies; social and structural determinants of health beyond SES and schooling; and most potentially modifiable risk factors (Livingston et al., 2024). In addition, there are a multitude of factors that affect survivorship, including child mortality, that differ over time and are not evenly distributed across SES and health. Expanding beyond those currently included in the simulator is a future direction (e.g., Supplementary Figure 7). As an example of data limitations, experimental data to constrain aging patterns in individuals above 80 have historically been underrepresented in studies of cognitive and brain aging, reflecting enrollment age restrictions, selective participation, and attrition (Merenstein and Bennett, 2022). It is thus interesting that, when the model could not produce the amount of dementia expected for the oldest old, the simulator self-documented in the captured Insights (Figure 6) that more data in the oldest individuals with longitudinal cognitive assessment and markers of pathology would be needed to test between the competing hypotheses.

Another limitation is the use of simplified models of causal cascades. For example, in the simulator’s AD cascade, amyloid is a trigger that causes tau to aggregate; tau aggregation into neurofibrillary tangles is then directly linked to accelerated neuronal death and cognitive decline (see Supplementary Figure 3). This is an incomplete model. It has no regional tau staging, does not account for tau propagation across neural systems, and has no AD-related atrophy or decline before tau positivity, both of which might occur in real cohorts (Jack et al., 2024). Current criteria stage AD biologically by how far tau has spread: tau most commonly begins in the medial temporal lobe, amyloid facilitates its spread into the neocortex (Jack et al., 2024), and neocortical tau was the strongest predictor of cognitive decline among cognitively unimpaired people with tau PET in A4 and LEARN (Sperling et al., 2024). Hippocampal atrophy also reflects more than medial temporal tau, as TDP-43 pathology and vascular disease contribute both independently and together with AD pathology (Small et al., 2011; Nelson et al., 2019), which the simulator’s LATE and cardiovascular chains capture in part.

A more sophisticated cascade could explore interactions between amyloid and tau, in keeping with evidence that amyloid does more than trigger tau pathology and acts synergistically with it (Busche and Hyman, 2020), or consider the possibility that non-tangle aggregates play a role in neurodegeneration. Alternatives that consider the rate of tau accumulation and seeding into neocortical regions, and interactions between amyloid and tau, prepare the model to explore predictions of specific mechanisms of toxicity and the impacts of intervention. Adding tau acceleration as a direct correlate of neurodegeneration specifically allows the simulator to accommodate the slowing of cognitive decline seen when antibodies clear plaques in early symptomatic AD (e.g., van Dyck et al., 2023; as assumed for Figure 4D). Version 100 adds a module to explore an expanded AD cascade while retaining the simplified model explored in the present paper. This extended model is one example of how an alternative causal cascade can be implemented and tested.

As another example of a simplification, the default aging cascade is set to change phenotypes in relation to age without any specific proposed mechanisms. Such a cascade is useful because it allows separation from other cascades associated with known biology that aggressively affect subsets of individuals (e.g., the AD cascade). However, it is incomplete because it becomes a nonspecific catch-all for processes affecting aging that are not otherwise labeled or understood. The presence of an aging cascade module should not be interpreted as evidence of a general aging mechanism (for discussion, see Meyer and Schumacher, 2024; Ikram, 2024). Rather, it is an agnostic placeholder for processes affecting aging that are not yet identified. As additional cascades are identified and built into the simulator, we anticipate that the nonspecific aging cascade will shrink and potentially even become unnecessary. An ideal simulator would have cascades for all biologically driven processes that are correlated with age and affect brain and cognition, and individual lives would simply be the aggregate of the interactions among them.

A final limitation, common to the emerging use of AI in experimental science, is that code written with an AI agent can contain errors that look plausible. The simulator was checked against the reality checks, by automated tests comparing every stage with earlier versions, by loading settings saved by earlier versions, and by the fresh-agent tests, whose reports led to corrections of errors and of documentation that had fallen behind the code. None of these checks replace scientific review of the assumptions, which remains the task of the field and experts. This reinforces the practical utility of a simulator able to evolve. An investigator who catches an error or disagrees with an assumption or desires a more nuanced and elaborated causal cascade can modify the simulation and rapidly assess the effects.

## Conclusions

The AgingBrain Simulator is provided as a generative, extensible model of brain and cognitive aging. By making explicit the totality of the data patterns that result from a hypothesis, simulations illustrate how a hypothesis can fail even when explaining a focused-upon result. The simulator’s value will grow if others change it, replace a causal chain with a better one, calibrate it to new data, add a stage that asks a new question, or adapt it to a different population (e.g., focused on childhood or adolescent development). Because it is a single file that explains itself, a researcher working with an AI agent can make such a change in hours, test it against the same checks or develop new ones, and share it as a new version of the simulator. The record of insights and puzzles captures what was learned, and the insights point to real-world experimental data that can resolve competing hypotheses. Whether the approach succeeds will depend less on any version of this model than on whether the field adopts shared, generative models of the aging brain.

## Methods

### Simulator

Results come from AgingBrain_Simulator_261023.html (version 99), except those in Figure 3, Figure 4D and Table 1, which come from AgingBrain_Simulator_v104_261115.html (version 104), those in Figure 6, which come from AgingBrain_Simulator_261014.html (version 88), and the two alternative worlds in Figure 4F, which use the settings for generational change of an experimental branch, AgingBrain_Simulator_v91_261018.html. Unless stated otherwise, the results use the default settings (world seed 7; 120,000 lives born from 1926 to 2008; study year 2026, in which 89,554 of the lives are alive). Figures 4 and 6 and Supplementary Figure 4 vary settings as described below (for Supplementary Figure 4, in its legend), and Figures 7B and 7C show files produced by the test agents that are not part of the distributions but can be constructed. In headless Chromium on a two-core cloud machine, the 120,000 lives were simulated in 6.9 seconds.

### The simulator’s parameters

The simulator is structured as a set of baseline determinants and causal cascades that target biological and cognitive constructs. The probabilities and rate constants were set to align with targets derived from real-world data. Each empirical target came from a published study or preprint, found by an AI agent searching PubMed, medRxiv, bioRxiv, and PsyArXiv, or supplied by the authors as a paper figure or table, supplementary file or value. Values reported only in figures were read from the plots and are approximate to the figure’s resolution; the rest came from text or tables. The authors continuously checked the values against their sources and confirmed key parameters. All parameters are explicitly exposed and can be visualized inside the application, and saved and loaded to establish separable parameter sets.

### Measurement and sampling

Measurements used the true value plus a normally distributed error. The default setting errors are 0.40 SD for episodic memory and executive function, 0.35 for processing speed, 0.30 for semantic memory, 1.2 points for the MMSE (then rounded to whole points), 0.15 mL for hippocampal volume, and 8 mL for intracranial volume. Cognitive scores are also in SD units, which approximate z-scores of 50- to 65-year-olds. A person receives the same error on a given occasion by default wherever that measurement is used. Note also that the Measurement process stage allows these error settings to be changed including differences in the error SD as a function of age, CDR Sum of Boxes, and sex.

Behavioral measures included practice gains across repetitions that fade over time. For episodic memory, the practice left by each test halved every 3 years, with a gain of 0.25 SD at a retest immediately after a single test and an asymptote at 0.45 SD with repeated testing; a retest five years after one test gains 0.10 SD. Samples were drawn using the Selection stage from everyone who met a strategy’s criteria in true values (all who enrolled under the application’s default recruitment) at random and without replacement, at the strategy’s share of women (55% for Random, Nondemented and MCI; 60% for SuperAgers and A4-like trials).

### Combining measures across studies

The simulator expresses each construct on a single scale, which requires values from published studies to be mapped onto it. Amyloid PET is measured in Centiloids (Klunk et al., 2015), which the model converts, as (Centiloid − 12) / 88, to an amyloid load on its own scale, 0 at 12 Centiloid or less and 1 at 100 Centiloid, that scales amyloid’s effects. The positivity threshold of 20 Centiloid (a load of 0.09) is intended to correspond approximately to a PiB SUVR above 1.42 in the Mayo Clinic Study of Aging (Jack et al., 2017; Roberts et al., 2018). The amyloid-negative threshold of 12 Centiloid (Figures 4A and 5) is where the model’s amyloid load begins, so people assigned as amyloid negative carry no amyloid load, whereas someone at 15 Centiloid, negative by the 20 Centiloid threshold, already carries 0.03 amyloid load. The A4 study used florbetapir SUVR combined with a visual read; its quantitative cut, an SUVR of 1.15, was approximated for Figure 4D to correspond to 33 Centiloid (a load of 0.24) (Sperling et al., 2024). The simulated trials in Figure 4D also required a CDR Sum of Boxes of 0, whereas A4 required a global CDR score of 0, an MMSE of 25–30 and a Logical Memory delayed recall score of 6–18. Tau PET is a single SUVR in a temporal region-of-interest (ROI), positive at 1.30, with a lower (conservative) threshold of 1.25 for excluding biomarker-positive people. The default simulation does not model tau PET spread.

Each cognitive domain is represented by one score, in SD units that approximate z-scores of 50- to 65-year-olds. The SuperAger rule’s cutoff on delayed recall of the Rey Auditory Verbal Learning Test became the mean episodic memory of dementia-free 56- to 66-year-olds, and its age- and education-adjusted limits on other tests became limits on executive function and semantic memory adjusted for age alone (see Figure 4C below). The rule of Josefsson et al. (2012), defined on a composite score based on several of Betula’s episodic memory tests, was applied to the simulator’s single episodic memory score. A composite’s higher reliability can be mimicked in the Measurement process stage by lowering a measure’s error; the defaults use an error of 0.40 SD for episodic memory (a test–retest r of 0.89) and perfect validity (1). The published stability of one general cognitive ability / intelligence test from ages 11 to 79 (Gow et al., 2011) served as the benchmark for the simulated correlation of early-life ability with a composite of memory, speed and executive function at older age. Clinical stages that the source studies diagnosed by consensus or by algorithm are defined here by CDR Sum of Boxes (unimpaired at 0, MCI at 0.5–2.5) and by the dementia threshold in global cognition.

### Reality checks

The reality check values are shown in “The properties of this simulated world” section of the World stage with their intended values and ranges from the literature. When parameters are changed that affect the simulated world, these reality check values automatically update. The checks come from Roberts et al. (2018); Manly et al. (2022); Corrada et al. (2010); Gow et al. (2011); Fraser et al. (2015); Social Security Administration (2026); Josefsson et al. (2012); Gorbach et al. (2020). Each check counts the people its source studied: the same calendar years or birth cohorts, or, where the source’s participants were born before the world’s first births (1926), the world’s ten earliest birth years at the same ages. The hippocampus–memory check includes only the ages the world covers at its source’s first tests (55–84 years, against 55–90 years). Amyloid positivity is 20 Centiloid or more (a PiB SUVR above 1.42 in the source), the cognitively unimpaired have a CDR Sum of Boxes of 0, and both amyloid checks are shares of everyone without dementia. New dementia per year is a yearly risk, converted from the published rate per 100 person-years as 1 − exp(−rate). The correlation with global cognition is an observed value, allowing for test reliabilities of 0.85 (early-life ability) and 0.80 (late-life cognition). The published value for early-life ability is the stability of one reasoning test as noted above. Median age at death comes from a period life table of the world’s deaths in 2023.

In Figures 2, 4C and 5, the SuperAger rule is applied to true values, with norms from 2026 (in Figure 4C, also to one measurement of each test, with the same thresholds). In the specific comparison of the two hypotheses of Figure 6, which used a rule that was subsequently updated, SuperAgers were defined by memory alone using episodic memory at or above the mean of dementia-free 50- to 65-year-olds (after Harrison et al., 2012). Neither rule adjusts the memory cutoff for education; the published limits on the other tests use norms for age and education (Harrison et al., 2012; Rogalski et al., 2013). Maintainers are found by the rule of Josefsson et al. (2012) in a simulated study whose participants were tested every 5 years from 1990 to 2005, with measurement error. Because the rule’s cutoff is 1 SD at the final score, it will yield about 16% for any roughly normal spread. The correlations of hippocampal and memory change come from a simulated two-visit study in 2010 and 2014, with measurement error. The ranges are tolerances set around each published value and, for the amyloid checks and the hippocampal–memory correlations, the published 95% confidence intervals.

### Figure construction

The detailed methods for each figure are described below. Measurement error was added after selection based on the parameters defined in the Measurement process stage, except where noted, and used as inputs to analyses. Analysis scripts computed each summary value plotted in figures (regressions, correlations, medians, group means, chained changes and shares), except that in Figure 4D the application’s own Intervention code fitted each participant’s slope and ran Welch’s t. The complete set of scripts is available in Markdown in the GitHub repository (https://buckner-lab.github.io/Aging-Brain-Simulator). For figure generation, the application was run in headless Chromium through Playwright (Figures 1–8 and Supplementary Figure 4) using a copy of the file with two changes that did not alter derived analyses. One change added a line that exposed its internal functions to the analysis scripts; the other raised the cap on the canvas pixel ratio for sharper screenshots. For Figure 1, a few further changes affected how plots were drawn (axes, tick labels and the ages shown in Follow-up), and for Figure 4F the engine was modified to include the generational settings. Settings for each main-figure analysis are given below; those for the Supplementary Figures are provided in their legends. Median age at death was computed as in the application, by interpolating survival from 40. Figures 6 and 7 and several Supplementary Figures include screenshots of the application.

### Figures 1 and 8

In Figure 8B, dementia begins when global cognition falls to −2.3 SD after a decline of at least 1.2 SD from its level at 40, at which point the MMSE is about 24. In Figure 1B, each chain’s effect is the difference it makes to the simulator’s reference person at 85: onset of the cardiovascular chain, the AD cascade and LATE at 55, 65 and 80, respectively, against no onset, and a terminal phase from 82 against none. Figures 1C–1H show one simulated study as the application runs it with the Nondemented strategy: 400 people aged 60–90 in 2026 (55% women) with MMSE ≥ 24, with episodic memory measured with the default error (SD 0.4). Figure 1F shows a second sample, drawn with the same criteria (selection seed 282; Figures 1D, 1E, 1G and 1H use seed 1). In Figure 1F, biomarker positive means amyloid PET ≥ 20 Centiloid (a load of 0.09) or tau PET ≥ 1.30 SUVR in 2026. In Figure 1G, visits are every two years and include practice effects, and, in Figure 1H, change is in true episodic memory, without measurement error or practice, over 10 years of follow-up.

### Figure 2

Figure 2 presents a comprehensive analysis of the effects of selection criteria. For simplicity, it uses only women. Results are similar for male or mixed cohorts. The analysis included every woman in the default world who lived to 85, whatever her birth year (31,702 women, born 1926– 2008, who turn 85 between 2011 and 2093), classified on true values at 85. Note that the simulator can restrict analysis to an arbitrary date (e.g., turn 85 in 2026) or set a criterion applied to all lives, as applied here, which includes individuals who will die in the future.

*MCI* at 85 was defined as a CDR Sum of Boxes of 0.5–2.5 without dementia (6,894 women). *SuperAgers* met the Harrison et al. (2012) and Rogalski et al. (2013) rule at 85, the limits for executive function and semantic memory being those of dementia-free 85- to 89-year-olds (6,294 women). *Josefsson maintainers* met the rule of Josefsson et al. (2012), applied to true episodic memory across four repeated measures (at 70, 75, 80 and 85). They were defined as women having memory at 70 plus 15 times the least-squares slope per year through the four values that was at least 1 SD above the same value from the mean of all women free of dementia at 85 (4,445 women). *Slope maintainers* had lost no more than 0.3 SD of episodic memory since 75 (5,730 women).

In panel A, each phenotype’s true change from 40 is divided by its spread among everyone alive at 85 (the 10th–90th percentile range divided by 2.563, with white matter lesions on a log scale), as in the application’s Life in detail expandable banner in the World stage. MCI marks the first age with a CDR Sum of Boxes of 0.5 or more, before dementia. In panel B, the engine splits each woman’s episodic memory at each age into the level set at 40 and each chain’s contribution at that age. Aging comprises aging’s losses independent of any other causal cascade and the person’s own random drift. Each group’s pie chart insert illustrates the selected group’s mean difference from the dementia-free women at 85; negative shares (in the slope maintainers’ pie, the level at 40, −11%, and a negligible cardiovascular share) were set to zero and the rest rescaled to 100%. The value written below is the group’s mean level at 40 relative to hers. The pie under the median woman gives each part’s share of the differences in memory at 85 among all 31,702 women (its covariance with memory divided by memory’s variance). In panel C, memory is each woman’s true episodic memory level at 40, and change is her true memory at 85 less that at 75. High is defined as memory above the median at 40 of the women free of dementia at 85 (+0.68 SD), and stable means a loss of no more than 0.3 SD in slope, the slope maintainers’ criterion. Shares are of each group’s own members and, under the median woman, of all women free of dementia at 85. Each of the five panels shows the same 3,000 women free of dementia at 85, drawn at random.

Panel D used the same procedures as A-C but added error before selection using the true value plus a normally distributed error with a 0.40 SD for episodic memory and executive function and 0.30 for semantic memory, and of 0.5 points for CDR Sum of Boxes, which was then rounded to half points. No practice gains were included. Episodic memory was measured at 40, 70, 75, 80 and 85, and executive function, semantic memory and CDR Sum of Boxes at 85. Dementia status was taken as known, so the population is the same 27,059 women matched to panel C. The criteria and their cutoffs are matched with panel C so only each woman’s measurement error differs between panels C and D. The same 3,000 women in panel C are plotted in panel D and, for each group, the same 900 selected members are plotted, at their measured values with error (they are plotted black if also selected on measured values, and dark green if missed). The incorrectly included (plotted red) were sampled at the same rate as those 900 members (817, 153, 211 and 716 women).

The Jaccard index is defined as the number of women selected on both true and measured values divided by the number selected on either, that is, correct / (correct + incorrectly included + missed). The Jaccard index is 100% if measurement error changes no person’s selection and 0% if every person is selected differently between true and measured values. Unlike the share of all women classified correctly, it falls with both kinds of error providing a useful single measure of the selection strategy’s sensitivity to measurement error. The first panel (all women) applies no selection criterion and so has no Jaccard index.

### Figure 3

In panel B, ages are shown while at least 30 lives remain. In panel C, the amyloid positivity threshold is 20 Centiloid (a load of 0.09) and cognitively unimpaired means a CDR Sum of Boxes of 0. The curves pool every year of every life, whereas the checks in panel E count only the people each source studied (see Reality checks). In panel D, the shares come from a 30,000-life sample, among the living every 5 years. Negative shares were set to zero and the rest rescaled.

### Figure 4A

The population was everyone alive in 2026 aged 60–90 (29,096 people). The criteria, in true values, were none (Random), an MMSE ≥ 24 (the Nondemented strategy; 27,366 people), and the same with amyloid PET below 12 Centiloid and tau PET below 1.25 SUVR (that strategy’s option to exclude biomarker-positive people, at its default thresholds; 15,458 people). Each criterion was sampled 100 times (220 women and 180 men), and each person’s episodic memory was measured once. In each sample, measured memory was regressed on age by ordinary least squares without covariates, and the slope was multiplied by 10. The reference slopes (black bars and dashed line) and the left plot (mean memory at each year of age, smoothed by a centered 3-year running mean) used true values, among all living individuals meeting the criteria.

### Figure 4B

Everyone alive in 2026 aged 33–87 with a true MMSE of ≥ 24 (64,615 people) was measured in 2026 and, if alive, again five years later with a new error. No visits were missed and no one dropped out, so only death removed people. If dementia emerged between the visits the individual was retained. The second measurement carried the practice gain of a retest after five years (0.10 SD for everyone). For the practice-adjusted estimate it was repeated without that gain and with the same error, so the gain was removed exactly rather than estimated. Thus, this simulates the practice effect and is not evidence for it. People were grouped by age at entry into 5-year groups. The cross-sectional estimate is each group’s mean measured memory at entry less that of the group centered on 35. The longitudinal estimates sum, from 35, the mean five-year changes of successive groups among those measured twice (the value at 45 is the sum of the changes of the groups centered on 35 and 40), and the true change (dashed) is chained in the same way from the same people’s true values.

### Figure 4C

The population was everyone alive and free of dementia in 2026 aged 80–89 (4,587 people). The SuperAger episodic memory threshold was the mean of dementia-free 56- to 66-year-olds (14,385 people; +0.06 SD), and the limits for executive function and semantic memory were 1 SD (population SD) below the mean of dementia-free people in the same 5-year age group (80–84 or 85–89). Applied to true values, the rule identified 713 SuperAgers. Labels applied the same cutoffs to one measurement of each of the three tests. The second assessment measured all three again in the same year, with new errors and, unlike the application’s own retest, no practice gain, and applied the rule again. To make the visualization clearer, the left plot shows every second person (2,294).

### Figure 4D

Each of the 100 trials drew 1,200 participants (720 women and 480 men) from the 4,801 people aged 65–85 in 2026 with amyloid PET of 33 Centiloid (a load of 0.24) or more and a CDR Sum of Boxes of 0 (the application’s A4 strategy, with its amyloid cutoff raised from 20 Centiloid to approximate A4’s enrollment cutoff; see Combining measures across studies), and randomized half of them to treatment, with the allocation seed equal to the selection seed. Treatment cleared 30% of plaque from the year after entry, which evokes the assumptions of the expanded AD cascade (version 100; Supplementary Figure 3) set so that plaque drives all of AD tau’s rise and AD’s yearly toll follows that rise (plaque drive 100%; weights 0, 1 and 0 for tau’s level, tau’s rise and amyloid). Untreated lives are then exactly those of the default world, and treated lives lose 30% less to AD each year, in the brain and cognition and so in dementia and survival. The cognitive toll of aging, age-related tau, the cardiovascular chain and LATE was not changed. Episodic memory was measured at entry and every 6 months, with error and with practice that accumulated with each test (0.23 SD at 6 months, 0.41 SD by 2 years). At 2, 4, 6, 8 and 10 years, each participant’s least-squares slope of measured memory on time was fitted to all visits made while alive (those who died before 6 months had a single visit and no slope; about 1,180 per trial were analyzed), and the difference in mean slope, treated minus control, was tested by Welch’s t (two-sided). The true effect is the mean, over participants, of each one’s slope of true memory with treatment minus the slope without it. Figure 4D plots each trial’s estimate, and the true effect, multiplied by trial length, and above each length the power to detect the true effect, Φ(μ/σ − 1.96), where Φ is the standard normal distribution function, μ the true effect and σ the standard deviation of the 100 trials’ estimates. The trials differ only in who was drawn and how they were allocated, since each person’s life and errors are fixed. Note that here and elsewhere, changes in assumptions will change the patterns. For example, in the default settings practice effects did not diminish with dementia progression, which could be implemented and enhance the detectability of the intervention effect. In cognitively unimpaired people, practice effects are smaller with greater amyloid and tau burden, and diminished practice effects predict cognitive decline (Jutten et al., 2022; Young et al., 2023; Papp et al., 2024; Kim et al., 2026). The simulated trials also measured episodic memory alone, whereas A4’s primary outcome, the Preclinical Alzheimer Cognitive Composite, combines episodic memory with timed executive function and global cognition (Donohue et al., 2014).

### Figure 4E

The definitions were applied to everyone free of dementia in 2026 aged 70–90 whose MMSE, measured once, was 24 or more (14,405 of 14,822 people). Norms came from the same lives 20 years earlier: the 81,583 people aged 18 or older and free of dementia in 2006, each test measured once with errors of the default sizes drawn by the analysis script (seed 12345). The memory cutoffs were the mean at 50–65 (H), the mean at 30–40 (D), the same-sex mean at 18–32 (S) and the same-sex median at 20–29 (M), and B’s limit for processing speed was 1.25 SD below the mean at 20–29; H also required age 80 or older. A domain was no more than 1 SD below the mean for age if its score was at least the mean less 1 SD of the same 5-year age group in 2006; for people 80 or older, that group was the 80-year-olds, the oldest alive in 2006. These domains were executive function and semantic memory for H, processing speed and executive function for D, and executive function for S. For M, memory and the MMSE (at least 29) were measured again 2 and 4 years later and had to meet the criteria at all three tests; for B, processing speed was measured again 2.5 years later, and its decline, the first score less the second, had to be below 0.5 SD. Retests carried new errors and the application’s practice gains, and people who died before a retest could not qualify by M or B. Shares are of all 14,405 people. The overlap plot illustrates, for each combination of definitions that yielded 45 or more people, the share of the 5,598 people classified by exactly that combination of definitions.

### Figure 4F

The three worlds simulated the same 120,000 lives (world seed 7). The world without gains set the cohort gain beyond schooling to zero and removed schooling’s effect on cognition. The default world adds 0.13 SD per decade of birth (relative to 1950) to episodic memory, processing speed and executive function at 40, beyond schooling’s 0.07 SD per year of schooling, which rises across birth years, so that each later decade of birth is about 0.17 SD higher at 60. The third world set the gain beyond schooling at 0.10 SD a decade, to approximate the gain at 60 reported by Fjell et al. (2026), and multiplied the rates of normal aging (not each person’s own drift or terminal decline) and the toll of AD, LATE and white matter lesions on cognition by 0.5 for each decade of birth after 1935, counting births before 1930 as 1930 and after 1960 as 1960 (factors 1.41 to 0.18). The curves use each life’s true episodic memory at every age reached, including after 2026. A cohort’s change from age 35 sums, year by year, the mean change over the preceding year of age among people born within 2 years of its birth year alive at that age. The cross-sectional estimate at each age is the mean true memory of everyone alive in 2026 within a year of that age (3-year age groups), including people with dementia. The dashed line chains, in the same way as Figure 4B, the mean change over the following year of age of people born within 2 years of those of each age in 2026. The application counted the reality checks within range in each world.

### Figure 5

The populations were everyone alive in 2026 aged 70–74, 75–79, 80–84 and 85–89 (5,592, 4,881, 3,496 and 2,096 people, including those with dementia). Samples of 400 were drawn as above from those meeting each criterion in true values: the application’s Random, Nondemented (MMSE ≥ 24) and MCI (CDR Sum of Boxes 0.5–2.5 and MMSE ≥ 24) strategies at their defaults. Random selected from everyone, and Biomarker negative was the Nondemented strategy keeping only people with amyloid PET below 12 Centiloid and tau PET below 1.25 SUVR. SuperAgers, without dementia, were selected following the rule of Harrison et al. (2012) and Rogalski et al. (2013) relaxed to include ages 70-79, with the limits of dementia-free people in each 5-year age group, and drawn as the application draws its SuperAger samples (240 women and 160 men). Where fewer women or men qualified than their quota, all who qualified were taken; at 85–89, only 191 people qualified as SuperAgers and 338 as biomarker negative, so every sample of either held the same people. Each strategy was drawn 100 times, and panels A–E show the first draw at 80–84. Hippocampal volume, episodic memory (expressed in SD of the true values of dementia-free 50- to 65-year-olds in 2026, from their mean) and intracranial volume were each measured once, so a person had the same measured values in every sample. Within each sample, hippocampal volume was regressed by least squares on intracranial volume and sex, and memory on sex alone, and r is the correlation of the two sets of residuals. Plotted points are the residuals plus the sample’s means, so the least-squares line through them has the adjusted slope; the reference line uses everyone’s coefficients, set to the sample’s mean intracranial volume and share of women. Panel F gives the median of each strategy’s 100 correlations and their 2.5th and 97.5th percentiles.

### Figure 6

Each variant changed one setting of the original model (gray). More AD risk raised the setting for amyloid positivity by age 70 from 27% to 45% (realized among the living at 70, 32% to 49%). A faster cascade halved every lag in the AD cascade and doubled tau’s rise; stronger LATE raised the setting for LATE by 85 from 12% to 30% (realized, 15% to 34%) and doubled its effects on cognition; and late-rising aging had aging’s rates at 70 double every 6 years. In panel C, people free of pathology had amyloid PET below 20 Centiloid (a load of 0.03) and tau PET below 1.30 SUVR, had not reached the onsets of LATE and white matter lesions, and were without dementia. In panel D, the course of dementia is chained from each year’s mean change in MMSE among those alive.

### Fresh-agent tests

Each test used a separate AI agent (Claude Opus 5.5, Anthropic) running in its own folder that contained only the version 78 file. Each agent received one request, phrased as a researcher would paste it into a new conversation, which it ran with a headless browser and Node.js, and was told not to read anything outside its folder. Each AI agent reported its start and end times, how it tested its work, the reality checks and a critique of the guide. Effort was set to maximum and speed to fast mode (2.5× acceleration). The second request set each person’s smoking exposure to the average of the smoking curve over the years in which they were 18 to 40 and multiplied the hazard of vascular risk by 2 raised to that exposure; because exposure is set by birth year and ranges from 10% to 43%, the factor ranges from 1.07 to 1.35.

### Use of AI in developing the simulator

An AI agent, Claude Opus 5.5 (Anthropic), wrote all of the simulator’s code; no code was written or edited by hand. The agent worked in Anthropic’s chat application with tools to run code (Node.js, Python and a headless Chromium browser). Versions 1 to 99 were built in a main design conversation, with side conversations that built experimental stages or checked the model against new values obtained from reference papers; version 99 is a single HTML file of 8,316 lines. Later versions, including version 102 (Supplementary Figure 6), were built the same way. RLB stated each change in plain language: what to model, each assumption and the published values that anchor it, the reality checks and their ranges, and the application’s layout and wording. He supplied data tables, supplementary figures, and original code where available to check against (e.g., the Josefsson maintainer rule implementation was checked against raw original code).

Embedded in the HTML is a record of the 100+ revisions and modifications, and how they arose. The agent turned each request into code, prototyped calculations in Node.js, ran the whole application in a headless browser, compared every reality check and stage with the previous version, wrote the documentation inside the file (the developer guide, How the model works, the version history and Insights and puzzles), and proposed calibrations and alternatives, which were applied only when RLB chose them. RLB corrected the agent when it overreached. Side conversations returned specifications that RLB carried into the main one: version 99’s five fixes (to APOE ε4’s risk, amyloid’s late rise, age-related tau, plasma pTau217 and AD-related hippocampal atrophy), for example, came from a check of version 96 against 64 recent biomarker papers, in which twelve parallel instances compared the papers with the model and a separate instance fact-checked the agent’s report before RLB chose which changes to make. In all, over 400 prompts spread across distinct chats and sometimes run in parallel were utilized to develop and refine the simulator. Example prompts and the AI agent’s interpretation are summarized in Supplementary Figure 8.

### Use of AI in writing this paper

The paper was written with the same model, in a separate conversation that began from the first author’s argument and abstract and version 78 of the simulator. The agent generated each figure by running the simulator (the engine in Node.js 22 and the application in headless Chromium through Playwright), drew initial figures, and drafted explanatory text, which was then heavily edited and revised by the authors. As an example, Figure 2 went through more than 30 successive versions. The agent proposed a general initial design on request, for which each life was plotted from age 40 and split into its level and each causal chain’s contribution. Among the many iterations, 12 substantial revisions were directed by authors, including the idea for the per-group pie charts, the selection by different criteria, the row showing level by change which was specifically motivated by discussions among multiple authors, and the row showing the effect of measurement error. In terms of the text evolution, by late versions of the paper, less than 20% of it had word strings of 6 or more words that matched the original AI-generated main text, with figure legends being the highest at nearly a third. The authors reviewed and approved every passage. Each time the paper moved to a newer simulator version, the agent re-ran the affected analyses from the new file (Figure 6 was kept at version 88), dividing the larger rebuilds among 10 or more parallel instances, and updated the numbers in the text. At major revisions, including each move to a newer version, a separate instance that had not produced the work recomputed each number cited in the changed passages from the saved data and checked each statement against the figures and code, and the discrepancies it found were corrected before the version was returned and then the resolutions reviewed.

### Responsible use of AI

The authors chose the questions, the assumptions and their published anchors, the reality checks, the analyses and the interpretations, and they take responsibility for the content. No participant data were used beyond published values and replots of published data. The option to allow Anthropic to use the code and conversations to improve its models was turned off.

### Availability

The simulator file, copies of historical versions, and the Markdown descriptions for each Figure panel are available in the GitHub repository (https://buckner-lab.github.io/Aging-Brain-Simulator).

## Acknowledgments.

Funding was generously provided by the National Institute on Aging (R01AG067420 and U24AG092760), the Simons Foundation, and the Vranos Family Fund. LN was supported by the Knut and Alice Wallenberg (KAW) Foundation.

## Footnotes

1 Aging, as discussed under Limitations and Future Directions, is a placeholder for processes affecting aging that are not otherwise labeled or understood.

## References

1. Andersen, K., Lolk, A., Martinussen, T., and Kragh-Sørensen, P. (2010). Very mild to severe dementia and mortality: a 14-year follow-up – the Odense study. Dement. Geriatr. Cogn. Disord. 29, 61–67. 10.1159/000265553

2. Arenaza-Urquijo, E.M., and Vemuri, P. (2018). Resistance vs resilience to Alzheimer disease: clarifying terminology for preclinical studies. Neurology 90, 695–703. 10.1212/WNL.0000000000005303

3. Banack, H.R., Kaufman, J.S., Wactawski-Wende, J., Troen, B.R., and Stovitz, S.D. (2019). Investigating and remediating selection bias in geriatrics research: the selection bias toolkit. J. Am. Geriatr. Soc. 67, 1970–1976. 10.1111/jgs.16022

4. Barnett, A.G., van der Pols, J.C., and Dobson, A.J. (2005). Regression to the mean: what it is and how to deal with it. Int. J. Epidemiol. 34, 215–220. 10.1093/ije/dyh299

5. Benkarim, O., Paquola, C., Park, B.-y., Kebets, V., Hong, S.-J., Vos de Wael, R., et al. (2022). Population heterogeneity in clinical cohorts affects the predictive accuracy of brain imaging. PLoS Biol. 20, e3001627. 10.1371/journal.pbio.3001627

6. Bott, N.T., Bettcher, B.M., Yokoyama, J.S., Frazier, D.T., Wynn, M., Karydas, A., et al. (2017). Youthful processing speed in older adults: genetic, biological, and behavioral predictors of cognitive processing speed trajectories in aging. Front. Aging Neurosci. 9, 55. 10.3389/fnagi.2017.00055

7. Box, G.E.P. (1976). Science and statistics. J. Am. Stat. Assoc. 71, 791–799. 10.1080/01621459.1976.10480949

8. Boyle, P.A., Wang, T., Yu, L., Wilson, R.S., Dawe, R., Arfanakis, K., et al. (2021). To what degree is late life cognitive decline driven by age-related neuropathologies? Brain 144, 2166–2175. 10.1093/brain/awab092

9. Boyle, P.A., Wilson, R.S., Yu, L., Barr, A.M., Honer, W.G., Schneider, J.A., et al. (2013). Much of late life cognitive decline is not due to common neurodegenerative pathologies. Ann. Neurol. 74, 478–489. 10.1002/ana.23964

10. Buckley, R.F., Townsend, D.L., Birkenbihl, C.J., Cuppels, M., Coughlan, G.T., Seto, M.T., et al. (2026). Prognostic value of blood-based p-tau217 levels for progression to cognitive impairment. JAMA 336, 950–959. 10.1001/jama.2026.12556

11. Buckner, R.L. (2004). Memory and executive function in aging and AD: multiple factors that cause decline and reserve factors that compensate. Neuron 44, 195–208. 10.1016/j.neuron.2004.09.006

12. Busche, M.A., and Hyman, B.T. (2020). Synergy between amyloid-β and tau in Alzheimer’s disease. Nat. Neurosci. 23, 1183–1193. 10.1038/s41593-020-0687-6

13. Chen, M., Tworek, J., Jun, H., Yuan, Q., Pinto, H.P.d.O., Kaplan, J., et al. (2021). Evaluating large language models trained on code. Preprint at arXiv. 10.48550/arXiv.2107.03374

14. Corrada, M.M., Brookmeyer, R., Paganini-Hill, A., Berlau, D., and Kawas, C.H. (2010). Dementia incidence continues to increase with age in the oldest old: the 90+ study. Ann. Neurol. 67, 114–121. 10.1002/ana.21915

15. Cronbach, L.J., and Furby, L. (1970). How we should measure “change”: or should we? Psychol. Bull. 74, 68–80. 10.1037/h0029382

16. Dang, C., Harrington, K.D., Lim, Y.Y., Ames, D., Hassenstab, J., Laws, S.M., et al. (2019). Superior memory reduces 8-year risk of mild cognitive impairment and dementia but not amyloid β-associated cognitive decline in older adults. Arch. Clin. Neuropsychol. 34, 585–598. 10.1093/arclin/acy078

17. Donohue, M.C., Sperling, R.A., Salmon, D.P., Rentz, D.M., Raman, R., Thomas, R.G., et al. (2014). The preclinical Alzheimer cognitive composite: measuring amyloid-related decline. JAMA Neurol. 71, 961–970. 10.1001/jamaneurol.2014.803

18. Epstein, J.M. (2008). Why model? J. Artif. Soc. Soc. Simul. 11, 12. https://www.jasss.org/11/4/12.html

19. Falk, E.B., Hyde, L.W., Mitchell, C., Faul, J., Gonzalez, R., Heitzeg, M.M., et al. (2013). What is a representative brain? Neuroscience meets population science. Proc. Natl. Acad. Sci. USA 110, 17615–17622. 10.1073/pnas.1310134110

20. Fjell, A.M., Grødem, E.O.S., Lunansky, G., Vidal-Piñeiro, D., Rogeberg, O.J., and Walhovd, K.B. (2026). Generational gains in memory capacity and stability may account for declining dementia incidence rates in Europe and the United States. Preprint at medRxiv. 10.64898/2026.04.14.26350835

21. Folstein, M.F., Folstein, S.E., and McHugh, P.R. (1975). “Mini-mental state”: a practical method for grading the cognitive state of patients for the clinician. J. Psychiatr. Res. 12, 189–198. 10.1016/0022-3956(75)90026-6

22. Fraser, M.A., Shaw, M.E., and Cherbuin, N. (2015). A systematic review and meta-analysis of longitudinal hippocampal atrophy in healthy human ageing. Neuroimage 112, 364–374. 10.1016/j.neuroimage.2015.03.035

23. Goldberg, T.E., Harvey, P.D., Wesnes, K.A., Snyder, P.J., and Schneider, L.S. (2015). Practice effects due to serial cognitive assessment: implications for preclinical Alzheimer’s disease randomized controlled trials. Alzheimers Dement. (Amst.) 1, 103–111. 10.1016/j.dadm.2014.11.003

24. Gorbach, T., Pudas, S., Bartrés-Faz, D., Brandmaier, A.M., Düzel, S., Henson, R.N., et al. (2020). Longitudinal association between hippocampus atrophy and episodic-memory decline in non-demented APOE ε4 carriers. Alzheimers Dement. (Amst.) 12, e12110. 10.1002/dad2.12110

25. Gow, A.J., Johnson, W., Pattie, A., Brett, C.E., Roberts, B., Starr, J.M., et al. (2011). Stability and change in intelligence from age 11 to ages 70, 79, and 87: the Lothian Birth Cohorts of 1921 and 1936. Psychol. Aging 26, 232–240. 10.1037/a0021072

26. Grødem, E.O.S., Vidal-Piñeiro, D., Sørensen, Ø., Bartrés-Faz, D., Brandmaier, A.M., Cattaneo, G., et al. (2026). Stable individual differences dominate adult brain volume variation until later life. Imaging Neurosci. 4, IMAG.a.1242. 10.1162/IMAG.a.1242

27. Guest, O., and Martin, A.E. (2021). How computational modeling can force theory building in psychological science. Perspect. Psychol. Sci. 16, 789–802. 10.1177/1745691620970585

28. Hansson, O. (2021). Biomarkers for neurodegenerative diseases. Nat. Med. 27, 954–963. 10.1038/s41591-021-01382-x

29. Harrison, T.M., Weintraub, S., Mesulam, M.M., and Rogalski, E. (2012). Superior memory and higher cortical volumes in unusually successful cognitive aging. J. Int. Neuropsychol. Soc. 18, 1081–1085. 10.1017/S1355617712000847

30. Hayes-Larson, E., Mobley, T.M., Mungas, D., Seamans, M.J., Glymour, M.M., Gilsanz, P., et al. (2022). Accounting for lack of representation in dementia research: generalizing KHANDLE study findings on the prevalence of cognitive impairment to the California older population. Alzheimers Dement. 18, 2209–2217. 10.1002/alz.12522

31. Hedden, T., and Gabrieli, J.D.E. (2004). Insights into the ageing mind: a view from cognitive neuroscience. Nat. Rev. Neurosci. 5, 87–96. 10.1038/nrn1323

32. Hofer, S.M., and Sliwinski, M.J. (2001). Understanding ageing: an evaluation of research designs for assessing the interdependence of ageing-related changes. Gerontology 47, 341–352. 10.1159/000052825

33. Hoffman, L., Hofer, S.M., and Sliwinski, M.J. (2011). On the confounds among retest gains and age-cohort differences in the estimation of within-person change in longitudinal studies: a simulation study. Psychol. Aging 26, 778–791. 10.1037/a0023910

34. Ikram, M.A. (2024). The use and misuse of ‘biological aging’ in health research. Nat. Med. 30, 3045. 10.1038/s41591-024-03297-9

35. Jack, C.R., Jr., Andrews, J.S., Beach, T.G., Buracchio, T., Dunn, B., Graf, A., et al. (2024). Revised criteria for diagnosis and staging of Alzheimer’s disease: Alzheimer’s Association Workgroup. Alzheimers Dement. 20, 5143–5169. 10.1002/alz.13859

36. Jack, C.R., Jr., Knopman, D.S., Jagust, W.J., Petersen, R.C., Weiner, M.W., Aisen, P.S., et al. (2013). Tracking pathophysiological processes in Alzheimer’s disease: an updated hypothetical model of dynamic biomarkers. Lancet Neurol. 12, 207–216. 10.1016/S1474-4422(12)70291-0

37. Jack, C.R., Jr., Knopman, D.S., Jagust, W.J., Shaw, L.M., Aisen, P.S., Weiner, M.W., et al. (2010). Hypothetical model of dynamic biomarkers of the Alzheimer’s pathological cascade. Lancet Neurol. 9, 119–128. 10.1016/S1474-4422(09)70299-6

38. Jack, C.R., Jr., Wiste, H.J., Weigand, S.D., Therneau, T.M., Lowe, V.J., Knopman, D.S., et al. (2017). Defining imaging biomarker cut points for brain aging and Alzheimer’s disease. Alzheimers Dement. 13, 205–216. 10.1016/j.jalz.2016.08.005

39. Jansen, W.J., Ossenkoppele, R., Knol, D.L., Tijms, B.M., Scheltens, P., Verhey, F.R.J., et al. (2015). Prevalence of cerebral amyloid pathology in persons without dementia: a meta-analysis. JAMA 313, 1924–1938. 10.1001/jama.2015.4668

40. Jedynak, B.M., Lang, A., Liu, B., Katz, E., Zhang, Y., Wyman, B.T., et al. (2012). A computational neurodegenerative disease progression score: method and results with the Alzheimer’s Disease Neuroimaging Initiative cohort. Neuroimage 63, 1478–1486. 10.1016/j.neuroimage.2012.07.059

41. Jorm, A.F., and Jolley, D. (1998). The incidence of dementia: a meta-analysis. Neurology 51, 728– 733. 10.1212/WNL.51.3.728

42. Josefsson, M., de Luna, X., Pudas, S., Nilsson, L.-G., and Nyberg, L. (2012). Genetic and lifestyle predictors of 15-year longitudinal change in episodic memory. J. Am. Geriatr. Soc. 60, 2308–2312. 10.1111/jgs.12000

43. Josefsson, M., Sundström, A., Pudas, S., Nordin Adolfsson, A., Nyberg, L., and Adolfsson, R. (2023). Memory profiles predict dementia over 23-28 years in normal but not successful aging. Int. Psychogeriatr. 35, 351–359. 10.1017/S1041610219001844

44. Jutten, R.J., Rentz, D.M., Fu, J.F., Mayblyum, D.V., Amariglio, R.E., Buckley, R.F., et al. (2022). Monthly at-home computerized cognitive testing to detect diminished practice effects in preclinical Alzheimer’s disease. Front. Aging Neurosci. 13, 800126. 10.3389/fnagi.2021.800126

45. Kapasi, A., Yu, L., Boyle, P.A., Barnes, L.L., Bennett, D.A., and Schneider, J.A. (2020). Limbic-predominant age-related TDP-43 encephalopathy, ADNC pathology, and cognitive decline in aging. Neurology 95, e1951–e1962. 10.1212/WNL.0000000000010454

46. Katzman, R., Terry, R., DeTeresa, R., Brown, T., Davies, P., Fuld, P., et al. (1988). Clinical, pathological, and neurochemical changes in dementia: a subgroup with preserved mental status and numerous neocortical plaques. Ann. Neurol. 23, 138–144. 10.1002/ana.410230206

47. Kawas, C.H., Kim, R.C., Sonnen, J.A., Bullain, S.S., Trieu, T., and Corrada, M.M. (2015). Multiple pathologies are common and related to dementia in the oldest-old: the 90+ Study. Neurology 85, 535–542. 10.1212/WNL.0000000000001831

48. Kim, H., Soberanes, D., Jutten, R.J., Dubbelman, M.A., Yang, H.-S., Marshall, G.A., et al. (2026). Combining p-tau217 and digital cognitive testing to predict cognitive decline. Alzheimers Dement. 22, e71328. 10.1002/alz.71328

49. Klunk, W.E., Koeppe, R.A., Price, J.C., Benzinger, T.L., Devous, M.D., Jagust, W.J., et al. (2015). The Centiloid Project: standardizing quantitative amyloid plaque estimation by PET. Alzheimers Dement. 11, 1–15.e4. 10.1016/j.jalz.2014.07.003

50. Knuth, D.E. (1984). Literate programming. Comput. J. 27, 97–111. 10.1093/comjnl/27.2.97

51. Kuhn, T.S. (1962). The Structure of Scientific Revolutions (University of Chicago Press).

52. LeWinn, K.Z., Sheridan, M.A., Keyes, K.M., Hamilton, A., and McLaughlin, K.A. (2017). Sample composition alters associations between age and brain structure. Nat. Commun. 8, 874. 10.1038/s41467-017-00908-7

53. Lindenberger, U., Singer, T., and Baltes, P.B. (2002). Longitudinal selectivity in aging populations: separating mortality-associated versus experimental components in the Berlin Aging Study (BASE). J. Gerontol. B Psychol. Sci. Soc. Sci. 57, P474–P482. 10.1093/geronb/57.6.P474

54. Lindenberger, U., von Oertzen, T., Ghisletta, P., and Hertzog, C. (2011). Cross-sectional age variance extraction: what’s change got to do with it? Psychol. Aging 26, 34–47. 10.1037/a0020525

55. Livingston, G., Huntley, J., Liu, K.Y., Costafreda, S.G., Selbæk, G., Alladi, S., et al. (2024). Dementia prevention, intervention, and care: 2024 report of the Lancet standing Commission. Lancet 404, 572–628. 10.1016/S0140-6736(24)01296-0

56. Lövdén, M., Fratiglioni, L., Glymour, M.M., Lindenberger, U., and Tucker-Drob, E.M. (2020). Education and cognitive functioning across the life span. Psychol. Sci. Public Interest 21, 6–41. 10.1177/1529100620920576

57. Maccora, J., Peters, R., and Anstey, K.J. (2021). Gender differences in superior-memory SuperAgers and associated factors in an Australian cohort. J. Appl. Gerontol. 40, 433–442. 10.1177/0733464820902943

58. Manly, J.J., Jones, R.N., Langa, K.M., Ryan, L.H., Levine, D.A., McCammon, R., et al. (2022). Estimating the prevalence of dementia and mild cognitive impairment in the US: the 2016 Health and Retirement Study Harmonized Cognitive Assessment Protocol Project. JAMA Neurol. 79, 1242– 1249. 10.1001/jamaneurol.2022.3543

59. Mayeda, E.R., Filshtein, T.J., Tripodis, Y., Glymour, M.M., and Gross, A.L. (2018). Does selective survival before study enrolment attenuate estimated effects of education on rate of cognitive decline in older adults? A simulation approach for quantifying survival bias in life course epidemiology. Int. J. Epidemiol. 47, 1507–1517. 10.1093/ije/dyy124

60. Mayeda, E.R., Tchetgen, E.J., Power, M.C., Weuve, J., Jacqmin-Gadda, H., Marden, J.R., et al. (2016). A simulation platform for quantifying survival bias: an application to research on determinants of cognitive decline. Am. J. Epidemiol. 184, 378–387. 10.1093/aje/kwv451

61. Merenstein, J.L., and Bennett, I.J. (2022). Bridging patterns of neurocognitive aging across the older adult lifespan. Neurosci. Biobehav. Rev. 135, 104594. 10.1016/j.neubiorev.2022.104594

62. Meyer, D.H., and Schumacher, B. (2024). Aging clocks based on accumulating stochastic variation. Nat. Aging 4, 871–885. 10.1038/s43587-024-00619-x

63. Muthukrishna, M., and Henrich, J. (2019). A problem in theory. Nat. Hum. Behav. 3, 221–229. 10.1038/s41562-018-0522-1

64. National Academies of Sciences, Engineering, and Medicine (2024). Foundational Research Gaps and Future Directions for Digital Twins (The National Academies Press). 10.17226/26894

65. Nelson, P.T., Dickson, D.W., Trojanowski, J.Q., Jack, C.R., Boyle, P.A., Arfanakis, K., et al. (2019). Limbic-predominant age-related TDP-43 encephalopathy (LATE): consensus working group report. Brain 142, 1503–1527. 10.1093/brain/awz099

66. Nilsson, L.-G., Sternäng, O., Rönnlund, M., and Nyberg, L. (2009). Challenging the notion of an early-onset of cognitive decline. Neurobiol. Aging 30, 521–524. 10.1016/j.neurobiolaging.2008.11.013

67. Nyberg, L., Boraxbekk, C.-J., Sörman, D.E., Hansson, P., Herlitz, A., Kauppi, K., et al. (2020). Biological and environmental predictors of heterogeneity in neurocognitive ageing: evidence from Betula and other longitudinal studies. Ageing Res. Rev. 64, 101184. 10.1016/j.arr.2020.101184

68. Nyberg, L., Lövdén, M., Riklund, K., Lindenberger, U., and Bäckman, L. (2012). Memory aging and brain maintenance. Trends Cogn. Sci. 16, 292–305. 10.1016/j.tics.2012.04.005

69. Nyberg, L., Magnussen, F., Lundquist, A., Baaré, W., Bartrés-Faz, D., Bertram, L., et al. (2021). Educational attainment does not influence brain aging. Proc. Natl. Acad. Sci. USA 118, e2101644118. 10.1073/pnas.2101644118

70. Papp, K.V., Jutten, R.J., Soberanes, D., Weizenbaum, E., Hsieh, S., Molinare, C., et al. (2024). Early detection of amyloid-related changes in memory among cognitively unimpaired older adults with daily digital testing. Ann. Neurol. 95, 507–517. 10.1002/ana.26833

71. Persson, J., Nyberg, L., Lind, J., Larsson, A., Nilsson, L.-G., Ingvar, M., et al. (2006). Structure-function correlates of cognitive decline in aging. Cereb. Cortex 16, 907–915. 10.1093/cercor/bhj036

72. Petersen, R.C., Aisen, P.S., Beckett, L.A., Donohue, M.C., Gamst, A.C., Harvey, D.J., et al. (2010). Alzheimer’s Disease Neuroimaging Initiative (ADNI): clinical characterization. Neurology 74, 201–209. 10.1212/WNL.0b013e3181cb3e25

73. Platt, J.R. (1964). Strong inference. Science 146, 347–353. 10.1126/science.146.3642.347

74. Powell, A., Lam, B.C.P., Foxe, D., Close, J.C.T., Sachdev, P.S., and Brodaty, H. (2024). Frequency of cognitive “super-aging” in three Australian samples using different diagnostic criteria. Int. Psychogeriatr. 36, 939–955. 10.1017/S1041610223000935

75. Raz, N., and Lindenberger, U. (2011). Only time will tell: cross-sectional studies offer no solution to the age–brain–cognition triangle: comment on Salthouse (2011). Psychol. Bull. 137, 790–795. 10.1037/a0024503

76. Raz, N., Lindenberger, U., Rodrigue, K.M., Kennedy, K.M., Head, D., Williamson, A., et al. (2005). Regional brain changes in aging healthy adults: general trends, individual differences and modifiers. Cereb. Cortex 15, 1676–1689. 10.1093/cercor/bhi044

77. Ritchie, S.J., and Tucker-Drob, E.M. (2018). How much does education improve intelligence? A meta-analysis. Psychol. Sci. 29, 1358–1369. 10.1177/0956797618774253

78. Roberts, R.O., Aakre, J.A., Kremers, W.K., Vassilaki, M., Knopman, D.S., Mielke, M.M., et al. (2018). Prevalence and outcomes of amyloid positivity among persons without dementia in a longitudinal, population-based setting. JAMA Neurol. 75, 970–979. 10.1001/jamaneurol.2018.0629

79. Rogalski, E.J., Gefen, T., Shi, J., Samimi, M., Bigio, E., Weintraub, S., et al. (2013). Youthful memory capacity in old brains: anatomic and genetic clues from the Northwestern SuperAging Project. J. Cogn. Neurosci. 25, 29–36. 10.1162/jocn_a_00300

80. Rönnlund, M., and Nilsson, L.-G. (2008). The magnitude, generality, and determinants of Flynn effects on forms of declarative memory and visuospatial ability: time-sequential analyses of data from a Swedish cohort study. Intelligence 36, 192–209. 10.1016/j.intell.2007.05.002

81. Rönnlund, M., Nyberg, L., Bäckman, L., and Nilsson, L.-G. (2005). Stability, growth, and decline in adult life span development of declarative memory: cross-sectional and longitudinal data from a population-based study. Psychol. Aging 20, 3–18. 10.1037/0882-7974.20.1.3

82. Salthouse, T.A. (2009). When does age-related cognitive decline begin? Neurobiol. Aging 30, 507–514. 10.1016/j.neurobiolaging.2008.09.023

83. Salthouse, T.A. (2010). Influence of age on practice effects in longitudinal neurocognitive change. Neuropsychology 24, 563–572. 10.1037/a0019026

84. Salthouse, T.A. (2019). Trajectories of normal cognitive aging. Psychol. Aging 34, 17–24. 10.1037/pag0000288

85. Savva, G.M., Wharton, S.B., Ince, P.G., Forster, G., Matthews, F.E., and Brayne, C.; Medical Research Council Cognitive Function and Ageing Study (2009). Age, neuropathology, and dementia. N. Engl. J. Med. 360, 2302–2309. 10.1056/NEJMoa0806142

86. Schaie, K.W. (1994). The course of adult intellectual development. Am. Psychol. 49, 304–313. 10.1037/0003-066X.49.4.304

87. Schaie, K.W. (2009). “When does age-related cognitive decline begin?” Salthouse again reifies the “cross-sectional fallacy”. Neurobiol. Aging 30, 528–529. 10.1016/j.neurobiolaging.2008.12.012

88. Shaw, C., Hayes-Larson, E., Glymour, M.M., Dufouil, C., Hohman, T.J., Whitmer, R.A., et al. (2021). Evaluation of selective survival and sex/gender differences in dementia incidence using a simulation model. JAMA Netw. Open 4, e211001. 10.1001/jamanetworkopen.2021.1001

89. Singh-Manoux, A., Kivimäki, M., Glymour, M.M., Elbaz, A., Berr, C., Ebmeier, K.P., et al. (2012). Timing of onset of cognitive decline: results from Whitehall II prospective cohort study. BMJ 344, d7622. 10.1136/bmj.d7622

90. Sliwinski, M., Lipton, R.B., Buschke, H., and Stewart, W. (1996). The effects of preclinical dementia on estimates of normal cognitive functioning in aging. J. Gerontol. B Psychol. Sci. Soc. Sci. 51B, P217–P225. 10.1093/geronb/51B.4.P217

91. Smaldino, P.E. (2017). Models are stupid, and we need more of them. In Computational Social Psychology, R.R. Vallacher, S.J. Read, and A. Nowak, eds. (Routledge), pp. 311–331. 10.4324/9781315173726-14

92. Small, S.A., Schobel, S.A., Buxton, R.B., Witter, M.P., and Barnes, C.A. (2011). A pathophysiological framework of hippocampal dysfunction in ageing and disease. Nat. Rev. Neurosci. 12, 585–601. 10.1038/nrn3085

93. Social Security Administration (2026). Actuarial life table: period life table, 2023, as used in the 2026 Trustees Report. https://www.ssa.gov/oact/STATS/table4c6.html (accessed October 10, 2026).

94. Sperling, R.A., Donohue, M.C., Raman, R., Rafii, M.S., Johnson, K., Masters, C.L., et al. (2023). Trial of solanezumab in preclinical Alzheimer’s disease. N. Engl. J. Med. 389, 1096–1107. 10.1056/NEJMoa2305032

95. Sperling, R.A., Donohue, M.C., Rissman, R.A., Johnson, K.A., Rentz, D.M., Grill, J.D., et al. (2024). Amyloid and tau prediction of cognitive and functional decline in unimpaired older individuals: longitudinal data from the A4 and LEARN studies. J. Prev. Alzheimers Dis. 11, 802–813. 10.14283/jpad.2024.122

96. Sperling, R.A., Rentz, D.M., Johnson, K.A., Karlawish, J., Donohue, M., Salmon, D.P., et al. (2014). The A4 study: stopping AD before symptoms begin? Sci. Transl. Med. 6, 228fs13. 10.1126/scitranslmed.3007941

97. Stern, Y. (2012). Cognitive reserve in ageing and Alzheimer’s disease. Lancet Neurol. 11, 1006–1012. 10.1016/S1474-4422(12)70191-6

98. Stern, Y., Arenaza-Urquijo, E.M., Bartrés-Faz, D., Belleville, S., Cantilon, M., Chetelat, G., et al. (2020). Whitepaper: defining and investigating cognitive reserve, brain reserve, and brain maintenance. Alzheimers Dement. 16, 1305–1311. 10.1016/j.jalz.2018.07.219

99. Sudlow, C., Gallacher, J., Allen, N., Beral, V., Burton, P., Danesh, J., et al. (2015). UK Biobank: an open access resource for identifying the causes of a wide range of complex diseases of middle and old age. PLoS Med. 12, e1001779. 10.1371/journal.pmed.1001779

100. Sun, F.W., Stepanovic, M.R., Andreano, J., Barrett, L.F., Touroutoglou, A., and Dickerson, B.C. (2016). Youthful brains in older adults: preserved neuroanatomy in the default mode and salience networks contributes to youthful memory in superaging. J. Neurosci. 36, 9659–9668. 10.1523/JNEUROSCI.1492-16.2016

101. Tschanz, J.T., Corcoran, C.D., Schwartz, S., Treiber, K., Green, R.C., Norton, M.C., et al. (2011). Progression of cognitive, functional, and neuropsychiatric symptom domains in a population cohort with Alzheimer dementia: the Cache County Dementia Progression study. Am. J. Geriatr. Psychiatry 19, 532–542. 10.1097/JGP.0b013e3181faec23

102. Tucker-Drob, E.M., Brandmaier, A.M., and Lindenberger, U. (2019). Coupled cognitive changes in adulthood: a meta-analysis. Psychol. Bull. 145, 273–301. 10.1037/bul0000179

103. van Alten, S., Domingue, B.W., Faul, J., Galama, T., and Marees, A.T. (2024). Reweighting UK Biobank corrects for pervasive selection bias due to volunteering. Int. J. Epidemiol. 53, dyae054. 10.1093/ije/dyae054

104. van Dyck, C.H., Swanson, C.J., Aisen, P., Bateman, R.J., Chen, C., Gee, M., et al. (2023). Lecanemab in early Alzheimer’s disease. N. Engl. J. Med. 388, 9–21. 10.1056/NEJMoa2212948

105. Viceconti, M., Henney, A., and Morley-Fletcher, E. (2016). In silico clinical trials: how computer simulation will transform the biomedical industry. Int. J. Clin. Trials 3, 37–46. 10.18203/2349-3259.ijct20161408

106. Vidal-Piñeiro, D., Sørensen, Ø., Strømstad, M., Amlien, I.K., Baaré, W., Bartrés-Faz, D., et al. (2025). Vulnerability to memory decline in aging revealed by a mega-analysis of structural brain change. Nat. Commun. 16, 11488. 10.1038/s41467-025-66354-y

107. Wang, J., Buto, P., Ferguson, E.L., Chen, R., Pederson, A., Choi, M., et al. (2026). Can the All of Us sample be reweighted to mirror a nationally representative sample? A comparison of mortality predictors. Epidemiology 37, 854–866. 10.1097/EDE.0000000000002016

108. Willis, S.L., and Schaie, K.W. (2005). Cognitive trajectories in midlife and cognitive functioning in old age. In Middle Adulthood: A Lifespan Perspective, S.L. Willis and M. Martin, eds. (Sage), pp. 243–276. 10.4135/9781452225999.n8

109. Wilson, G., Aruliah, D.A., Brown, C.T., Chue Hong, N.P., Davis, M., Guy, R.T., et al. (2014). Best practices for scientific computing. PLoS Biol. 12, e1001745. 10.1371/journal.pbio.1001745

110. Wilson, R.S., Yu, L., Leurgans, S.E., Bennett, D.A., and Boyle, P.A. (2020). Proportion of cognitive loss attributable to terminal decline. Neurology 94, e42–e50. 10.1212/WNL.0000000000008671

111. Yarkoni, T. (2022). The generalizability crisis. Behav. Brain Sci. 45, e1. 10.1017/S0140525X20001685

112. Young, A.L., Marinescu, R.V., Oxtoby, N.P., Bocchetta, M., Yong, K., Firth, N.C., et al. (2018). Uncovering the heterogeneity and temporal complexity of neurodegenerative diseases with Subtype and Stage Inference. Nat. Commun. 9, 4273. 10.1038/s41467-018-05892-0

113. Young, C.B., Mormino, E.C., Poston, K.L., Johnson, K.A., Rentz, D.M., Sperling, R.A., et al. (2023). Computerized cognitive practice effects in relation to amyloid and tau in preclinical Alzheimer’s disease: results from a multi-site cohort. Alzheimers Dement. (Amst.) 15, e12414. 10.1002/dad2.12414

