## Supplemental Figures and Methods for "An Open, Evolving Brain and Cognitive Aging Simulator Enabled by Agentic Artificial Intelligence"



**An agentic AI simulator was built to advance our understanding about how the human brain and cognition change with age. The simulator generates worlds based on hypothesized age-linked cascades and then simulates studies across varied experimental designs and selection criteria. Each simulated world begins with 120,000 individual lives initiated by baseline parameters (e.g., genetic risk, birth year) and then changed by age-dependent cascades (e.g., amyloid-dependent, cardiovascular). Examples illustrate how the simulator predicts distinct data patterns in cross-sectional versus longitudinal designs, how alternative participant selection strategies emphasize baseline differences versus age-related decline, and how variable levels of association emerge between brain and cognition dependent upon design choices. Simulations of clinical trials provide information about design, power, and outcome considerations. Experiments on alternative simulated worlds evaluate how well hypothesized cascades accommodate real-world data patterns, capturing the insights and open puzzles inside the application. Most critically, the application is built to evolve. Its code contains a complete description of how to construct and update itself so that any person or AI agent can explore a new finding, recalibrate to emerging data, or build and simulate a novel future study. The simulator is offered as a tool for understanding data patterns, teaching, and generation of hypotheses to be tested in future studies using real-world data collection.**

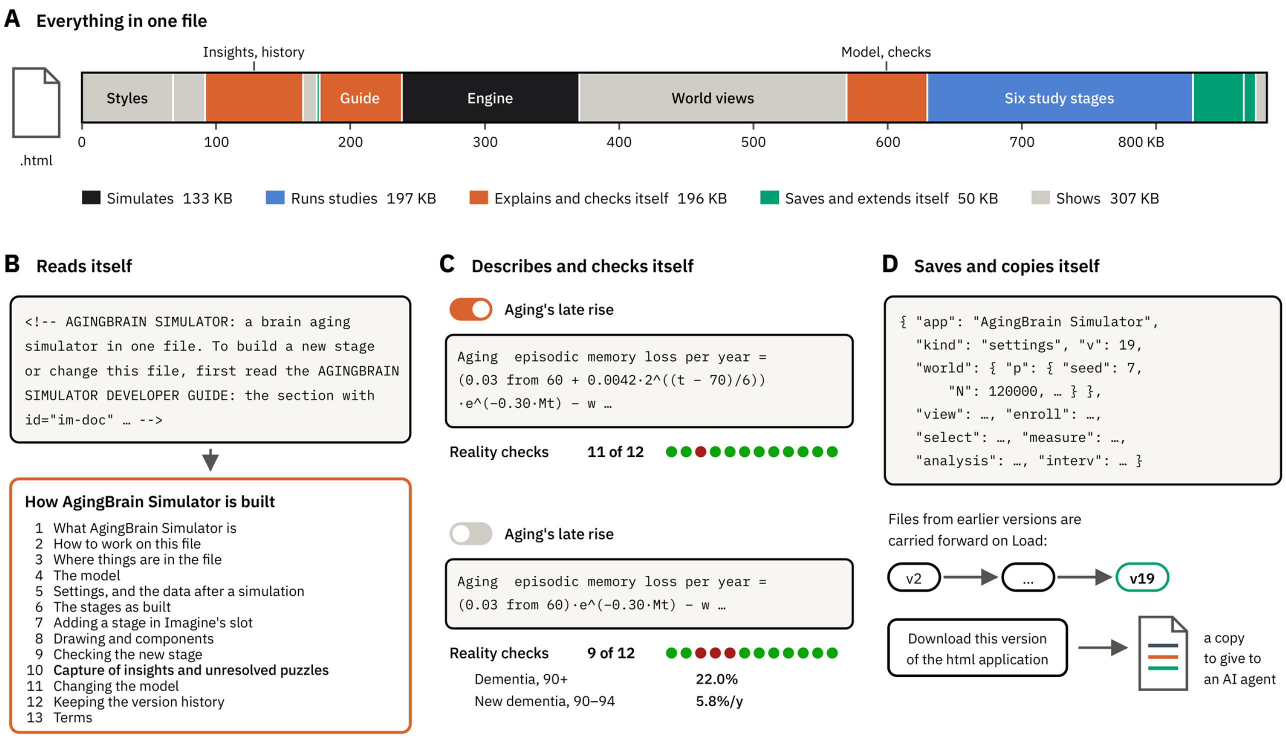


**Supplementary Figure 1. The simulator is one HTML file that runs as an application and contains instructions to rebuild itself.** (**A**) The HTML file contents are shown in order, colored by their functional role in the application: simulating lives (the engine, black), running study stages (blue), explaining and checking the model (orange: the developer guide, the model description, the reality checks, the insights and the version history), saving and extending the application (green: Save and Load, Imagine), and displaying the world (gray). (**B**) The HTML file's first lines direct readers, which could be a person or AI agent, to the embedded developer guide where section 10 sets the rules used by the application to capture and store insights that emerge as new findings are incorporated and stages are built. (**C**) The model's description and checks are generated from the current settings. As an example, turning aging's late rise on allows 11 of the 12 checks to pass. When it is switched off, the description changes as two further checks fail, providing information that late-rising aging, or an alternative mechanism, is required to generate data patterns that are consistent with the reality checks. (**D**) Each setting is saved as a versioned file (excerpt shown) that can be loaded and exchanged between users. The “Download this version of the html application” feature saves a copy of the full self-contained application which contains its code, instructions on how to modify, and the history of modifications to that point. The next developer (human or AI agent) begins new modifications from the same complete account of the simulation and its history at that point.

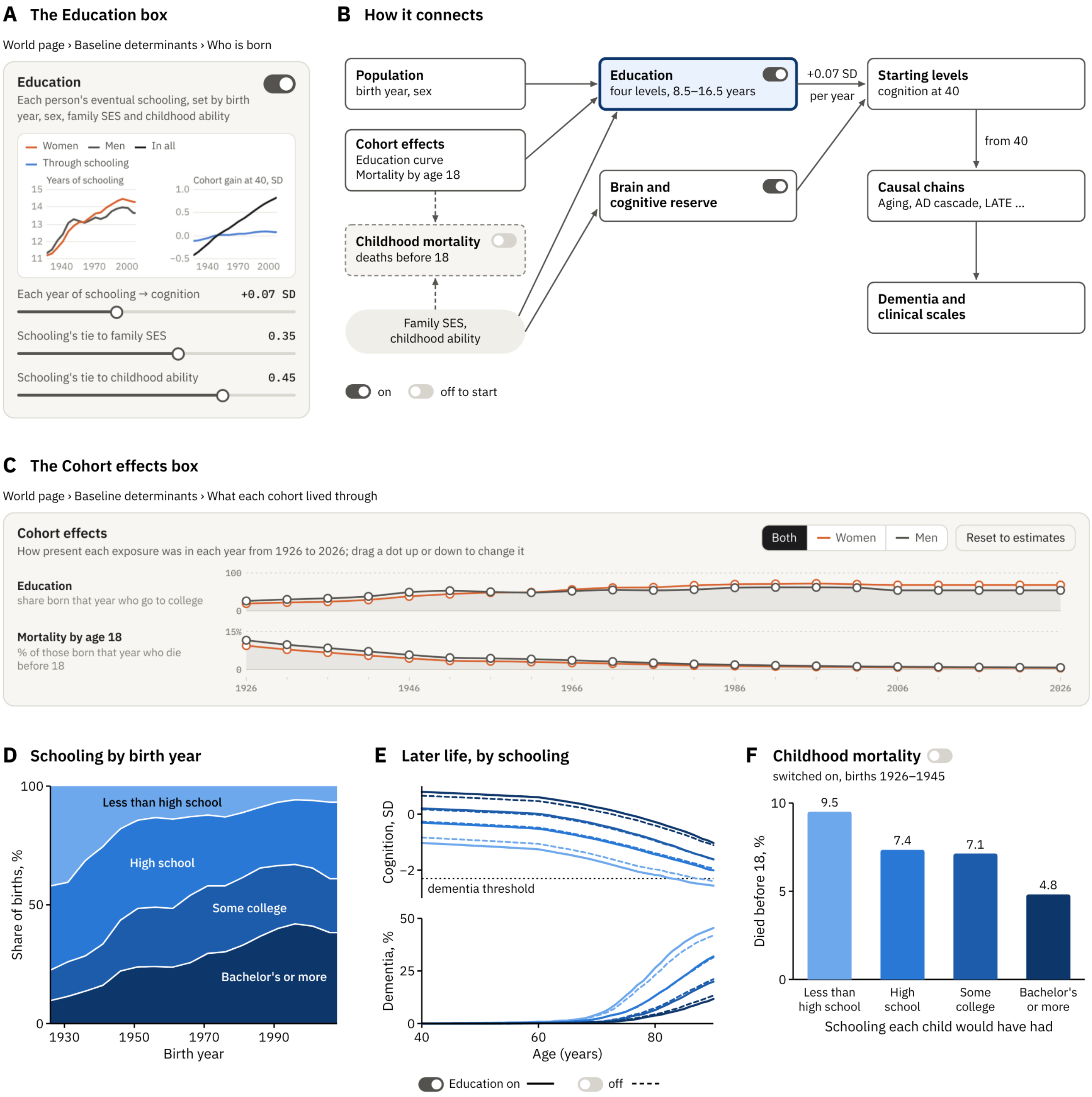
A The Education box
World page › Baseline determinants › Who is born
Education
Each person's eventual schooling, set by birth year, sex, family SES and childhood ability
Women
Men
In all
Through schooling
Years of schooling
Cohort gain at 40, SD
Each year of schooling → cognition +0.07 SD
Schooling's tie to family SES 0.35
Schooling's tie to childhood ability 0.45
B How it connects
Population
birth year, sex
Education
four levels, 8.5–16.5 years
+0.07 SD
per year
Starting levels
cognition at 40
Cohort effects
Education curve
Mortality by age 18
Brain and cognitive reserve
from 40
Causal chains
Aging, AD cascade, LATE ...
Childhood mortality
deaths before 18
Family SES,
childhood ability
Dementia and
clinical scales
on
off to start
C The Cohort effects box
World page › Baseline determinants › What each cohort lived through
Cohort effects
How present each exposure was in each year from 1926 to 2026; drag a dot up or down to change it
Both
Women
Men
Reset to estimates
Education
share born that year who go to college
Mortality by age 18
% of those born that year who die before 18
D Schooling by birth year
Less than high school
High school
Some college
Bachelor's or more
Share of births, %
Birth year
E Later life, by schooling
Cognition, SD
dementia threshold
Dementia, %
Age (years)
Education on
off
F Childhood mortality
switched on, births 1926–1945
Died before 18, %
9.5
7.4
7.1
4.8
Less than high school
High school
Some college
Bachelor's or more
Schooling each child would have had

**Supplementary Figure 2. Baseline determinants set the level and rates from which each life unfolds.** The simulator builds a world of people who are each born with individual abilities and propensities for how their brain and cognitive faculties will change with aging. The beginning parameters are set by the 11 modules under Baseline determinants on the World page. Modules can be switched on or off, and each has settings that determine values used by other modules to simulate a life. Education is shown as an example Baseline determinant, which itself interacts with the Cohort effects to simulate each person's life. When that within-person microsimulation is repeated 120,000 times, an overall population emerges that is influenced by all of the Baseline determinants and their settings. (**A**) The Education box is shown as it appears in the application (when switched on) displaying its charts and settings. (**B**) A causal diagram illustrates how Education is set and what it influences. Each person's schooling comes from a propensity, 0.35·SES + 0.45·ability + 0.72·e (SD 1), that sets one of four levels: less than high school, high school, some college, and a bachelor's degree or more (corresponding roughly to 8.5, 12, 14 and 16.5 years). The share who goes to college comes from the Education curve in Cohort effects. The shares of simulated lives who finish a bachelor's degree and who leave without a high school diploma follow US Census Bureau estimates (US Census Bureau, 2026). Each year of schooling above 13.0 adds 0.07 SD to episodic memory, processing speed and executive function to people's starting levels at age 40. From there, causal chains (e.g., Aging, AD cascade, LATE) impact cognition which, in turn, impacts dementia (Dementia and the CDR-SB). Family SES and childhood ability also act on cognition directly (Brain and cognitive reserve). With Education on, part of their effect runs through schooling. Childhood mortality, off to start, draws on the same two traits and on the Mortality by age 18 Cohort effect. (**C**) The Cohort effects box is illustrated with two of its ten curves shown separately for women (orange) and men (gray). Education (top) is the share of each birth year who went to college and Mortality (bottom) is the percentage of lives that die by age 18. Eight other Cohort effects are not illustrated. Within each Cohort effect, dragging a dot changes a curve's values allowing users to explore the impacts of alternative settings and how they shape the simulated population that emerges. (**D**) The shares of the four levels of Schooling are shown as a function of birth year. These shares contribute to the simulation of cohort effects. For example, people having a bachelor's degree or more rises from 10% of births in 1926 to 38% in 2008. (**E**) The effects of Schooling are shown on cognition and dementia expressed in later life as an example of how Baseline determinants shape the population of simulated lives. The lines plot 14,090 people born in 1945–1954 who are alive at 40. Mean global cognition among the living is displayed (top) and the share who have developed dementia (bottom). The dotted line shows the dementia threshold. More schooling sets the cognition level to be higher on average at 40. Cognition declines at a similar pace, but dementia is detected later because of the baseline differences at age 40. With Education switched off (dashed), schooling no longer adds to cognition and the gap narrows. The gap nonetheless remains to a lessened degree because family SES and childhood ability, which set schooling, act directly on cognition as well. This reflects another feature that exists in many of the simulator's values: each individual life is its own microsimulation with multiple determinants that act both through direct and indirect paths. (**F**) Childhood mortality. When switched on, deaths before 18 follow the Mortality by age 18 curve, scaled by ×0.87 per SD of higher family SES and ×0.81 per SD of higher childhood ability. The children who die are therefore mostly those who would have had less schooling. For example, among births in 1926–1945, 9.5% of children who would not have finished high school die before 18, against 4.8% of those who would have earned a bachelor's degree or more and, as a result, the adults who remain have slightly more schooling. Values come from the default world (seed 7, 120,000 lives). Switching a module on or off, or changing one of its settings, influences what the downstream modules produce, which is how the simulator lets a user see what each assumption contributes to the overall world's population and to the individual lives within it.

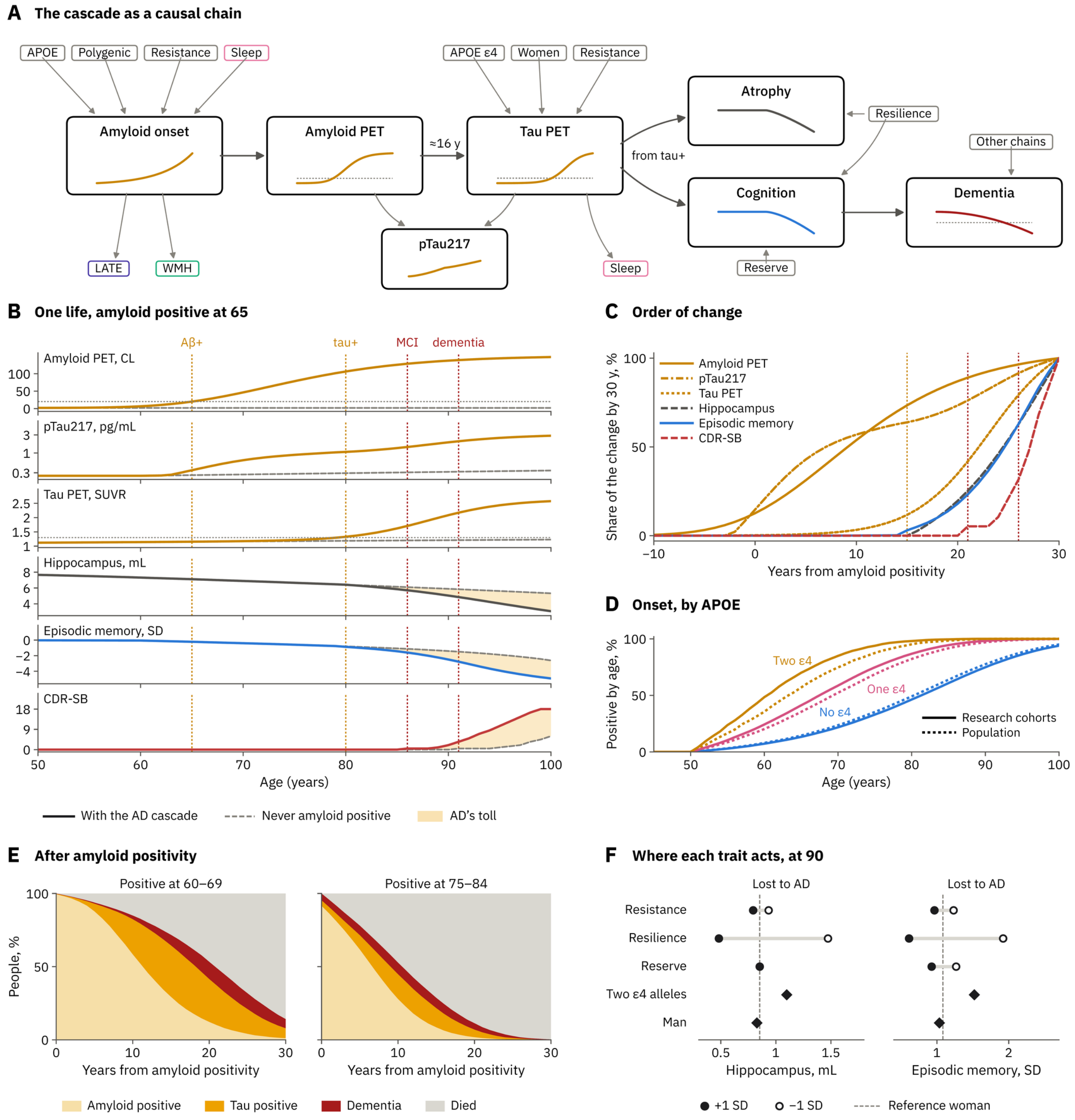
A The cascade as a causal chain
APOE
Polygenic
Resistance
Sleep
Amyloid onset
LATE
WMH
Amyloid PET
≈16 y
pTau217
APOE ε4
Women
Resistance
Tau PET
Sleep
from tau+
Atrophy
Resilience
Cognition
Reserve
Other chains
Dementia
B One life, amyloid positive at 65
Aβ+
tau+
MCI
dementia
Amyloid PET, CL
pTau217, pg/mL
Tau PET, SUVR
Hippocampus, mL
Episodic memory, SD
CDR-SB
Age (years)
With the AD cascade
Never amyloid positive
AD's toll
C Order of change
Share of the change by 30 y, %
Amyloid PET
pTau217
Tau PET
Hippocampus
Episodic memory
CDR-SB
Years from amyloid positivity
D Onset, by APOE
Positive by age, %
Two ε4
One ε4
No ε4
Research cohorts
Population
Age (years)
E After amyloid positivity
Positive at 60–69
Positive at 75–84
People, %
Years from amyloid positivity
Amyloid positive
Tau positive
Dementia
Died
F Where each trait acts, at 90
Lost to AD
Resistance
Resilience
Reserve
Two ε4 alleles
Man
Hippocampus, mL
Episodic memory, SD
+1 SD
−1 SD
Reference woman

**Supplementary Figure 3. The AD cascade is a chain of explicit, adjustable links that specify the pathway and rates from amyloid onset to dementia.** The AD cascade is one example of several included causal cascades that are all used collectively by the simulator to change brain and cognitive levels across an individual's life. These cascades are each displayed inside the simulator's Causal chains module within the World section. (**A**) The basic AD cascade, which was introduced in version 5 of the simulator, is shown as it is implemented. A separate, expanded AD cascade was introduced in version 100 as an alternative (turned off by default). The presence of two AD cascades is intended to illustrate the flexibility of the simulator to test alternative sets of assumptions, as either cascade can be used and adjusted. An entirely different cascade can also be constructed based on new findings or a different set of assumptions. In the basic AD cascade, amyloid onset (the age at which amyloid PET reaches 20 Centiloid, CL) is set as a risk curve $h(a) = h_{70} \cdot e^{(\ln 2/8)(a - 70)}$ from age 50, implemented such that 24% of APOE ε3/ε3 carriers are positive by 70 (anchored to the Mayo Clinic Study of Aging; Roberts et al., 2018) and multiplied by 3.76 with one ε4 allele and 8.93 with two (Castellano et al., 2025), 0.6 per ε2 allele (against 0.69 for ε2/ε3 in the same study), 1.25 per SD of polygenic score (without APOE), $e^{-0.4R}$ for resistance R and 1.25 per SD of a poor-sleep trait. Amyloid PET then rises along an S-curve (rate 0.13 a year, shape 0.2) about 150 CL above a baseline of 2 ± 4 CL (Bollack et al., 2024; Gantenberg et al., 2026). The ± 4 CL is the spread of true values. With the default measurement error of 4 CL (Supplementary Figure 4A), measured values spread about ± 5.7 CL, as in the amyloid-negative reference group of Bollack et al. (2.4 ± 5.7 CL). AD tau (temporal meta-ROI) follows a logistic curve to about 1.4 SUVR above the person's baseline (1.12 ± 0.03 SUVR, plus age-related tau of 0.0022 SUVR a year on average from 50, varying between people with a log SD of 0.6; Ossenkoppele et al., 2025), timed to reach positivity (1.30 SUVR) about 16 years after amyloid. Its rate, 0.25 a year, is ×1.30 per ε4 allele, ×1.08 in women and $\times e^{-0.2R}$. Plasma pTau217 (pg/mL on the scale of the ALZpath assay; Ashton et al., 2024) is affected by both: 0.25 pg/mL × 3.3 at full plaque × 2.6 at full tau, with its plaque component rising from 12 CL and leveling off as $1 - \exp(-(CL - 12)/30)$, scaled to 1 at 100 CL (Insel et al., 2025). In the basic AD cascade, the brain loses volume each year after tau becomes positive in proportion to the tau present (hippocampus 3.1% per SUVR, cortex 1.8%, whole brain 1.0%, plus small plaque terms), and cognition falls with accumulated tau-years D as $w \cdot 40(1 - e^{-D/40})$ (w = 0.30 SD per SUVR-year for episodic memory, 0.15 for executive function and 0.12 for speed). Atrophy and cognitive decline are parallel correlates of tau. Hippocampal volume does not specifically drive memory in this version of the model; the two decline together because the same cascades affect both. Dementia is diagnosed, whatever its causes, when global cognition falls to −2.3 SD after a decline of at least 1.2 SD since age 40. The basic AD cascade also affects other chains. Tau adds 0.5 SD of sleep disruption per SUVR, and amyloid onset doubles the risk of LATE and raises that of white matter lesions 1.3-fold. These are all assumptions and rates that can be changed. The expanded AD cascade allows for more complex interactions of amyloid and tau, and further allows direct effects of both on evolving phenotypes such that their rates of change can be drivers. In this manner the expanded cascade, but not the basic cascade, is able to model the effects of amyloid clearance (see the AD cascade versus the AD cascade expanded in the Causal chains module of the simulator). The implementation of a model should not be taken as an endorsement that it is correct or the preferred model, as the simulator is explicitly designed to implement and contrast alternative models. (**B**) An example reference person is plotted to illustrate how the AD cascade affects relevant biomarkers and values as her life unfolds (a woman born in 1950, APOE ε3/ε3, average traits and draws, no vascular risk or LATE). This simulated person is plotted with the AD cascade turned off and never becomes amyloid positive (dashed lines) or with the AD cascade turned on and amyloid positivity occurs at 65 (solid lines). The top three plots show the AD cascade's effects on biomarkers and the bottom three plots show their derived effects on expressed phenotypes. Yellow shading illustrates the cascade's toll on the derived phenotypes. For this example person, tau PET turns positive at 80 (age-related tau brings it a year ahead of the lag); MCI (CDR-SB 0.5) emerges at 86 and dementia emerges at 91. (**C**) The same changes replotted to align to when the simulated life became amyloid positive with all values plotting change as positive. Plasma pTau217 is halfway after 7 years and amyloid PET after 9 years (pTau217's plaque part levels off while amyloid PET keeps rising), tau PET after 22 years, hippocampus and episodic memory after 24–25 years, and CDR-SB after 27 years. Thus, plaque markers rise first, then tau, then hippocampus and memory, and clinical symptoms last, consistent with commonly referenced biomarker models (Jack et al., 2013; Hansson, 2021). Dotted lines illustrate tau positivity, MCI and dementia. Alternative models that expand to include subtle (sensitive) cognitive assessments or examine the interactions among biomarkers are possible. (**D**) Onset across the 120,000 lives of the default world. The share who would be amyloid positive by each age if they lived, by number of APOE ε4 alleles (22%, 59% and 85% by 70; median ages 81, 68 and 60). These curves ignore death; only 62% of lives turn positive before death. Solid lines use the default ε4 risks, ×3.76 with one allele and ×8.93 with two, hazard ratios estimated in research cohorts in which 36% carried ε4 (Castellano et al., 2025). Dotted lines use an alternative fitted to amyloid positivity in ε4 carriers and noncarriers of the population-based Mayo Clinic Study of Aging, in which 28% carried ε4 (Roberts et al., 2018): ×2.8 and ×5.5, with 26% of ε3/ε3 carriers positive by 70 so that overall positivity is unchanged (24%, 52% and 75% by 70; median ages 80, 69 and 63). Both are settings in the application's AD cascade, so users can choose either set, or other values, and rerun any analysis under that assumption. (**E**) Illustration of how amyloid positivity has distinct effects at different ages. What follows amyloid positivity is plotted separately for the subgroups within the simulated world who turned amyloid positive at 60–69 (20,262) and those who turned positive at 75–84 (19,666). Of those amyloid positive at 60–69, 68% become tau positive and 48% develop dementia (left panel). Of those amyloid positive at 75–84, only 42% become tau positive and 42% develop dementia; the other 58% die without dementia (right panel). (**F**) How the basic AD cascade impacts the hippocampus and episodic memory in relation to trait values set by the Baseline determinants (resistance, resilience, reserve, ε4 alleles, and sex). Each trait's effect is plotted at +1 SD (filled) or −1 SD (open) of its level. The dashed line illustrates the reference woman. Resistance slows tau, resilience protects the brain and cognition, and reserve protects cognition selectively (note that for reserve the hippocampus loses the same amount at either level but cognition differs). Every link is a setting in the simulator, so a skeptical user can change assumptions and values, and explore how those decisions change what is simulated.

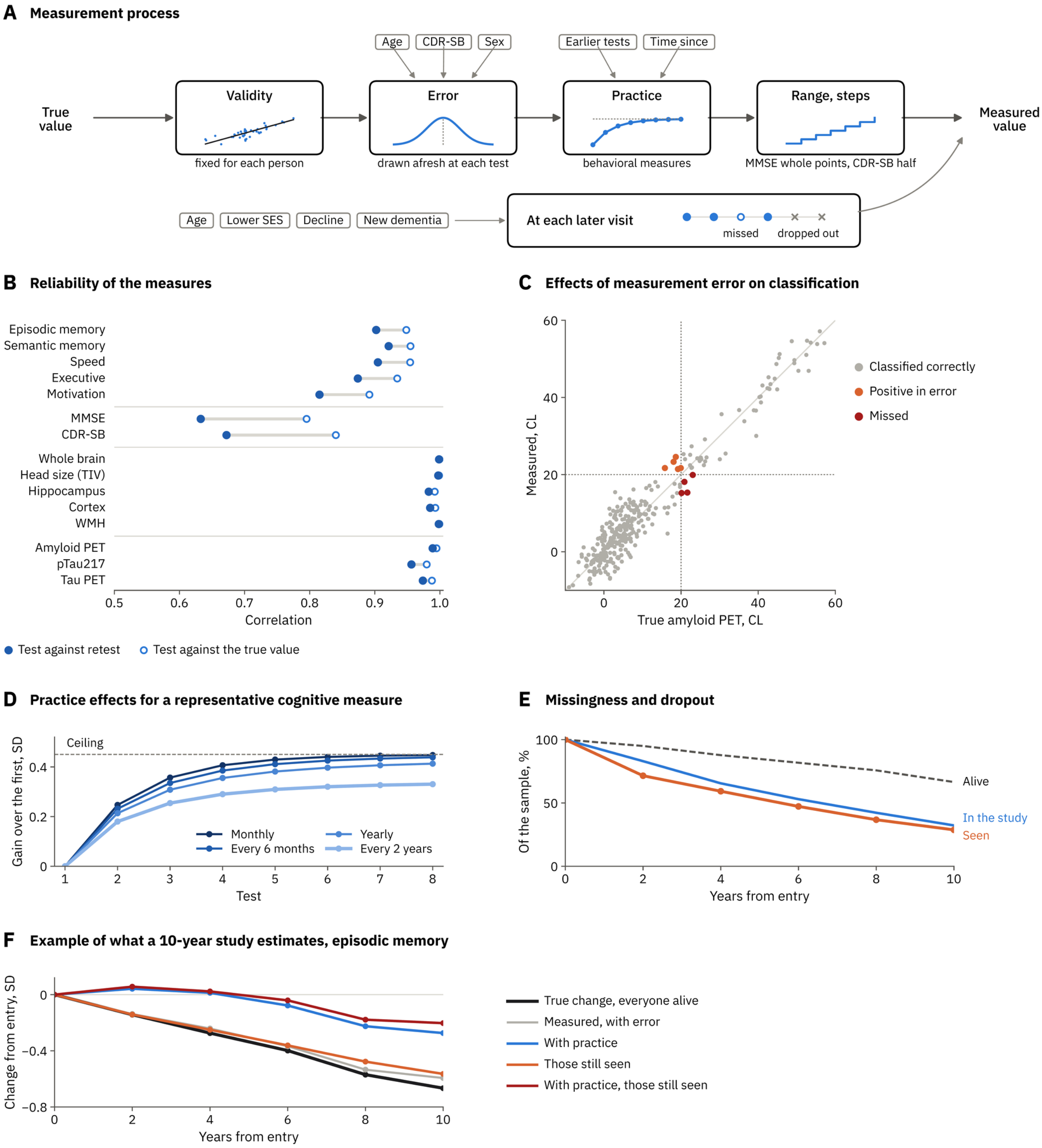

A Measurement process
Age
CDR-SB
Sex
Earlier tests
Time since
True value
Validity
fixed for each person
Error
drawn afresh at each test
Practice
behavioral measures
Range, steps
MMSE whole points, CDR-SB half
Measured value
Age
Lower SES
Decline
New dementia
At each later visit
missed
dropped out
B Reliability of the measures
Episodic memory
Semantic memory
Speed
Executive
Motivation
MMSE
CDR-SB
Whole brain
Head size (TIV)
Hippocampus
Cortex
WMH
Amyloid PET
pTau217
Tau PET
0.5
0.6
0.7
0.8
0.9
1.0
Correlation
Test against retest
Test against the true value
C Effects of measurement error on classification
Measured, CL
True amyloid PET, CL
Classified correctly
Positive in error
Missed
D Practice effects for a representative cognitive measure
Ceiling
Gain over the first, SD
Test
Monthly
Every 6 months
Yearly
Every 2 years
E Missingness and dropout
Of the sample, %
Years from entry
Alive
In the study
Seen
F Example of what a 10-year study estimates, episodic memory
Change from entry, SD
Years from entry
True change, everyone alive
Measured, with error
With practice
Those still seen
With practice, those still seen

**Supplementary Figure 4. The Measurement process turns true values into measured values impacted by error, practice effects, and attrition.** A key feature of the simulator is that it simulates lives based on how Baseline determinants and causal cascades interact, resulting in true, error-free values of the constructs across each simulated person's life. The Measurement process stage simulates how measurement instruments estimate those true values, allowing for imperfect reliability and validity as well as practice effects. (**A**) The steps of the simulated Measurement process are illustrated. A stable part carries the measure's validity r: $s = \mu + r(t - \mu) + \sqrt{(1 - r^2)}\cdot\sigma\cdot u$, where t is the true value, μ and σ the variable's mean and SD among everyone alive and u a draw fixed for each person and variable (r = 1 by default, so s = t). An error is added with SD = $e_{70} \times a^{(age - 70)/10} \times c^{CDR\text{-}SB}$, times a factor in women, where $e_{70}$ is the error at 70 (episodic memory 0.4 SD, semantic memory 0.3, speed 0.35, executive function 0.4, motivation 0.5; MMSE 1.2 points; CDR-SB 0.5; hippocampus 0.15 mL, whole brain 5 mL, head size 8 mL, cortex 0.02 mm, WMH 0.5 mL; amyloid PET 4 CL, pTau217 15% of the value (Ashton et al., 2024), tau PET 0.03 SUVR) and a, c and the women's factor are 1 by default. Behavioral measures then gain from practice, $G\cdot(1 - (1 - g_1/G)^E)$, where $g_1$ is the gain at the first retest, G the ceiling and $E = \Sigma\ 2^{-\Delta t/h}$ the exposure left by earlier tests, each attenuating with half-life h (for episodic memory 0.25 SD, 0.45 SD and 3 years). Brain and biomarker measures are set by default to have no practice effects. Values are then transformed into what the measurement instruments would be expected to provide (MMSE in whole points from 0 to 30, CDR-SB in half points to 18; volumes, tau and pTau217 kept positive). At each later visit a participant may miss the visit or leave the study, at rates for a 70-year-old of average SES multiplied per decade of age, per SD of lower SES, per SD of decline in global cognition since entry and once dementia begins after entry (none by default). Every draw is fixed by the world, the person, the variable and the visit. (**B**) Reliability in the study of Figure 1C–1H (400 people aged 60–90 in 2026 with MMSE ≥ 24), with the default errors. Each row plots the correlation coefficient of one test with a retest (filled) and with the true value (open). Measurement error least impacts values where between-person differences are large (e.g., brain and biomarker measures correlate 0.95 or more with a retest and cognitive tests 0.82–0.92), while the MMSE (0.63) and CDR-SB (0.67), which are narrow in a sample without dementia and scored in coarse steps, show lower correlation values. The cognitive test reliability values are set high by default, as occurs with composite scores, such as in the Betula study. Cognitive test measures using a single test instrument would generally have lower reliability, but the specific values would reflect the exact test being simulated. (**C**) Classification errors are illustrated at a threshold to reveal how the error in measured values can cause misclassification. Amyloid PET measured once against its true value in the same sample (the 316 people between −10 and 60 CL shown). With an error of 4 CL, 5 people are called positive in error and 4 of the 134 truly positive are missed, all with true values between 15 and 24 CL. (**D**) The practice model for episodic memory is illustrated. The expected gain over the first test at each of eight tests a month, six months, a year or two years apart. Close together, the second test gains 0.25 SD and later tests approach the 0.45 SD ceiling. When the model assumes two years apart, the second gains 0.18 SD and the eighth 0.33. Practice effects can be a major challenge to designs or, alternatively, used as a feature. For example, detection of early stages of clinical dementia might use the lack of benefit from practice effects to increase sensitivity. (**E**) Missingness and dropout, which can be set within the Measurement process stage, are plotted for illustration. 10% of visits missed and 5% leaving each year at 70, rising 1.3- and 1.5-fold per decade of age, 1.2- and 1.3-fold per SD of lower SES, 1.5- and 2-fold per SD of decline in global cognition since entry and 2- and 3-fold once dementia begins (the application's factors). The same sample visited every 2 years for 10 years. By year 10, 67% are alive, 32% are still in the study and 29% provide test values. (**F**) To illustrate the combined effects of the multiple measurement processes, change from entry is plotted over 10 years for varied scenarios that include true values among everyone alive (−0.67 SD by year 10), values measured with error (−0.59), values measured with both error and practice (−0.27), true values measured among those still participating (−0.56), and finally values measured with practice among those still participating (−0.20). Error leaves the mean unbiased on average, though in this sample it moves the 10-year change by +0.07 SD by chance; without it, practice hides all of the decline over the first 2 years and 48% over 10, attrition 15%, and the two together 61%. Because the simulation keeps the true and the measured value of every test, it can illustrate how each step of the measurement process distorts what a study measures.

### A Distributions

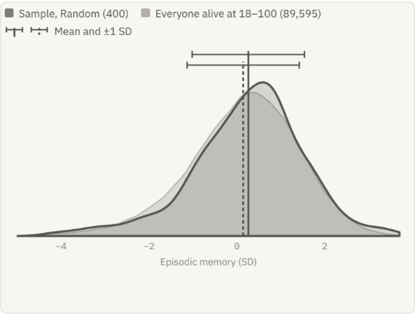


### B Power

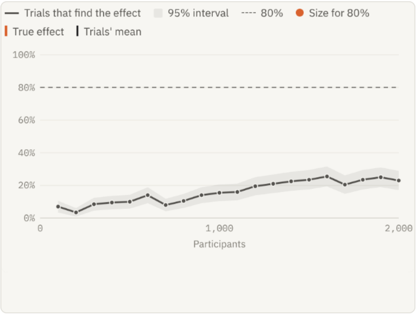


### C Biomarker timelines

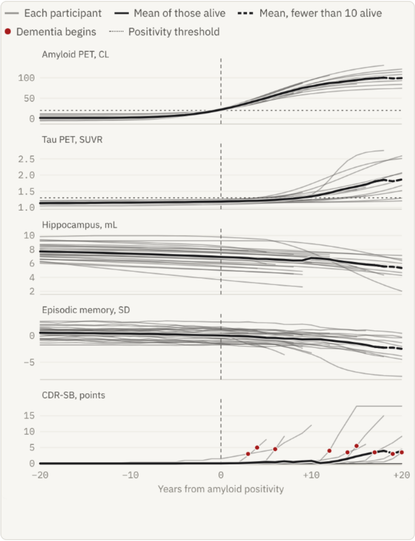


### D Compare definitions

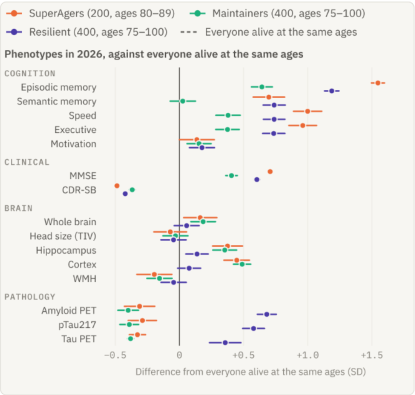


### E Export

**The files**

The participants Selection drew, as comma-separated files for a spreadsheet or a statistics program.

STUDY YEAR

`AgingBrain_world7_random_studyyear_261022.csv`

400 rows · 44 columns · 98 KB

Download study year

FOLLOW-UP VISITS

`AgingBrain_world7_random_visits_261022.csv`

1,600 rows · 37 columns · 296 KB

Download follow-up visits

DATA DICTIONARY

`AgingBrain_world7_random_dictionary_261022.csv`

81 rows · 4 columns · 9 KB

Download data dictionary

VISITS AND MEASUREMENT

| | |
|---|---|
| Visits | every 2 years for 6 years, 4 in all |
| Measured | with the error and validity set in Measurement process |
| Practice | on, as on the Follow-up page |
| Leaving | by dying only |

Set on the Follow-up page and in Measurement process

### F Life map

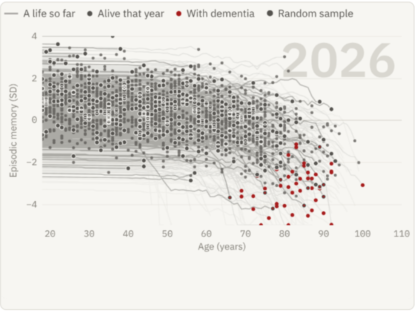


### G Generational anchors

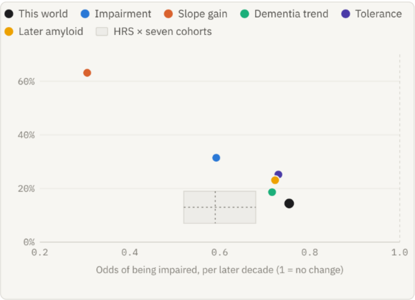


### H Alternative worlds

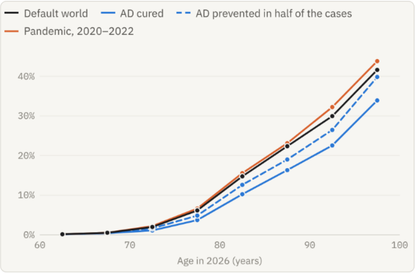


### I Diagnosis by decline

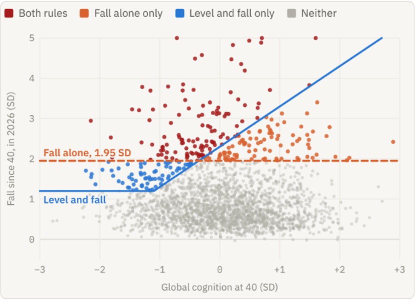


### J Childhood mortality worlds

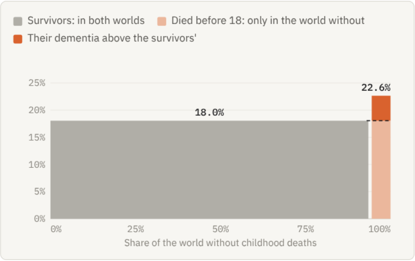

**Supplementary Figure 5. Novel stages constructed by an AI agent.** A central aspect of the AgingBrain Simulator is its ability to evolve. Illustrated are examples of novel stages that were constructed by supplying an AI agent (Claude Opus 5.5) the HTML file and one of the prompts displayed in the Imagine stage. Panels show the stages as they resulted from the example prompts. (**A**) The *Distributions* example illustrates a measured phenotype (e.g., hippocampus volume, episodic memory) in a simulated sample of people against everyone alive at the same ages, allowing noisy estimated values to be compared with true values. (**B**) The *Power* example illustrates how a random sample of people would fare in an intervention trial by repeating the intervention 200 times across a range of sample sizes. The true intervention effect is set to slow aging by 30% (0.005 SD a year). The intervention effect is detected by only 23% of trials of 2,000 participants. (**C**) The *Biomarker timelines* example aligns 30 participants on the year their amyloid turned positive to illustrate the relative timing of change in other biomarkers and cognitive levels. Tau followed amyloid positivity by a median of 9 years, and dementia by 15 years. (**D**) The *Compare definitions* example applies three different selection strategies at once to show how their application identifies subgroups with distinct phenotypic profiles: SuperAgers, maintainers, and resilient people. These subgroups exceed their typical age peers in episodic memory by 1.55, 0.64 and 1.19 SD, and 3%, 16% and 13% develop dementia within 10 years (peers, 30–35%). (**E**) The *Export* example writes the sample's values, true and measured, at entry and each visit where a measurement was made as a CSV file with a data dictionary for analysis using external tools. Such an output can be used as a teaching tool or can be used to test and validate planned analyses in a preregistration. (**F**) The Life map example follows 2,000 lives' memory through any year as they would exist in 1990 to 2040. The example image shown is static, but the actual example dynamically progresses the plots sequentially over the simulated years to reveal the impact of cohort effects and how the population of people selected by the same nominal rules changes as the decades pass. (**G**) As an example of updating the simulation to emergent findings, the *Generational anchors* example explores the implications of setting later birth cohort gains and lifelong change to the anchors of Fjell et al. (2026), recently posted on medRxiv. (**H**) The *Alternative worlds* example simulates what would happen if AD were fully cured or partially mitigated, yielding contrasting worlds. In the default world, 7.7% of people 65 and older have dementia. That value falls to 5.7% when AD is cured and to 6.8% when AD is prevented in half of those affected. That value rises to 8.1% after a pandemic that lowers the population's reserve. (**I**) The *Diagnosis by decline* example explores the use of alternative diagnostic criteria as implemented in the simulation to contrast a rule that identifies dementia by both absolute level and change over time (the simulation's default) versus an alternative rule where diagnosis is only sensitive to change. (**J**) The *Childhood mortality worlds* example compares dementia at 80–90 in two worlds that result from the same births that differ in whether there are childhood deaths. Of the 5.5% of people at 80–90 who would have died as children, 22.6% would have dementia in the world where they all survive, against 18.0% of the survivors in a world where the population is shaped by childhood mortality.

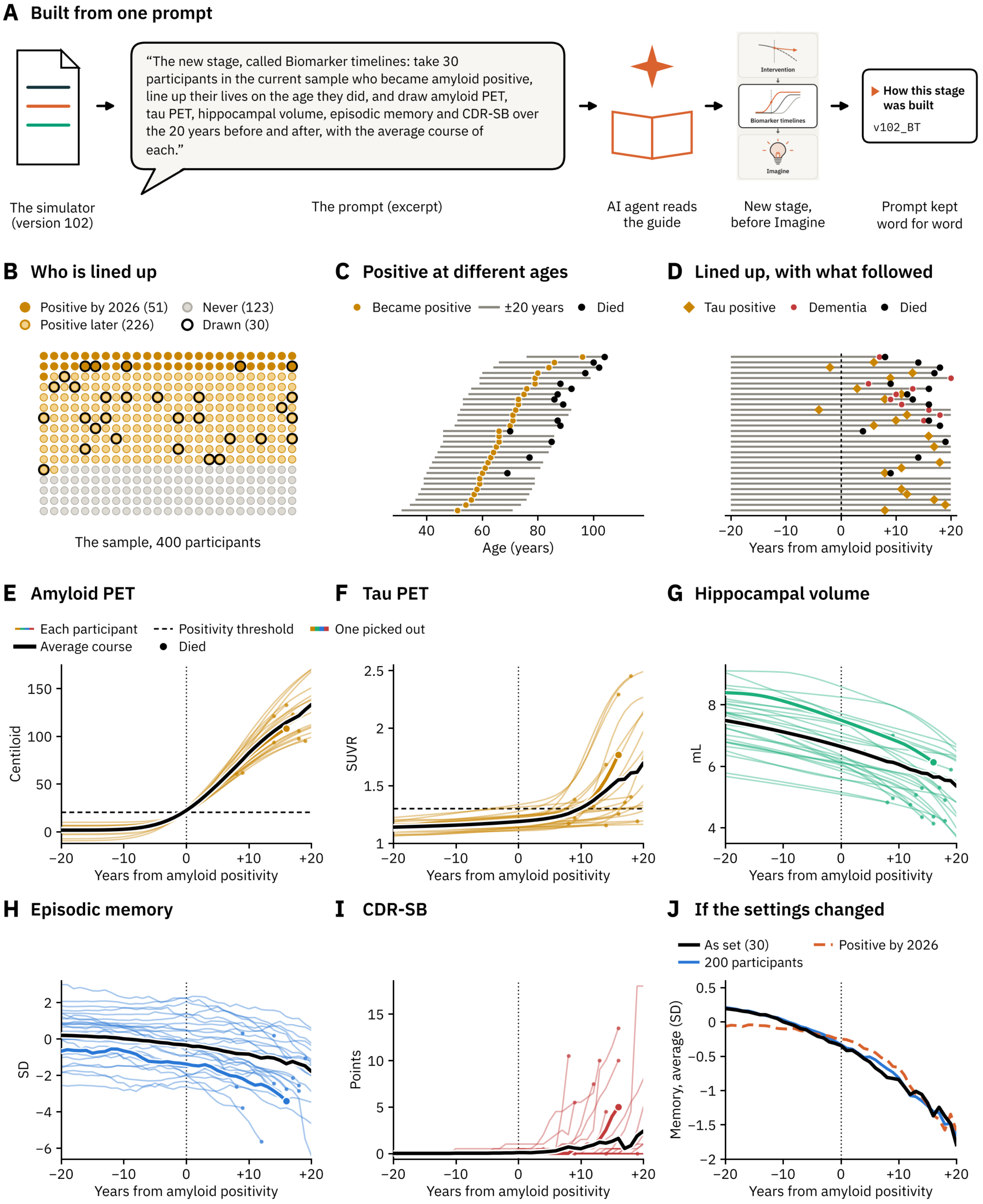
A Built from one prompt
“The new stage, called Biomarker timelines: take 30 participants in the current sample who became amyloid positive, line up their lives on the age they did, and draw amyloid PET, tau PET, hippocampal volume, episodic memory and CDR-SB over the 20 years before and after, with the average course of each.”
Intervention
Biomarker timelines
Imagine
How this stage was built
v102_BT
The simulator (version 102)
The prompt (excerpt)
AI agent reads the guide
New stage, before Imagine
Prompt kept word for word
B Who is lined up
Positive by 2026 (51)
Positive later (226)
Never (123)
Drawn (30)
The sample, 400 participants
C Positive at different ages
Became positive
±20 years
Died
Age (years)
D Lined up, with what followed
Tau positive
Dementia
Died
Years from amyloid positivity
E Amyloid PET
Each participant
Average course
Positivity threshold
Died
One picked out
Centiloid
Years from amyloid positivity
F Tau PET
SUVR
Years from amyloid positivity
G Hippocampal volume
mL
Years from amyloid positivity
H Episodic memory
SD
Years from amyloid positivity
I CDR-SB
Points
Years from amyloid positivity
J If the settings changed
As set (30)
200 participants
Positive by 2026
Memory, average (SD)
Years from amyloid positivity

**Supplementary Figure 6. An in-depth description of the new stage constructed from the Biomarker timelines example.** (**A**) The *Biomarker timelines* example is one of the ten provided examples on the Imagine stage, illustrated partially in Figures 7A and 7B of the main text and also in Supplementary Figure 5C. Here a build on version 102 is followed in detail. A fresh AI agent was provided only the HTML application file, with no access to the development history, and the Imagine stage's Copy prompt for this example: the standard opening (read the developer guide first, keep everything else as it is, record how the stage was built), and then the Biomarker timelines specific prompt (shown in full in the speech bubble). The AI agent read the guide, built the stage in 0.6 hours, and saved the new HTML version as v102_BT. Values come from the stage in the default world (seed 7, study year 2026) with Selection's random sample of 400 participants aged 18–100. (**B**) The agent read "who became amyloid positive" as amyloid PET of 20 Centiloid or more at any age, since every life is simulated to death. With this interpretation of the prompt, 277 of the simulated people qualify (51 are amyloid positive by 2026 and 226 would be positive at a later date), and 123 do not become positive before death. Of those who qualify, 30 are drawn at random (rings; 5 positive by 2026, 25 later). (**C**) The 30 people who qualify are replotted to show that the age they became positive ranged from 51 to 96 (median 66). Each person is followed over the 20 years before and after (lines) or until death (dots). (**D**) The 30 people are replotted once again lined up to the age of their amyloid positivity (year 0, the first year of age at 20 Centiloid or more) with markers showing when they became tau positive and when they became demented. By 20 years after amyloid positivity, 21 were tau positive (1.30 SUVR or more; a median of 11 years after, 2 of them 2 and 4 years before), 11 had dementia (a median of 13 years after) and 15 had died. (**E**–**I**) Five different measures are plotted for each of the 30 people aligned to when they became amyloid positive. The plots show their measures' true values without measurement error (thin; dot, death), and the average course (bold), which is the mean of those alive in each year where 5 or more people are still alive. One example individual is shown in bold so that they can be tracked across the multiple measures. This individual becomes amyloid positive at 71 (panel E), tau positive at 81 (panel F), demented at 86 (panel I), and died at 87 (where the line ends across panels). The sequence the stage shows – amyloid first, tau about a decade later and impairment in the second decade – is the simulator's AD cascade as set (Supplementary Figure 3), replotted as a study that lines up its participants on amyloid positivity. (**J**) Changing the settings changes the plots. For example, 200 participants rather than 30 (blue) barely move the average course (−1.65 SD at +20, against −1.79), while 30 participants drawn from the 51 positive people by 2026 (dashed), born a generation earlier, start lower (−0.07 SD at −20) and end at −1.59.

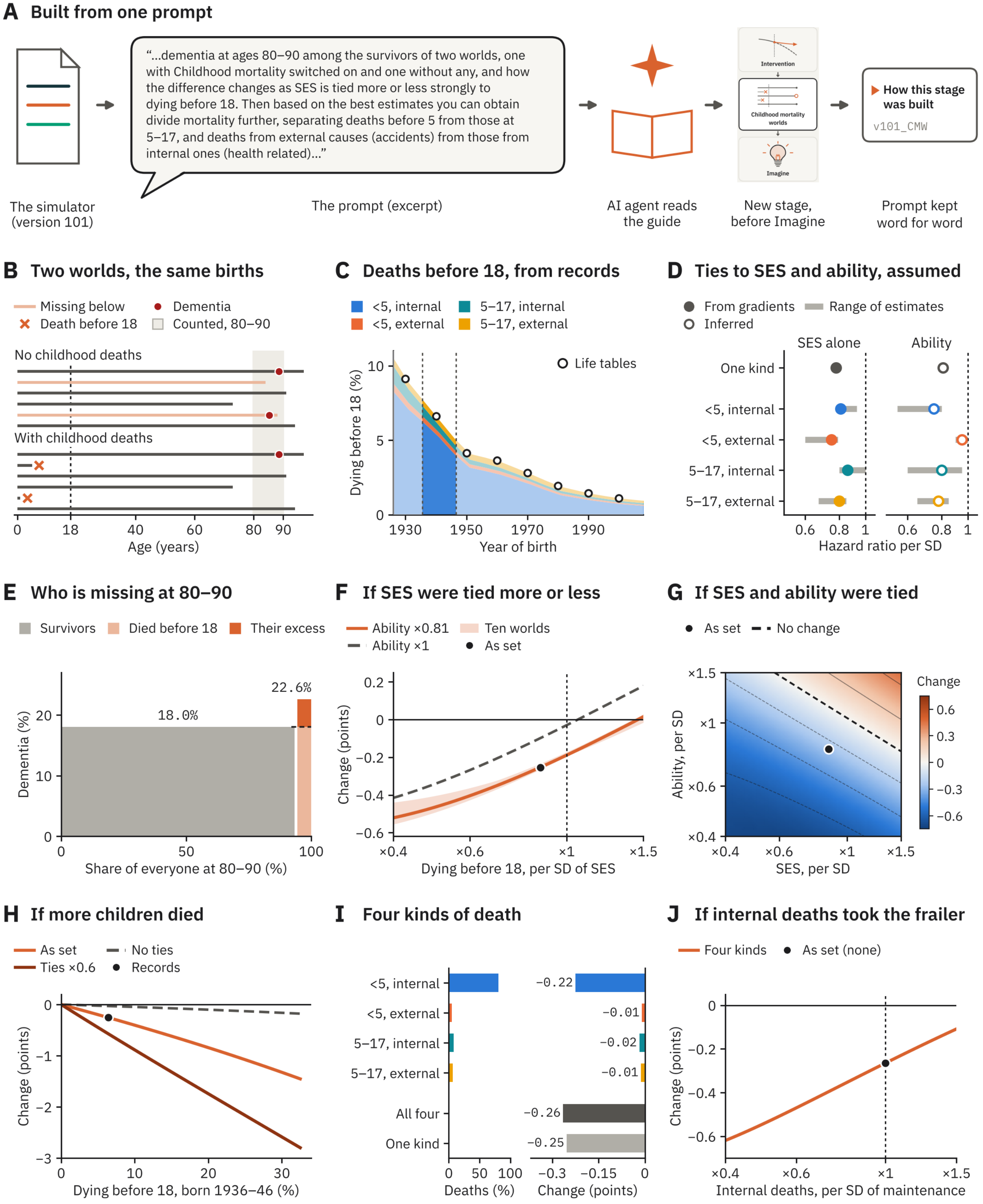
A Built from one prompt
“...dementia at ages 80–90 among the survivors of two worlds, one with Childhood mortality switched on and one without any, and how the difference changes as SES is tied more or less strongly to dying before 18. Then based on the best estimates you can obtain divide mortality further, separating deaths before 5 from those at 5–17, and deaths from external causes (accidents) from those from internal ones (health related)...”
Intervention
Childhood mortality worlds
Imagine
How this stage was built
v101_CMW
The simulator (version 101)
The prompt (excerpt)
AI agent reads the guide
New stage, before Imagine
Prompt kept word for word
B Two worlds, the same births
Missing below
Death before 18
Dementia
Counted, 80–90
No childhood deaths
With childhood deaths
Age (years)
C Deaths before 18, from records
<5, internal
<5, external
5–17, internal
5–17, external
Life tables
Dying before 18 (%)
Year of birth
D Ties to SES and ability, assumed
From gradients
Inferred
Range of estimates
SES alone
Ability
One kind
<5, internal
<5, external
5–17, internal
5–17, external
Hazard ratio per SD
E Who is missing at 80–90
Survivors
Died before 18
Their excess
18.0%
22.6%
Dementia (%)
Share of everyone at 80–90 (%)
F If SES were tied more or less
Ability ×0.81
Ability ×1
Ten worlds
As set
Change (points)
Dying before 18, per SD of SES
G If SES and ability were tied
As set
No change
Change
Ability, per SD
SES, per SD
H If more children died
As set
Ties ×0.6
No ties
Records
Change (points)
Dying before 18, born 1936–46 (%)
I Four kinds of death
<5, internal
<5, external
5–17, internal
5–17, external
All four
One kind
−0.22
−0.01
−0.02
−0.01
−0.26
−0.25
Deaths (%)
Change (points)
J If internal deaths took the frailer
Four kinds
As set (none)
Change (points)
Internal deaths, per SD of maintenance

**Supplementary Figure 7. A second in-depth description of a new stage shows how childhood deaths, set from historical records, influence dementia among the simulated survivors.** An underappreciated facet of lifespan aging is that individuals observed as older adults are survivors whose cognitive and brain levels are shaped by their early lives. As one illustration of such an effect, the present example shows how Imagine can be used to create a stage that examines the effects of Childhood mortality on late-life dementia. In this instance, the effect is simulated to be small, but the example nonetheless illustrates how the simulator can be used to model the many early life influences that shape the population. (**A**) The prompt was again the Imagine page's standard opening but here followed by a description of how the new stage should explore effects of childhood mortality (excerpt shown in speech bubble). The AI agent read the guide, searched for mortality estimates (see C below), built the stage in 0.9 hours and saved the new HTML version as v101_CMW. Values come from the stage in the default world (seed 7, study year 2026, 5,858 people aged 80–90). (**B**) Two worlds are plotted made from the same births but under two distinct survival pressures. In the top plot, children do not die and, in the bottom plot, children can die before 18 (darker lines, red X's mark deaths). In the top plot, the lives the children would have lived if there were no childhood mortality are shown as lighter lines. Dementia onset is shown as red dots. Every surviving life is the same in both, so the worlds differ only in who is missing, and dementia is estimated at 80–90 (gray). Harm to children who live through an illness, famine, or other harm is not modeled in this particular simulation, but could be in another simulation. (**C**) How many children died is set from historical records. Mortality by 18 by year of birth (sexes averaged) is estimated from the Social Security Area's cohort life tables (dots; Bell and Miller, 2005, Actuarial Study No. 120, Table 7), split by the tables' share of deaths before 5 and by external causes' share of deaths at each age (Children's Bureau, 1945, for 1939–41; National Center for Health Statistics, 1960–2019; 1950 estimated), taken at the years each cohort passed through those ages. Practical assumptions were made for implementation, such as estimating external deaths as the composite of motor-vehicle deaths plus homicide and suicide. The bright highlighted blue zone is the births of 1936–46, aged 80–90 in 2026, of whom 6.4% died before 18. (**D**) Plots show how SES and childhood ability affect the model of who died. Ties are hazard ratios per SD (below 1, fewer deaths at higher SES or ability). SES ratios (filled circles) are set from gradients of infant and child mortality by parents' schooling, income and class measured in later cohorts in the US and Europe. The ratio for SES alone, not holding ability constant, is the SES ratio times the square root of the ability ratio ($0.87 \times \sqrt{0.81} \approx 0.78$); ×0.78 alone means the lowest fifth of SES dies before 18 twice as often as the highest. Ability ratios (open circles) are inferred from IQ in youth and later deaths, the mortality of children with intellectual disability and the lower IQ of children born preterm. Bars show the approximate span of the estimates. Alternative models and values are possible. This example is meant to illustrate how such a model can be implemented, and not to advocate for the specific values set in this particular model. (**E**) As set, 5.5% of the people at 80–90 in the world without childhood deaths would have died as children (width on x axis). Of them, 22.6% would have dementia against 18.0% of the survivors, as they are lower in ability and SES. (**F**–**H**, **J**) Each sweep moves one assumption from its setting (black circles), tracing simulated consequences of the assumptions. These plots show a particularly useful feature of simulation as real data do not have these values but, within the bounds of the simulator's assumptions, insights can be gained by generating these plots as illustrations of how various factors *could* affect measured data. (**F**) The SES tie is plotted, holding ability constant, with ability's tie as set (×0.81) or with none (×1). (**G**) Both SES and ability are plotted at once (color, the change in points; dashed, no change) spanning −0.72 to +0.46 points. The change follows ability's tie more than SES's (each alone, −0.19 and −0.10 points), as ability shapes dementia more in the model (see the Brain and cognitive reserve Baseline determinant where the simulator sets Childhood ability → cognition, 0.65 SD; SES → cognition, 0.20 SD). (**H**) The level, from no deaths to six times the records, is plotted with the ties as set in the simulation, both at ×0.6, or none (−0.03 points at the records, from birth year and sex alone). The change grows nearly in proportion. (**I**) Four kinds of death are modeled and their shares of the deaths (left) and their parts of the change (right) plotted. Deaths before 5 from internal causes account for 81% of the deaths. (**J**) A final plot shows how the simulation can provide intuitions about what a hypothesized mechanism might do, here modeling what would happen if childhood mortality caused greater attrition among the frail children. To do so, internal deaths are set to differentially affect the children lower in maintenance, the trait that would ultimately slow each person's aging late in life. At ×0.4 per SD the change more than doubles, to −0.62 points. At the mortality recorded for these cohorts, childhood deaths nonetheless only lower dementia at 80–90 by about a quarter of a point, because relatively few children died and those who did differed only modestly from the survivors. The simulator is able to trace the entire cascade from the deaths in individuals to how it shapes the population to its impacts on dementia.

| Version | Prompt from RLB (excerpt) | The AI agent's interpretation and result |
|---|---|---|
| **v1**<br>What to model | "I would like to remake the [AgingBrain] simulator... Have a series of boxes that are the underlying world features, such as AD pathology prevalence and how it impacts brain and cognition... The goal here is that these defining boxes will be used to adjust the true trajectories of each person in the simulation, so that at every year they will have values for cognition, brain structures [and] pathology... Thus the boxes define what relations and risks exist in the world, and then the people are simulated that make up that world..." | Rebuilt the simulator around a rail of stages and a working World page. The boxes set each life's onset ages for amyloid, tau spread, LATE and metabolic syndrome, and every year's cognition, brain measures, pathology, dementia status and risk of death. |
| **v15**<br>Published anchor | "...Total intracranial volume can be referenced against this paper, noting the male versus female difference. There shouldn't be an age effect... Whole brain volume at age 20 would be nearly perfectly correlated with total intracranial volume, and [the hippocampus] should be a more or less fixed percentage of whole brain volume at that young age... Total intracranial [volume] does not atrophy with age." | Read the attached paper (Buckner et al., 2004) and took head-size norms from its Table 3: 1,413 mL for women and 1,579 mL for men, with a within-sex SD of 132 mL. Made the hippocampus 0.64% of whole brain at 20, with whole brain 0.84 × intracranial volume. |
| **v31–32**<br>Reality checks | "...Also add [an] expandable box above causal chains that is 'The properties of this simulated world'... Make an expandable pair of banners... 'How the model works'... The second... should be 'Reality checks', with reality checks like as shown, updated as is appropriate to [the] current version."<br>**Later:** "Yes, recalibrate." | Built the section, with the model's equations filled in with the current settings and 11 checks against published values, 5 of them in range. Offered to recalibrate toward the other 6 and, when asked, retuned the starting settings so all 11 fell in range, flagging one setting that sat at the high end of published estimates. |
| **v63**<br>Self-documentation | "Make a last box at the bottom on the left that is Imagine and uses a lightbulb... as its icon. This panel should bring up a tab that instructs an investigator how to load this HTML... and give example prompts that would replace that Imagine option with their own analysis... Embed into the HTML a self-documenting section... that has all of the descriptions from this chat as would be needed to provide the context to an entirely naive new chat from a new user to... build a new panel." | Added the Imagine stage, with example prompts and a developer guide inside the file. Then tested it: an agent that had never seen the conversation, given only the downloaded file and one example prompt, built a working new stage in Imagine's place, though it had to read about half the code for details the guide left out; the guide was then extended and retested. |
| **v65**<br>Published anchor | "Digest this image. This is a map of the pattern of how biomarkers amyloid, pTau, tau PET, MRI ([hippocampus]) and cognition move when affected by the AD cascade. The distance between the amyloid PET and tau PET is about 10 years. Note how some of the biomarkers saturate (e.g., hit a plateau)... Can you adjust the rates and constants so that these timings and patterns are generally held in our model?" | Retimed the AD cascade so that amyloid PET reaches half its rise about 10 years before tau PET. Across 24 points compared with the figure, a typical person's timing is now off by 0.9 years on average (4.9 before), and all 11 reality checks still pass. |
| **v75–76**<br>Model failure | "Can you articulate more specifically why an AD cascade cannot hit prevalence marks and still accommodate the very old getting AD?..."<br>**Then:** "What kind of process, if one was added, could yield the AD prevalence in the oldest old without it [being] simply aging?"<br>**Then:** "Yes, build it... Add a first paragraph on how resilience that falls with aging is needed and why, with its limitations and complexities." | Explained that raising AD risk lifts dementia at every age rather than steepening its rise. Having found that a faster cascade, more amyloid and steeper LATE all fell short, proposed resilience that falls with age, tested it in a scratch copy, and built it only when asked (v75). At RLB's request it became the first dated entry in Insights and puzzles (v76). |
| **v83**<br>Outside critique | "...Here is what a colleague said: '...the field at large do not believe that episodic memory or executive functions decline linearly from age 25 to late life...' Can you summarize and refer me to the papers that support his claim..."<br>**Then:** "Yes, draft a response, and let's modify simulation and paper to be consistent with the data in hand. Also make me a list of papers to gather for me to read fully and decide for myself." | Judged the colleague partly right: only two of the five cohorts he named can date when decline begins. Raised two modeling choices before building, then made normal episodic-memory and executive decline begin at 60, while processing speed still declines from 25. All 11 reality checks still pass. |
| **v89**<br>Correction | "Remove 'Amyloid anchored to the Mayo population' as an insight. It's not..."<br>**Then:** "...We already have a generation gain separate from school. See attached." | Removed the entry it had added on its own initiative. Conceded that a claim from its own audit, which it had repeated, was wrong: the model already had a generational gain of 0.13 SD per decade of birth beyond the 0.02 that schooling adds. |
| **v96_M&S**<br>New stage | "...The new stage, called 'What makes a maintainer?': In this stage allow a user to select either maintainer or SuperAger, using the default definitions, as used in the attached [figure], and illustrate the properties of the average 85-year-old in the group and 10 exemplars..." | In a new conversation given the file and the figure, added the stage just before Imagine and reproduced the figure. Listed where it departed from the prompt and the guide: it re-ran the simulation, against the guide's advice, because the main run keeps no values from 75 to 95, and it added the paper's pie chart, which was not requested. |
| **v99**<br>Side conversation | "...Please check the text and simulator against these papers and generally against one another to find errors."<br>**Then:** "What are comments that I can paste into the chat with the simulator to instruct it to make updates and corrections based on this review?"<br>**In the design conversation:** "Do these: 'Five cheap changes (2a, 3, 4a, 5a, 6a) work together and leave the checks where they are'..." | In the side conversation, read the 64 papers in 12 topic groups, replicated their designs in the simulator where possible, had a separate instance fact-check the report (about 30 corrections), and drafted prompts to paste into the design conversation. There, it tested each fix on a copy of the model and recommended five, which it applied once RLB chose them; 12 of 13 checks still pass. |
| **v100**<br>What to model | "Build me a new version of the simulation that has two separate AD cascades that each have on/off toggles... a second cascade that is 'AD cascade expanded', where tau absolute levels and also acceleration levels are associated with the downstream cascades... They should have settings that make them equivalent in the typical world; they would differ if an intervention lowered amyloid versus lowered tau..." | Built version 100 with the existing cascade untouched and on by default, and an expanded cascade that cannot run at the same time. Read "acceleration" as tau's rate of rise and said why. With the defaults, all 120,000 lives matched version 99 exactly. |
| **v101_MT**<br>Original code | "I received the response from the authors of the Josefsson rule, pasted here. Is this consistent with our implementation? I also paste their code to check." | Ran the authors' R code, unchanged, on the same simulated worlds: it gave the same slopes and picked the same 8,187 maintainers, person for person. Noted that their rule deliberately credits a high starting level, and asked before changing its report. |

**Supplementary Figure 8. Example consequential prompts and how the AI agent interpreted them.** Each row shows one simulator version that resulted from a novel prompt, with the request (left), excerpts of RLB's prompts (middle), and the agent's interpretation and result (right). Prompts are lightly edited for spelling, capitalization and punctuation; ellipses mark omitted text, and square brackets mark added or replaced words, including [AgingBrain] for the simulator's earlier names. Labels with a suffix (v96_M&S, v101_MT) are builds that added a stage outside the main line of versions (when the Imagine stage is used, the new simulator version adds a stage and renames the simulator with a suffix). Most exchanges took place in the main design conversation; v83 and the first two v99 prompts come from side conversations, and v96_M&S, v100 and v101_MT from other separate conversations. Across the rows, RLB states what to model, the assumptions with their published anchors and the reality checks, and corrects the AI agent; the AI agent turns each request into code, says where it departed from the request, and applies the calibrations and alternatives it proposes with feedback from the user.

### Supplementary References


Ashton, N.J., Brum, W.S., Di Molfetta, G., Benedet, A.L., Arslan, B., Jonaitis, E., et al. (2024). Diagnostic accuracy of a plasma phosphorylated tau 217 immunoassay for Alzheimer disease pathology. JAMA Neurol. 81, 255–263. https://doi.org/10.1001/jamaneurol.2023.5319

Bell, F.C., and Miller, M.L. (2005). Life Tables for the United States Social Security Area 1900–2100. Actuarial Study No. 120 (Social Security Administration, Office of the Chief Actuary). https://www.ssa.gov/oact/NOTES/as120/TOC.html

Bollack, A., Collij, L.E., Vállez García, D., Shekari, M., Altomare, D., Payoux, P., et al. (2024). Investigating reliable amyloid accumulation in Centiloids: results from the AMYPAD Prognostic and Natural History Study. Alzheimers Dement. 20, 3429–3441. https://doi.org/10.1002/alz.13761

Buckner, R.L., Head, D., Parker, J., Fotenos, A.F., Marcus, D., Morris, J.C., et al. (2004). A unified approach for morphometric and functional data analysis in young, old, and demented adults using automated atlas-based head size normalization: reliability and validation against manual measurement of total intracranial volume. Neuroimage 23, 724–738. https://doi.org/10.1016/j.neuroimage.2004.06.018

Castellano, T., Wang, T.C., Nolan, E., Wu, Y., Zhang, M., Clifton, M., et al. (2025). APOE, ABCA7, and RASGEF1C are associated with earlier onset of amyloid deposition from more than 4000 harmonized positron emission tomography images. Alzheimers Dement. 21, e71006. https://doi.org/10.1002/alz.71006

Children's Bureau (1945). Childhood Mortality from Accidents: By Age, Race, and Sex and by Type of Accident. Publication No. 311, prepared by G. Wolff (U.S. Government Printing Office). https://fraser.stlouisfed.org/title/8579

Fjell, A.M., Grødem, E.O.S., Lunansky, G., Vidal-Piñeiro, D., Rogeberg, O.J., and Walhovd, K.B. (2026). Generational gains in memory capacity and stability may account for declining dementia incidence rates in Europe and the United States. Preprint at medRxiv. https://doi.org/10.64898/2026.04.14.26350835

Gantenberg, J.R., La Joie, R., Heston, M.B., and Ackley, S.F.; Alzheimer's Disease Neuroimaging Initiative (2026). Do amyloid trajectories reach a ceiling? Evidence from iterative approximation and simulation. Alzheimers Dement. 22, e71781. https://doi.org/10.1002/alz.71781

Hansson, O. (2021). Biomarkers for neurodegenerative diseases. Nat. Med. 27, 954–963. https://doi.org/10.1038/s41591-021-01382-x

Insel, P.S., Mattsson-Carlgren, N., Langford, O., Caruso, V.M., Leuzy, A., Young, C.B., et al. (2025). Concurrent changes in plasma phosphorylated tau 217, tau PET, and cognition in preclinical Alzheimer disease. JAMA Neurol. 82, 985–993. https://doi.org/10.1001/jamaneurol.2025.2974

Jack, C.R., Jr., Knopman, D.S., Jagust, W.J., Petersen, R.C., Weiner, M.W., Aisen, P.S., et al. (2013). Tracking pathophysiological processes in Alzheimer's disease: an updated hypothetical model of dynamic biomarkers. Lancet Neurol. 12, 207–216. https://doi.org/10.1016/S1474-4422(12)70291-0

National Center for Health Statistics (1960–2019). Deaths by age and cause, United States (Health, United States; America's Children; leading causes of death by age). https://www.cdc.gov/nchs/hus/index.htm

Ossenkoppele, R., Coomans, E.M., Apostolova, L.G., Baker, S.L., Barthel, H., Beach, T.G., et al. (2025). Tau PET positivity in individuals with and without cognitive impairment varies with age, amyloid-β status, APOE genotype and sex. Nat. Neurosci. 28, 1610–1621. https://doi.org/10.1038/s41593-025-02000-6

Roberts, R.O., Aakre, J.A., Kremers, W.K., Vassilaki, M., Knopman, D.S., Mielke, M.M., et al. (2018). Prevalence and outcomes of amyloid positivity among persons without dementia in a longitudinal, population-based setting. JAMA Neurol. 75, 970–979. https://doi.org/10.1001/jamaneurol.2018.0629

US Census Bureau (2026). Table A-1. Years of school completed by people 25 years and over, by age and sex: selected years 1940 to 2025. CPS Historical Time Series Tables. https://www.census.gov/data/tables/time-series/demo/educational-attainment/cps-historical-time-series.html (accessed November 17, 2026).
